# Environmental modulators of algae-bacteria interactions at scale

**DOI:** 10.1101/2023.03.23.534036

**Authors:** Chandana Gopalakrishnappa, Zeqian Li, Seppe Kuehn

**Affiliations:** Department of Physics. The University of Illinois at Urbana-Champaign. Urbana, IL 61801; Center for the Physics of Evolving Systems. The University of Chicago. Chicago, IL 60637; Department of Ecology and Evolution. The University of Chicago. Chicago, IL 60637

## Abstract

Photosynthetic microbes associated with non-photosynthetic, heterotrophic, bacteria play a key role in the global primary production. Understanding these phototroph-heterotroph associations is therefore important, but remains challenging because they reside in chemically complex aquatic and terrestrial environments. We do not understand how the myriad of environmental parameters from nutrient availability to pH impact interactions between phototrophs and their heterotrophic partners. Here, we leverage a massively parallel droplet microfluidic platform that enables us to interrogate algae-bacteria interactions in *>*100,000 communities across ∼525 environmental conditions with varying pH, carbon availability and phosphorous availability. By developing a statistical framework to dissect interactions in this complex dataset, we reveal that dependance of algae-bacteria interactions on nutrient availability is strongly modulated by pH and buffering capacity. Furthermore, we show that the chemical identity of the available organic carbon source controls how pH, buffering capacity, and nutrient availability modulate algae-bacteria interactions. By leveraging a high-throughput platform, our study reveals the previously underappreciated role of pH in modulating phototroph-heterotroph interactions.

## INTRODUCTION

Microbial communities occupy nearly every niche on Earth, from animal hosts, to soils, and oceans. These complex consortia often contain many interactions between members whereby one species impacts the abundances of another. Interactions in these communities can determine the outcome of invasions [1], metabolic processes such as carbon and nitrogen remineralization [2], or the phenotype of the host [3]. Crucially, however, interactions between members of a microbial consortium depend on the environmental context. For example, changes in pH, nutrient availability, temperature, or toxic metabolic byproducts, can strongly modulate interactions between members of a collective [4–7]. As a result, an important question in ecology is understanding how environmental parameters impact interactions.

Understanding how environmental parameters influence ecological interactions in consortia faces two related challenges. First, the physicochemical environment in natural microbial communities is high-dimensional in the sense that there are many possible parameters that change in time and space and can impact the outcome of an interaction [8]. This high-dimensionality means that experimentally interrogating how interactions depend on the environment is a daunting task. For example, if we wanted to measure how four different environmental variables (say, pH, carbon, nitrogen, phosphorous availability) impacted an interaction this would require 10.000 experiments if we included just 10 levels for each environmental variable. To determine interactions between just two taxa would require measuring their growth alone and in pair-culture in each one of these conditions – meaning 30.000 measurements would be required, a huge undertaking.

The second problem is conceptual. Is it necessary to understand how interactions depend on each and every environmental parameter or are there simple, global, patterns across environmental conditions that can be discerned? In essence, are interactions idiosyncratic from one environmental condition to the next, meaning that to elucidate the impact of the environment on an interaction we must study many distinct environmental conditions? Or, are there patterns that exist across conditions that make it easier to understand the impact of environmental variables on interactions? For example, one possibility is that environmental variables impact interactions in hierarchical fashion where some high-level parameters are very important in determining the outcome of an interaction and others are successively less critical.

Here we address these two questions using a massively parallelized droplet microfluidic platform [9] to interrogate interactions between a photosynthetic alga (phototroph) and a heterotrophic bacterium. Phototroph-heterotroph interactions form the basis of biomass production in ecosystems on a global scale, from marine to freshwater ecosystems to soils and industrial photobioreactors [10, 11]. In this capacity, phototroph-heterotroph interactions form a critical link in the global cycle of carbon by converting inorganic carbon to biomass. Moreover, interactions between phototrophs and heterotrophs are mediated by a myriad of environmental factors from carbon, nitrogen, and phosphorous availability to temperature, light, and small molecule exchanges [12–19]. In addition, the outcome of phototroph-heterotroph interactions are important for understanding eutrophication and dead zones [20]. Finally, these communities hold key biotechnological importance in the context of biofuel production, or the production of industrially important precursors [11].

We interrogate phototroph-heterotroph interactions across hundreds of environmental conditions using a microfluidic platform that leverages nanoliter droplets, with contents barcoded using fluorescent dyes, to measure abundance dynamics in *>*20.000 cultures in a single experiment. Using this approach, we measure the interaction between the model alga *Chlamydomonas reinhardtii* and the bacterium *Escherichia coli* in ∼525 environmental conditions in *>*10 replicates each for both monoculture and pair-culture.

Within the droplets, we measure algae-bacteria abundance dynamics via microscopy across a range of carbon sources and concentrations, phosphorus concentrations, pH, and buffering capacities. The resulting dataset proves amenable to statistical analysis where a regression reveals the key environmental drivers of algae-bacteria interactions. While previous studies suggest that nutrient availability is the key driver of interactions between phototrophs and heterotrophs, we find that pH and buffering capacity qualitatively alter how the availability of nutrients impacts interaction between algae and bacteria. Thus, we show that across a huge range of environmental conditions, pH and the ability of the environment to resist changes in pH (buffering capacity), act as important regulators of the interaction between phototrophs and heterotrophs. Finally, the role of the environmental factors -pH, buffering capacity, and nutrient availability in regulating interactions is modified by the chemical identity of an exogenously available organic carbon. These results suggest that chemical composition of organic carbon and pH interact to qualitatively determine the outcome of algae-bacteria interactions.

## RESULTS

### The model system and environmental conditions

The microbial community under study comprises the alga, *Chlamydomonas reinhardtii*, commonly found in soils and freshwater [21], as the phototroph, and the soil-dwelling bacterium [22], *Escherichia coli*, as the heterotroph. We note that these microbes are not known to coexist in the wild and so we expect no strong co-evolutionary history between these organisms. Despite this, these two microbes have been widely used in studies as a model phototroph and heterotroph due to their thorough biological characterization, ease of cultivation, and accessibility to molecular techniques and quantitative measurements. Previous studies of closed microbial communities including these two microbes, in addition to a ciliate, have revealed strongly deterministic dynamics on timescales of months and rich spatiotemporal and phenotypic processes [23, 24]. Another study demonstrated the presence of higher-order interactions between this alga and bacteria mediated by a ciliate. [1]. Thus, the interactions between these two model organisms constitute a tractable test bed for understanding phototroph-heterotroph interactions.

In this study, interactions between the algae, *C. reinhardtii*, and the bacteria, *E. coli* were assayed in the modified 1/2x Taub media (a freshwater mimic media) that varied in five environmental factors - initial pH, buffering capacity, phosphorus concentration, carbon concentration, and carbon source identity. Although resource competition and exchange are identified as key players in driving phototroph-heterotroph interactions [10, 11, 13– 16, 19, 25], several studies have reported a strong correlation between microbial communities compositions and environmental factors such as pH and concentration of nutrients -carbon, nitrogen and phosphorus [26, 27]. Additionally, it is well known that the identity of the carbon source affects *E. coli* metabolism via impacting the growth rate and the nature of the metabolic products, which could potentially lead to different interactions with *C. reinhardtii* [28–30]. Therefore, we reasoned that a multitude of abiotic factors such as pH, buffering capacity, light level, including nutrient concentration may contribute to phototroph-heterotroph interactions (Fig. 1). Hence, we choose the above 5 factors. The values of each of the environmental factors were chosen to be in the biological range: 6.1-7.5 for initial pH, ∼0-3.5 mM for buffering capacity, 0.01 mM - 4 mM for phosphorus concentration, 2 mM - 10 mM for the carbon concentration (Section 3.2, SI). And the different carbon sources considered for the study were glycerol, glucose, galactose, pyruvate, and acetate. For each of these carbon sources, the algae-bacteria interactions were assayed in a total of ∼105 environmental conditions varying in initial pH, buffering capacity, and phosphorus and carbon concentration, using the high throughput platform discussed in the following section.

**Figure 1.**
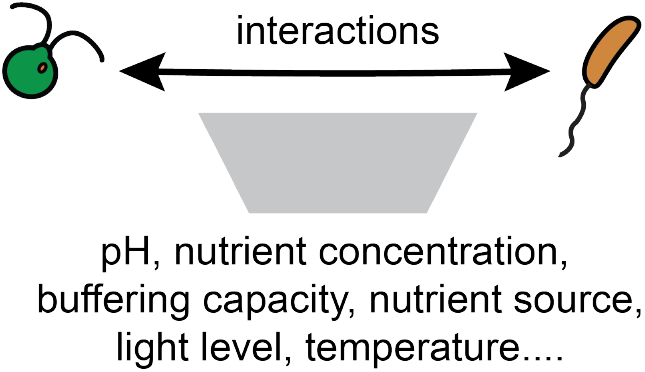
Dependence of algae-bacteria interactions on environmental factors is complex. Cartoon illustration of our hypothesis that diverse interactions between algae and bacteria are mediated by a multitude of chemical factors in the environments such as concentration of nutrients, pH, buffering capacity, light level, and temperature.

### High-dimensional characterization of phototroph-heterotroph interactions

In this study, we used droplet-based microfluidic chip (“kChip” with k=2) to rapidly assay the phototroph-heterotroph interactions in hundreds of environmental conditions in parallel. The kChip platform has previously been utilized for drug discovery, pathogen detection, and the study of bacterial interactions [9, 31–33]. Briefly, the experiment proceeds by first generating a library of environmental conditions that vary in the initial pH, buffering capacity, concentration of phosphorus, and concentration of carbon of a chemically defined minimal medium (Fig. 2A; Section 3.2, SI). Initial pH refers to the starting pH of the environment, which we varied by using buffers and titration. To vary the buffering capacity of the environment, we added different concentrations of organic buffers (Tris or MOPS). We fluorescently barcoded each environmental condition using three fluorescent dyes in low concentrations and added algae and bacteria independently. Using these precultures, a commercial droplet generator was used to create thousands of nanoliter water-in-oil droplets containing algae or bacteria in each of the predefined nutrient conditions. These droplets were then pooled and loaded into a kChip microfluidic chip platform which contains ∼25,000 microwells, each of which randomly groups two droplets containing microbes in predefined media conditions, resulting in the formation of all possible combinations of communities (monocultures and co-cultures) and environmental conditions (Fig. 2A; Section 5, SI). The chip is then imaged to identify the fluorescent dye barcodes and thereby infer the environmental conditions present in each microwell (Section 6.2, SI). Subsequently, the droplets in each microwell were merged via exposure to an alternating electric field, leading to the formation of the phototroph-heterotroph communities in hundreds of environmental conditions. Thereafter, the kChip was incubated at 30 °C under light (68.5 *µ*mol *m*^2^*s*^−1^) to allow for growth. The chip was then imaged at regular intervals (approximately 0 h, 12 h, 21 h, 45 h, 68 h) to track the growth of the microbes using chlorophyll fluorescence for *C. reinhardtii* and genetically encoded GFP fluorescence for *E. coli* (Fig. 2B; Section 6.3, SI). Algal and bacterial abundances over time were determined by analyzing the microscopy images, generating microbial growth curves, and estimating growth (difference between the initial abundances and the final abundances at the end of the experiment) for both the phototroph and heterotroph in the ∼1395 microbial communities constructed in the kChip experiments for all the carbon sources (Fig 2C; Section 5, SI). We note that the abundances of the microbes do not saturate with time in several of the environmental conditions during the duration of our experiments.

**Figure 2.**
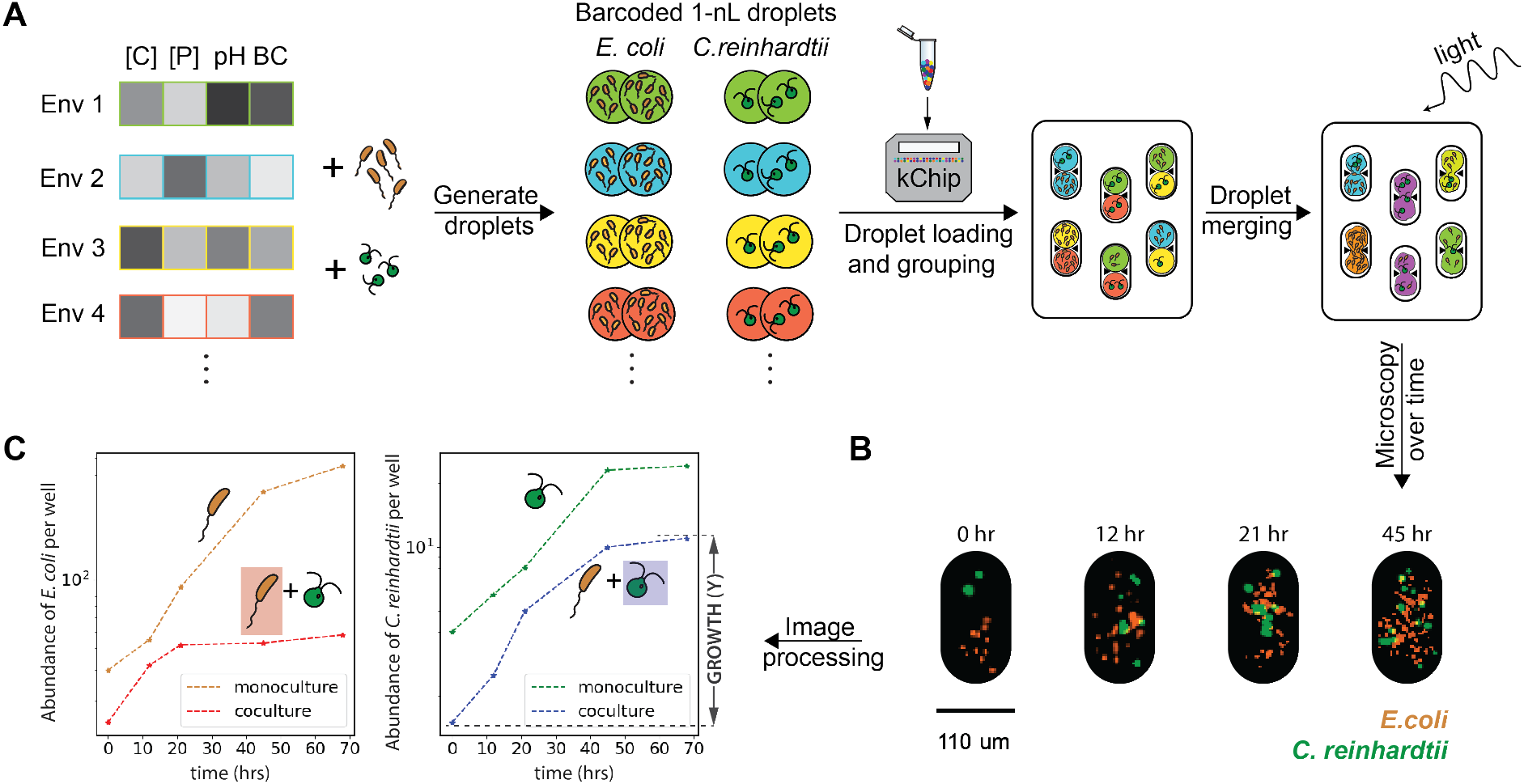
A high-throughput droplet platform for measuring algae-bacteria growth in hundreds of environments. (A) Setting up the microfluidic chip. Environments (media conditions) varying in the factors - Initial pH, buffering capacity, phosphorus concentration, and carbon concentration, are prepared and barcoded using three fluorescent dyes (Section 3.2, SI). After adding the bacteria (brown) and algae (green) independently to each barcoded media, nanoliter droplets of each of the microbes in the barcoded environments are generated. The generated droplets are pooled together and loaded on the microfluidic chip which randomly groups two droplets in each of its microwells. The chip is then imaged for fluorescent barcodes using a widefield fluorescence microscope, to infer the values of the environmental factors in the microwells via image processing (Section 6.2, SI). Following exposure of the chip to an alternating electric field, droplets in the microwells merge to form replicates of bacterial monocultures, algal monocultures, and algae-bacteria cocultures in all combinations of the environments that were present in the initial droplets. The chip is then incubated at 30 °C under light (68.5 *µ*mol *m*^2^*s*^−1^). (B) Microscopy images of a single microwell showing the growth of algae and bacteria over time. The GFP fluorescence image representing the bacteria (in brown) and the chlorophyll fluorescence image representing the algae (in green) are overlayed in these images. The first image shows the bacteria and the algae in the separate compartments of the well, prior to the merging of the droplets. The later images show the increase in the abundance of the algae and bacteria at 12 h, 21 h, and 45 h. (C) Example growth curves of algae and bacteria in monoculture and coculture in an environmental condition. The images of the chip are analysed to infer the abundances of the microbes in the microwells over time (Section 6.3, SI). The growth Y of algae and bacteria are then obtained by estimating the increase in their respective abundances at 68 h from their abundances at 0 h (black arrow labeled ”GROWTH (Y)” right panel).

Previous studies utilizing this platform studied bacteria. So, we modified existing protocols to make the measurement compatible with algae. Specifically, we added the functionality for imaging chlorophyll fluorescence to track the growth of *C. reinhardtii* and devised a computational pipeline to remove the bleed-through between chlorophyll fluorescence and one of the barcoding dyes (Section 6.1, SI). This expanded the number of fluorophores that can be probed on the kChip from four to five.

### Algae inhibit bacterial growth and bacteria weakly affect the algal growth

To begin, we compared the growth of both algae and bacteria in cocultures to their growth in monocultures. The bacterial growth in cocultures were lower than their respective growth in monocultures in all the environmental conditions, suggesting inhibition of *E. coli* by *C. reinhardtii* (Fig. 3A; Fig. S8 top panels, SI). Additionally, the *E. coli* cells show greater aggregation in monocultures than in cocultures (Fig. S6, SI). These results are consistent with a previous study that showed that introducing bacteria into algal cultures results in the inhibition of bacterial growth and the dispersal of bacterial aggregates [1].

**Figure 3.**
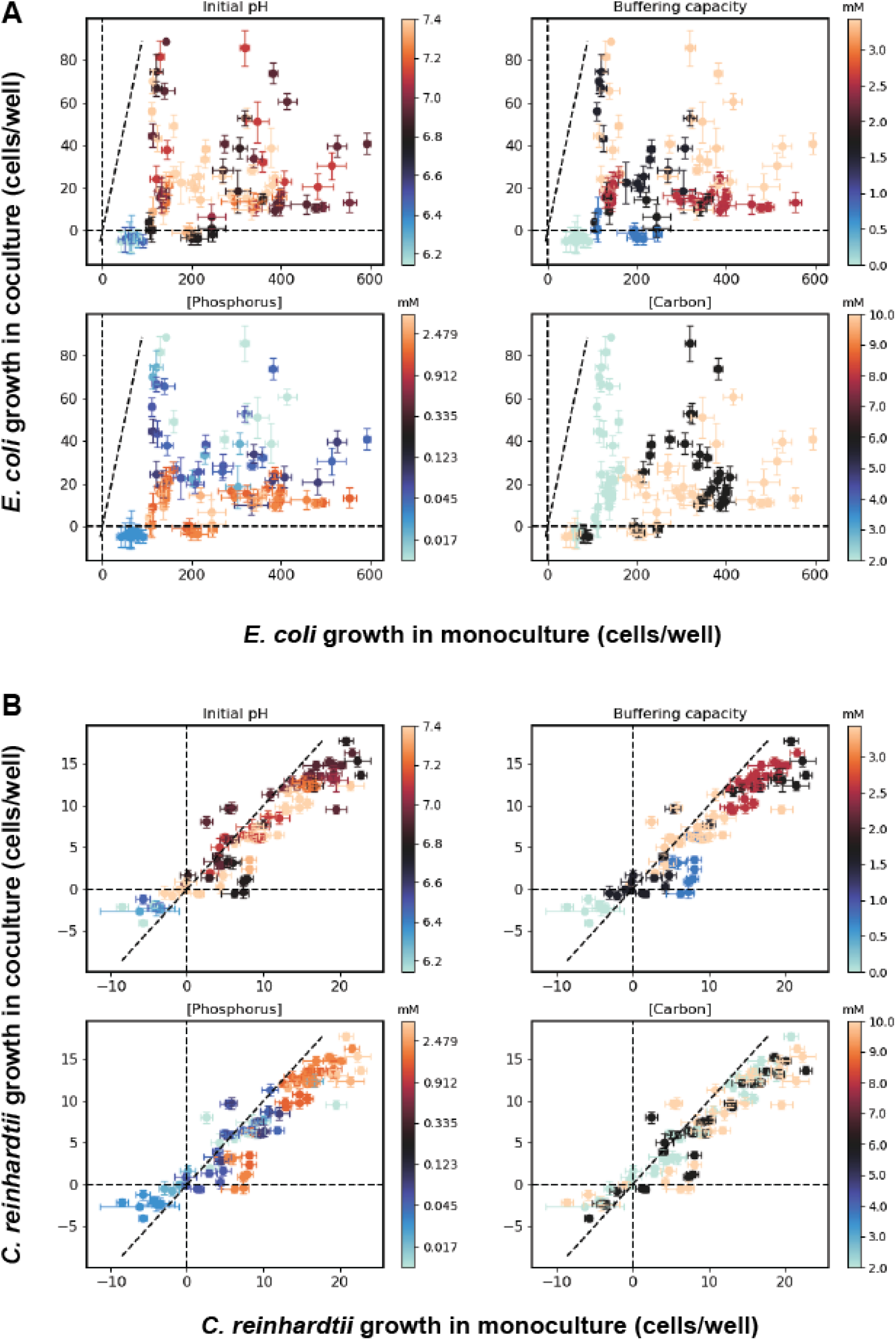
Complex dependence of algae-bacteria interactions on the environmental factors. (A) Panels show bacterial growth in monoculture (x-axis) and co-culture (y-axis). Each point indicates median growth (Fig. 2C) of *E. coli* in co-culture and monoculture computed across replicates of each environmental condition. Error bars indicate the standard error of the mean growth. The dashed line indicates equal growth in monoculture and co-culture. Note the fact that all points lie below this line indicating the pervasive inhibition of bacteria by algae. The data on each panel are the same, but the colormaps represent each of the four environmental factors - Initial pH (top left), buffering capacity (top right), phosphorus concentration (bottom left), and carbon concentration (bottom right). The colormap for phosphorus is logarithmic. The carbon source is glycerol. See Fig. S8 for the other carbon sources’ data. (B) Identical plots as in (A) but for algal growth in monoculture and co-culture. The fact that most data lie near the dashed line indicate overall weaker impacts on algal growth by bacteria.

*C. reinhardtii*, on the other hand, has similar growth in cocultures and monocultures in most cases, indicating a weak effect of *E. coli* on the growth of *C. reinhardtii* (Fig. 3B; Fig. S8 bottom panels, SI). There do exist a few environments where *C. reinhardtii* is suppressed or enhanced in co-culture relative to monoculture, indicating an impact of the presence of the bacteria.

### Algae-bacteria interactions show complex dependence on the environmental factors

Next, we sought to understand the dependence of algae-bacteria interactions on environmental factors. To visualize this, we plotted the growth in cocultures against the growth in monocultures, color-coding the data for each of the four environmental variables considered - Initial pH, buffering capacity, concentration of carbon and phosphorus (Fig. 3; Fig. S8 top panels, SI). These plots show no distinct grouping of the data based on any of the four environmental factors and indicate a complex dependence of algae-bacteria interactions on the environmental factors. For example, in the case of *E. coli*, while the low carbon concentration (the light green points in Fig. 3A bottom right panel) sets the growth in monocultures to low values, the variation in other environmental factors (pH, buffering capacity) causes the coculture growth to span from low to high values. There also exist cases where a single factor determines the effect of the environment on monoculture and coculture growth. For example, low buffering capacity, not initial pH or nutrients’ concentration, appears to give rise to death of *C. reinhardtii* (light green points have growth less than zero) (Fig. 3B top right panel).

When we compute correlations between the environmental factors and growth, we see significant statistical relationships between each parameter and the bacterial or algal growth (Fig. S7, SI) across carbon sources. These correlations reinforce the idea that there is a complex interplay between nutrients’ concentration, pH, buffering capacity and identity of the carbon source in determining algae-bacteria interactions.

The interesting aspect of the above result is that initial pH and buffering capacity are shown to affect algae-bacteria interactions. This result agrees with surveys of communities in the wild which show that pH is an important environmental factor in determining community structure [26, 27]. In contrast, most previous experimental interrogations of interactions between phototrophs and heterotrophs focus on the role of nutrient concentration and competition [12, 14–16, 34]. We expect that pH and buffering capacity are likely affecting interactions by influencing physiology including nutrient uptake rates.

Next we sought a framework to quantify interaction between algae and bacteria in our experiment. We considered consumer-resource models to quantify competition for carbon, nitrogen, and phosphorus. However, the interactions in our community cannot be described by a model that considers only these nutrients. For example, the overall inhibition of *E. coli* does not depend in a simple way on the concentration of nutrients. Similarly, variations in pH are not naturally modeled in a consumer-resource framework. Hence, a simple consumer-resource model approach is not suitable for dissecting the interactions in our data. We, therefore, took a statistical approach using simple linear regressions to model interactions as a function of the environmental factors.

### Quantifying algae-bacteria interactions statistically

Our goal is to quantify how the presence of algae or bacteria impacts the growth of the other species across all the environmental conditions tested. To do this, we developed a simple framework for estimating interactions in the algae-bacteria communities via regression analyses. Specifically, we used a linear regression formalism to predict algal or bacterial growth (Fig. 2C) using environmental factors (pH, buffering capacity, phosphorus concentration and carbon concentration) as independent variables. We performed independent regressions to predict algal and bacterial growth across all conditions.

Our regression approach can be explained mathematically using a simple example. Imagine communities of algae and bacteria where the growth are affected by a single environmental factor *X* and by the presence of the other species via an interaction. In this case, the model for predicting the growth of *E. coli* in monoculture and coculture would take the following form:

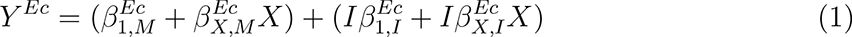

where *Y ^Ec^* is the growth of *E. coli* and the 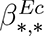 are regression coefficients. *I* is a variable that indicates the presence of *C. reinhardtii* (*I* = 0 in monoculture, *I* = 1 in coculture). The coefficient 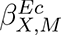 represents the change in growth in monoculture per unit change in *X* and 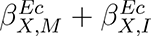 represents the change in growth in coculture per unit change in *X* (Fig. 4A). Hence, 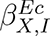, estimates the average change in growth per unit *X* in coculture relative to monoculture. In other words, 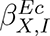 represents the effect of *C. reinhardtii* on *E. coli* as *X* increases, in coculture. A positive coefficient would represent enhancement of *E. coli* growth by *C. reinhardtii* as *X* increases (Fig. 4B left panels). Similarly, a negative coefficient would represent suppression of *E. coli* growth by *C. reinhardtii* as *X* increases (Fig. 4B right panels). An identical regression is used to estimate the impact of *E. coli* on *C. reinhardtii* growth.

**Figure 4.**
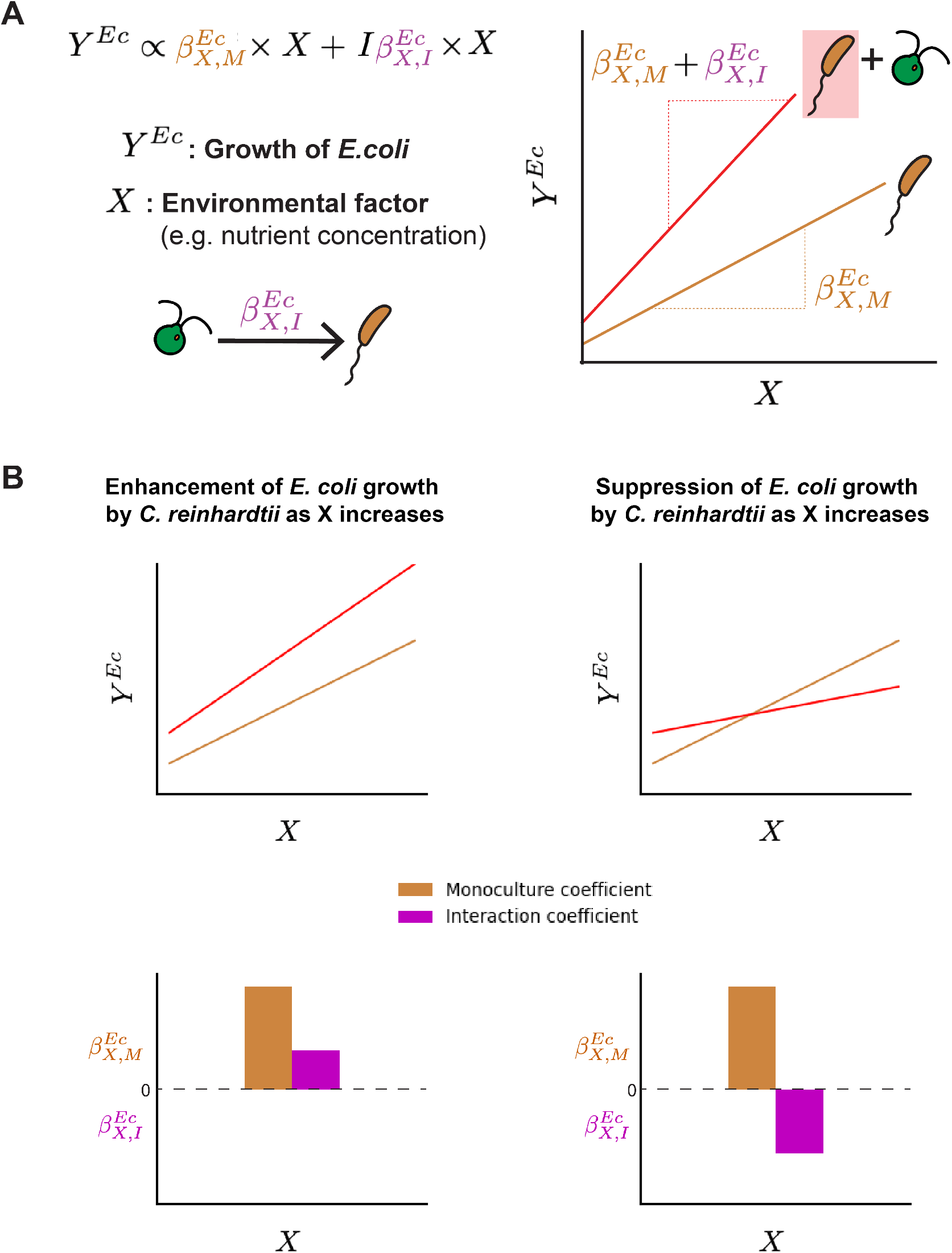
Quantifying algae-bacteria interactions using regression analyses. (A) Formulation of the regression model for predicting growth from environmental conditions, here using *E. coli* as an example. *Y ^Ec^*is the growth of *E. coli* in monocultures and cocultures and *X* is an environmental factor that determines the growth. The indicator variable *I* is set to 0 for growth in monoculture and 1 for growth in co-culture. The coefficient 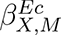 represents the change in growth in monoculture with *X* and is referred to as a monoculture coefficient. The coefficient 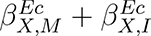 represents the changes in growth in coculture with *X* (shown schematically in the plot on the right). Hence, the coefficient 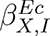 represents the change in the effect of *X* on growth in coculture relative to monoculture. The coefficient 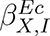 is referred to as an interaction coefficient. (B) Illustration of enhancement and suppression of *E. coli* growth by *C. reinhardtii* as *X* increases. The growth of *E. coli* in monoculture (in brown) and coculture (in red) vs the environmental factor *X* plotted in the case of enhancement (top left panel) and suppression (top right panel) of *E. coli* growth by *C. reinhardtii* as *X* increases. The panels on the bottom row show the corresponding regression coefficients. The monoculture coefficient 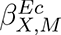 (in brown) and interaction coefficient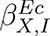 (in magenta) in the case of enhancement (bottom left panel) and suppression (bottom right panel) of *E. coli* growth by *C. reinhardtii* as *X* increases.

We extended the above model to include the effect of multiple environmental factors in determining growth of both species (Section 8, SI). For our dataset comprising of four environmental factors - initial pH (*pH*), buffering capacity (*BC*), phosphorus concentration ([*P*]), and carbon concentration ([*C*]), the model includes the following terms: [*P*], [*C*]*, pH*[*P*]*, pH*[*C*]*, BC*[*P*]*, BC*[*C*], [*P*][*C*]. For each term, we estimated a coefficient for monoculture and interaction as described above. For simplicity, we refer to coefficients of features without the indicator variable *I* as monoculture coefficients and coefficients of features with the indicator variable as interaction coefficients.

We did not include linear terms in *pH* or *BC* in our model because biologically *pH* alone does not generate biomass, but instead modulates the ability of cells to grow on the available nutrients. Thus, we included only interaction effects between nutrients and *pH* or *BC*. Therefore, the coefficient 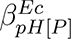 represents the susceptibility of growth to phosphorus concentration modulated by pH. The feature [*P*][*C*] was included to capture interactions between nutrients. Additionally, our model being simple, cannot capture nonlinearities in the growth as a function of a nutrient concentration. Despite these limitations, this statistical approach allows us to achieve a unified and interpretable picture of interactions between these microbes across a wide range of environmental conditions.

Finally, to account for the fact that algae globally inhibit bacterial growth in our experiment, we standardize the growth of both *E. coli* and *C. reinhardtii* prior to performing the regression above (Section 8.2, SI). Thus, our regressions describe variation in bacterial growth after removing the effect of this global inhibition. To facilitate interpretation, we also standardized all the independent variables in the regression. As a result, the regression coefficients describe the relative change in growth per unit change in each environmental factor and do not quantify the broad inhibition of bacteria by the alga. This standardization also allows us to compare coefficient values for regressions performed on different carbon sources despite variation in the growth on those nutrients. To perform the regression, we fit the growth measured in each well using a weighted least-square approach (Section 8.3, SI).

In general, we find that this model provides good predictions of growth across environmental conditions in our experiment, the fits being better for some carbon sources (glucose, glycerol, acetate) than others (galactose) (Fig. S9, SI). We note that a more complex model, such as a decision tree regression, gives superb fits to the data at the expense of interpretability (Fig. S12, SI).

### pH and buffering capacity modulate nutrient dependence of algae-bacteria interactions

Using the linear regression approach outlined above, we modeled the dependence of algal and bacterial growth on the environmental factors for each of the five carbon sources in monoculture and coculture. We first looked at the regression coefficients describing the growth of *E. coli* in one particular carbon source (glycerol, Fig. 5A). Of all the monoculture coefficients (brown bars in top panel Fig. 5A) obtained from fitting *E. coli* growth in glycerol, the coefficient of *BC*[*C*] is the largest, suggesting a strong interaction of buffering capacity with carbon concentration in determining the monoculture growth. Thus when *BC* is high, there is significantly more growth per unit [*C*] than when *BC* is low. These results are consistent with the greater acidification of the environment at lower buffering capacity observed in the microtiter plate experiments (Section 10, SI); this greater acidification likely negatively impacts *E. coli*. Therefore, the *E. coli* growth is expected to be higher at a higher buffering capacity for the same carbon concentration, which is reflected in the high value of the *BC*[*C*] coefficient. In addition to *BC*[*C*], there also exist statistically significant interactions between pH and carbon concentration, and buffering capacity and phosphorus concentration, with the magnitude of the coefficients of *pH*[*C*] and *BC*[*P*] being comparable or greater than the coefficients of [*P*] and [*C*] alone. Mechanistically interpreting each of these coefficients is beyond the scope of the present work, but could be pursued via additional experiments in the droplet platform or lower throughput batch cultures.

**Figure 5.**
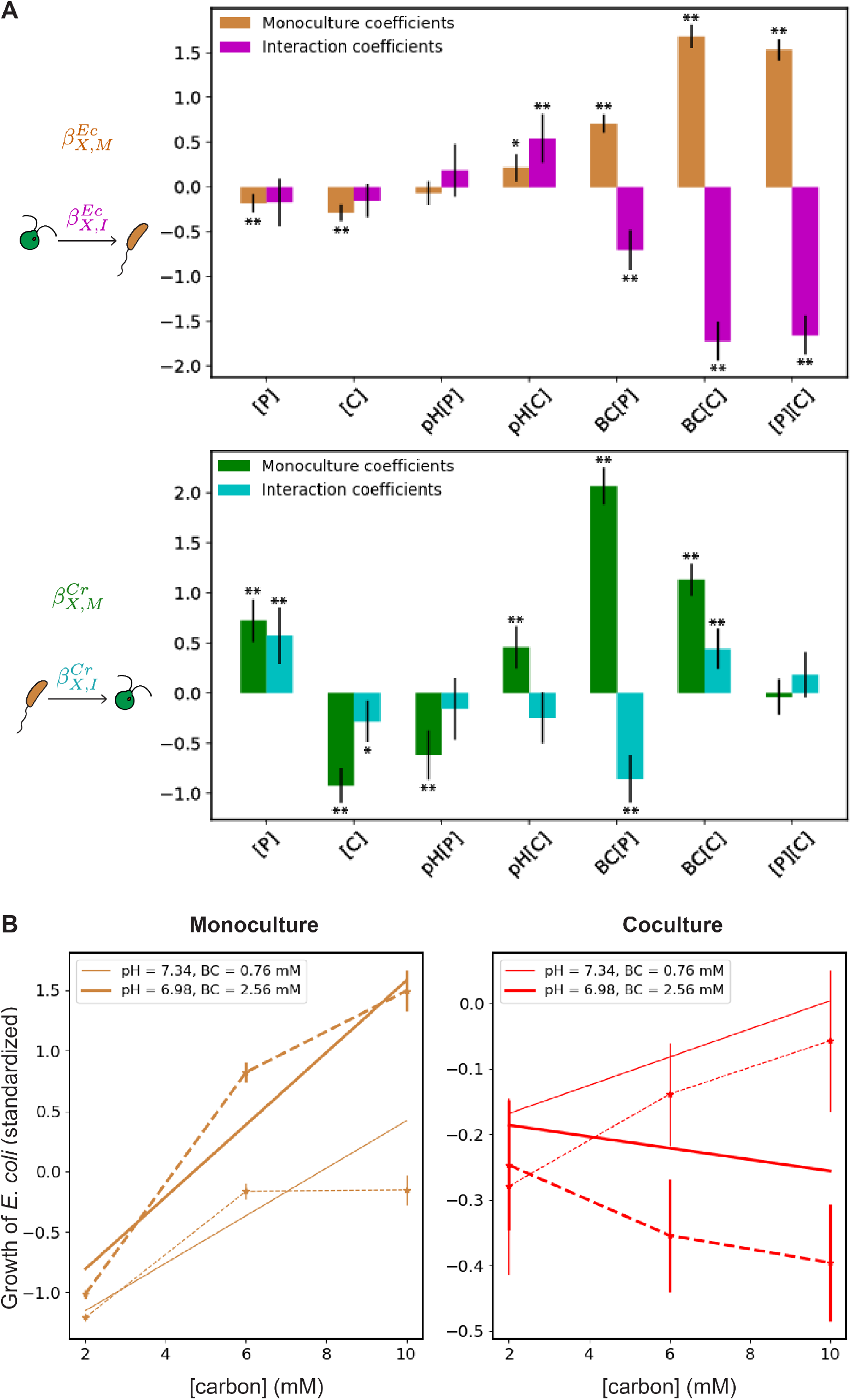
pH and buffering capacity modulate nutrient dependence of algae-bacteria interactions. (A) The coefficients for regressions predicting algal and bacterial growth in coculture and monoculture in glycerol. The results for the other carbon sources are shown in Fig. S10. The top panel reports the monoculture coefficients 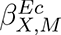 (brown bars) and the interaction coefficients 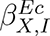 (magenta bars) of the corresponding features on the x-axis obtained for the regression model predicting the growth of *E. coli* in monocultures and cocultures. The interaction coefficients (magenta bars) indicate the effects of *C. reinhardtii* on *E. coli* growth with an increase in the corresponding features in coculture. The bottom panel reports the monoculture coefficients 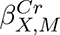 (green bars) and the interaction coefficients 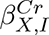 (cyan bars) of the corresponding features on the x-axis obtained from the regression model predicting the growth of *C. reinhardtii* in monocultures and cocultures. The interaction coefficients (cyan bars) indicate the effects of *E. coli* on *C. reinhardtii* growth with an increase in the corresponding features in coculture. The error bars represent the 95% confidence intervals. ** indicates a p-value *<*0.001 and * a p-value *<*0.05. (B) Example data illustrating modulation of the effect of carbon concentration on the growth of *E. coli* by pH and buffering capacity. The median bacterial growth in monoculture and coculture are plotted as a function of carbon concentration at [P]∼1.51 mM in the left and right panels respectively. The experimental data are represented by circles and connected with dashed lines. The error bars represent the standard error about the mean bacterial growth. The solid lines represent the model prediction. Darker or thicker lines represent the results at low pH (6.98) and high buffering capacity (2.56 mM) and lighter or thinner lines represent the results at high pH (7.34) and low buffering capacity (0.76 mM).

Next, among the interaction coefficients containing the factors *pH* and *BC* (magenta bars in Fig. 5A top panel), the coefficients of *pH*[*C*], *BC*[*P*] and *BC*[*C*] are non-zero and compare in magnitude with their respective monoculture coefficients. This reveals that the effects of *pH*[*C*], *BC*[*P*], and *BC*[*C*], on bacterial growth in coculture are significantly different compared to their effects in monoculture. We conclude from this that the interaction between *C. reinhardtii* on *E. coli* is strongly mediated by the factors - pH and buffering capacity. This is a central finding of our study.

The fact that pH and buffering capacity of the environment can strongly influence interactions is illustrated by looking at a specific example from the data (Fig. 5B). Choosing a subset of data corresponding to a specific phosphorus concentration ([*P*] ∼1.51 mM), we compared the change in growth with carbon concentration in monocultures and cocultures at the different pH and buffering capacities values. The change in *E. coli* growth in monoculture with carbon concentration at the different buffering capacities shows different behavior (Fig. 5B, left panel). Particularly, the increase in the growth with carbon concentration is observed to be higher in the condition with high buffering capacity (and low pH) compared to the increase in the condition with low buffering capacity (and high pH) as expected, with the trends in the model and the data being in good agreement. Next, we compare these results to *E. coli* growth in coculture. The trends in *E. coli* growth with carbon concentration in coculture are distinct from monoculture and depend on the pH and buffering capacity values (Fig. 5B right panel). The growth appreciably declines with carbon concentration in the condition with low pH (and high buffering capacity) whereas there is an increase in growth with carbon concentration at high pH (and low buffering capacity), with the model reasonably capturing the trend in the data. These results agree with the positive coefficient of *pH*[*C*] and ∼0 coefficient of *BC*[*C*] obtained when the model is evaluated for *E. coli* growth in coculture (sum of brown and magenta *pH*[*C*] and *BC*[*C*] bars in Fig. 5A top panel; Fig. S11A, SI).

Finally, the trends in the change in growth with carbon concentration are also different between monocultures and cocultures in their respective conditions - growth can either increase or decrease with an increase in carbon concentration in cocultures whereas it only increases with carbon concentration in monocultures. In terms of interactions between *E. coli* and *C. reinhardtii*, this can be stated as follows: while only enhancement of *E.coli* growth is observed as carbon concentration increases in monoculture, the effect on *E. coli* by *C. reinhardtii* in coculture as carbon concentration increases becomes inhibitory at low pH and high buffering capacity, but facilitatory at high pH and low buffering capacity (evidenced by the positive interaction coefficient of *pH*[*C*] and the negative interaction coefficient of *BC*[*C*] obtained from regressing *E. coli* growth, purple bars in Fig. 5A top panel). This example illustrates that *C. reinhardtii* modulates the capacity of *E. coli* growth on carbon in a manner that depends on pH and buffering capacity of the environment.

Algal abundance dynamics also depend strongly on pH and buffering capacity. The regression coefficients for predicting algal growth on glycerol in monoculture and co-culture are shown in Fig. 5A bottom panel. In this regression, we observe a similar interplay between pH and buffering capacity and nutrients’ concentration i.e the monoculture coefficients of *pH*[*P*], *pH*[*C*], *BC*[*P*], and *BC*[*C*] (green bars in the bottom panel (Fig. 5B)), are all non-zero and statistically significant, showing the presence of modulation effect of pH and buffering capacity on nutrient concentration in determining *C. reinhardtii* growth in monoculture. Here again, the largest monoculture coefficient is for the *BC*[*P*] term indicating an increase in the growth of *C. reinhardtii* with phosphorus concentration and buffering capacity. While the growth of *C. reinhardtii* is known to increase with phosphorus concentration [35], we speculate that the increased phosphorus uptake leads to increased N utilization (the N source here is ammonium). Ammonium utilization by algae causes acidification of the environment [36], which is known to negatively affect the growth of *C. reinhardtii* [37]. Therefore, We reason that the environments with high buffering capacity potentially prevent this acidification and hence favor increased growth of *C. reinhardtii*, as reflected in the high coefficient of *BC*[*P*].

The modulation of algal growth by bacteria also depends on pH and buffering capacity in a fashion similar to what we observe with bacteria. For example, the interaction coefficients of *BC*[*P*] and *BC*[*C*] (cyan bars in the bottom panel of Fig. 5B), being significant means that the impacts of *E. coli* on *C. reinhardtii* growth is mediated by an interplay between buffering capacity and nutrient concentration. In other carbon sources, the impacts of *E. coli* on *C. reinhardtii* growth mediated by an interplay between both pH and buffering capacity and nutrient concentration are observed (Fig. S10, SI).

Overall, the result that the interactions between algae and bacteria are mediated by pH and buffering capacity, through their differential impacts on nutrient dependence on monoculture and coculture growth holds across carbon sources (Fig. S10, SI)

### Effect of environmental factors on algae-bacteria interactions depends on the identity of carbon source

Finally, we investigated if the dependence of algae-bacteria interactions on the environmental factors - pH, buffering capacity, phosphorus concentration, and carbon concentration, is further modulated by the identity of the carbon source available in the communities. Between several carbon source pairs, we found some apparent differences in the effect of the environmental factors on algae-bacteria growth. For example, differences in several of the monoculture and interaction coefficients (which quantify the effect of environmental factors on growth and interactions) between glucose and galactose are clearly observed (Fig. 6A). While the feature *BC*[*C*] has the highest effect in predicting *E. coli* growth in the case of glucose, *BC*[*P*] is the feature with the highest importance in the case of galactose. And the effect of *BC*[*C*] in predicting the *E. coli* growth is the opposite between glucose and galactose. Additionally, for *E. coli*, the coefficients of [*P*] and [*C*] show different patterns in glucose and galactose, with generally negative coefficients in glucose and coefficients of opposing sign for monoculture and interaction coefficients in galactose. Qualitatively similar patterns are observed in coefficients describing algal growth (Fig. S10, Fig. S13, SI). These observations suggest that the identity of the carbon source modulates how environmental factors impact algae-bacteria interactions.

**Figure 6.**
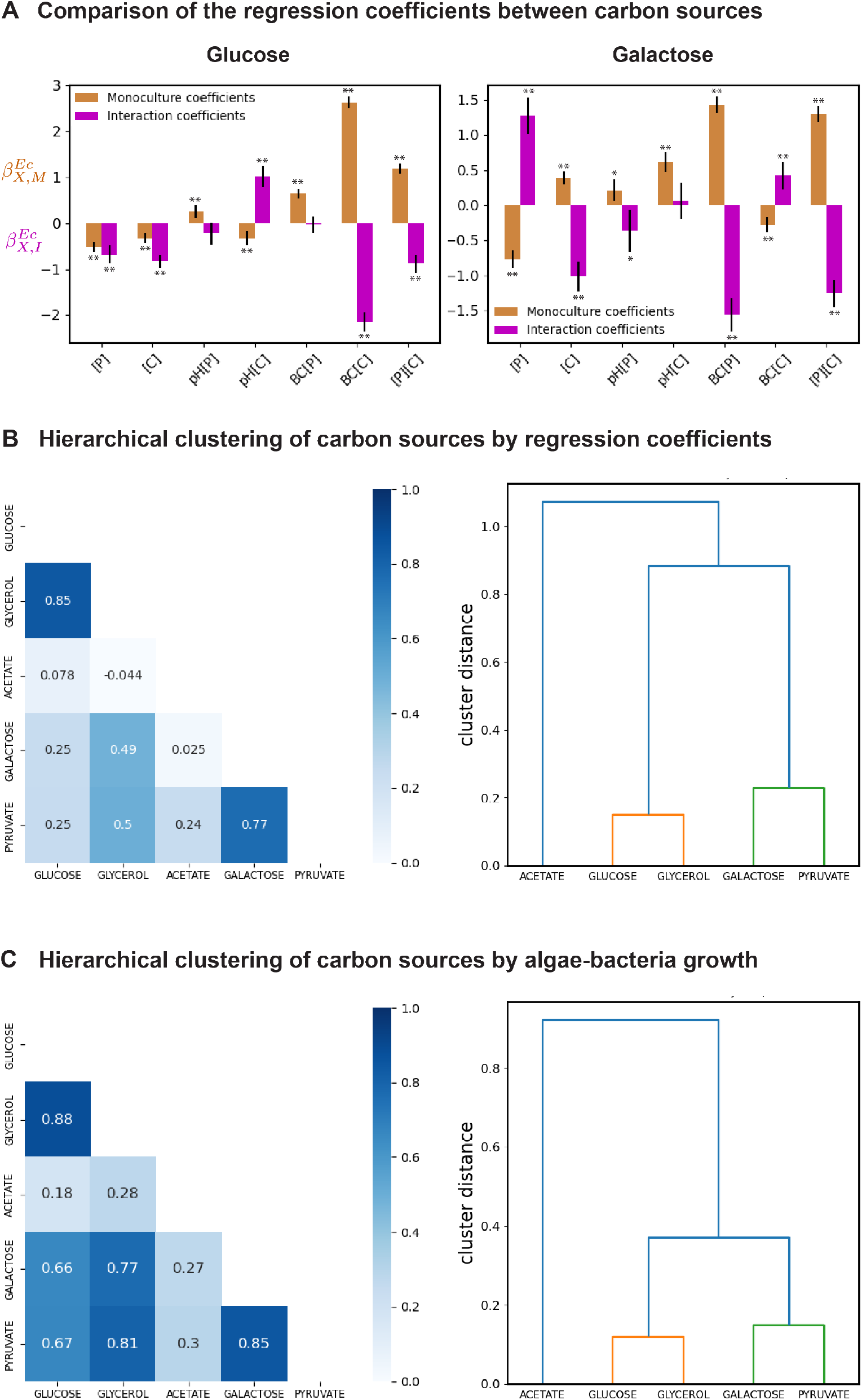
Effect of environmental factors on algae-bacteria interactions depends on the identity of carbon source. (A) Comparison of the regression coefficients between glucose and galactose. The monoculture coefficients 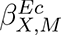 (brown bars) and the interaction coefficients 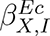 (magenta bars) of the corresponding features on the x-axis obtained from the regression model predicting the growth of *E. coli* in monocultures and cocultures for glucose (on the left) and galactose (on the right). ** indicates a p-value *<*0.001 and * a p-value *<*0.05. (B) Hierarchical clustering of carbon sources by the monoculture and interaction coefficients obtained from the regression models predicting the growth of *E. coli* and *C. reinhardtii*. The correlation matrix computed for the hierarchical clustering on the left and the resulting dendrogram on the right (See Section 9, SI). (C) Hierarchical clustering of carbon sources by the median growth of algae and bacteria in monocultures and cocultures in all the environmental conditions. The correlation matrix computed for the hierarchical clustering on the left and the resulting dendrogram on the right (See Section 9, SI).

To interrogate these patterns further, we classified carbon sources based on their modulation of the effect of the environmental factors on algae-bacteria growth. To do this, we computed correlations between the regression coefficients (which quantify the effect of environmental factors on growth and interactions) obtained for predicting algae-bacteria growth, between all pairs of carbon sources. We performed hierarchical clustering of the carbon sources based on the monoculture and interaction coefficients of [*P*],[*C*],*pH*[*P*],*pH*[*C*],*BC*[*P*],*BC*[*C*] and [*P*][*C*], obtained from the regressions for the carbon sources (Section 9, SI). The correlation matrix computed for the hierarchical clustering showed that glycerol is most similar to glucose, galactose is most similar to pyruvate, and acetate has no strong correlation with any of the carbon sources (Fig. 6B left panel). Therefore, hierarchical clustering identified three clusters of carbon sources in our dataset, with glucose and glycerol forming one cluster, and galactose and pyruvate forming another cluster, and acetate forming a cluster of its own (Fig. 6B right panel).

We wondered why these different carbon sources would have such divergent impacts on interactions. We suspected that bacterial utilization of distinct carbon sources could have differing impacts on pH. To test this idea, we grew *E. coli* in plates in each of the 5 carbon sources and measured the final pH. We found that glucose and glycerol both showed large drops in pH while the other three carbon sources did not (Section 10, SI). Thus, we speculate that heterotrophic utilization of organic carbon might play a key role in modulating pH and thus the interactions between algae and bacteria.

Finally, we wanted to check whether this result was dependent on the details of the regression formalism we defined for quantifying growth across environments. To do this, we quantified similarities in growth across environments in a model-independent fashion. We classified carbon sources based on the similarity in algae-bacteria growth. The classification of the carbon sources was done by computing the correlation between carbon sources in algae-bacteria growth across all the environmental conditions and culture conditions (Fig. 6C; Section 9, SI). Here again, we found the carbon sources within the same clusters - glycerol and glucose, and galactose and pyruvate, to have the greatest correlation in the algae-bacteria growth with each other than with any other carbon sources. We concluded that this apparent clustering of carbon sources does not depend on details of our model specification.

## DISCUSSION

By using a high-throughput droplet microfluidic platform, we were able to perform a massively parallel screening of algae-bacteria interactions in several hundreds of environmental conditions varying in pH, buffering capacity, phosphorus availability, carbon availability, and carbon source identity. To our knowledge, this is the largest screen exploring the combintorial effect of environmental factors on phototroph-heterotroph interactions in a systematic way via a bottom-up approach. Studies in the past have tested for the effect of nutrient availability on phototroph-heterotroph relationships, but have been mostly limited to only a handful of nutrient types/availabilities or have involved uncontrolled experimental conditions such as uncharacterized phototrophic and heterotrophic species, often in the presence of organisms from other trophic levels [12, 16, 38, 39]. Our observation of the complex dependence of algae-bacteria interactions on environmental factors underscores the importance of undertaking such high-dimensional studies. This is especially important in light of the chemical complexity of environments wild microbial communities are exposed to [8].

Our study is also novel with respect to exploring the effect of the chemical properties of the environment - pH and buffering capacity, on algae-bacteria interactions. The central finding of the study is that pH and buffering capacity significantly alter algae-bacteria interactions by manipulating the impact of nutrient availabilities on growth. For most carbon sources, the role of pH and buffering capacity in determining algae-bacteria interactions were comparable to, or significantly higher than, the effect of nutrient availabilities alone, underscoring the importance of the effects of pH and buffering capacity on algae-bacteria interactions. This result suggests that chemical factors in the environments play an important role in mediating phototroph-heterotroph interactions which are largely considered as being driven by resource exchange and competition [12–14, 16, 19, 25, 34].

Recently, microbial ecologists have encouraged the use of statistical modeling approaches to derive general governing principles in ecology [40, 41]. In this regard, we highlight the mechanistic insights provided by the statistical framework implemented here to predict algae-bacteria growth. Our statistical approach for predicting algae-bacteria growth in different environments permitted us to dissect the contribution of the different environmental factors on the inter-species interactions. Even though our modeling approach is largely agnostic to the detailed mechanisms of the effect of environmental factors on algae-bacteria interactions, we find that the regression results do align qualitatively with some known processes. For example, *E. coli* can acidify its environment when growing on glycolytic substrates at sufficiently high growth rates through the process of overflow metabolism [42]. In this case, the bacterium could be acidifying the medium in conditions where buffering capacity is weak and carbon levels are relatively high. However, overflow occurs at relatively high growth rates of approximately 0.71/h-0.81/h, and microtiter measurements indicate that our strain

in these conditions grows slower than this (Section 10, Table S5, SI). Further, our measurements cannot accurately capture bacterial growth rates in droplets due to limited temporal sampling, but we cannot rule out the possibility that overflow causes growth to modify pH in the droplets. Similarly, it is known that *C. reinhardtii* will acidify the environment due to ammonia uptake and this may also play a role in the importance of pH and buffering capacity in determining growth in these experiments. It remains an important avenue for future work to uncover the mechanisms underlying the interactions discovered here. Our hope is that large-scale screens like those enabled by this platform can contribute new insights into the mechanisms by which environmental factors contribute to algae-bacteria interactions.

Our exploration of the impact of carbon source identity on algae-bacteria interactions showed that the effect of the environmental factors - pH, buffering capacity, and nutrient availability, on the interspecies interactions depends on the carbon source identity. This result suggests that the chemical identity of the available reduced organic carbon plays a key role in determining how algae-bacteria interactions play out. Therefore, considering the role of individual nutrients such as phosphorous [43] in these interactions might be too simple a picture. Additionally, our analyses revealed three groups of carbon sources, showing that the impact of the environmental factors - pH, buffering capacity, and nutrient availability, on algae-bacteria interactions was approximately conserved between the carbon sources within the same group. Such an apparent similarity between the different carbon sources within the groups hints that there may be some relatively simple structure in how the carbon source identity and the other environmental factors conspire to determine the outcome of an interaction. Whether this is the case or not awaits a broader survey of additional carbon sources, mixtures of carbon sources, and a deeper mechanistic understanding of the physiology underlying these processes.

While kChip offers a massive throughput advantage to perform a screen of this magnitude, the interactions inferred in the confined environments of droplets on the kChip could potentially differ from the interactions in the well-mixed, open, environments in the lab or the wild. For example, the rate of gas exchange, particularly O_2_, and CO_2_ will determine respiration, photosynthesis, and pH and thereby modulate interactions in the droplets. In fact, a recent microfluidic-based study has shown that droplet size substantially modifies the degree of syntrophic interaction between bacterial species [44]. Consistent with these findings, we observe differences in bacterial growth between microtiter plates and droplets (Fig. S14, SI). Hence, it remains an important avenue for future work to understand how confinement impacts the algae-bacteria interactions observed here, as this process could well be important in the wild.

As our study of phototroph-heterotroph interactions was undertaken in a community of algae and bacteria that are not known to associate in the wild, it remains to be seen how our results relate to communities of phototroph and heterotroph with wild associations and shared evolutionary history. For example, the mechanism by which *C. reinhardtii* inhibits *E. coli* growth is not precisely known, and it is unclear whether other bacterial taxa would also be subjected to similar strong inhibitory effects. Studies between several strains of the phototroph, *Prochlorococcus*, and of oligotrophic and copiotrophic bacteria, have revealed strain-dependant interactions [45, 46]. Thus, it would be interesting to repeat these experiments with a broader sampling of bacterial taxa including those that are known to associate with the alga in the wild [47]. By expanding this study to wild associations, we would hope to more broadly capture the relevance of these findings for consortia in complex environments.

