## Supplemental Appendix for "Environmental modulators of algae-bacteria interactions at scale"

### Supplementary Appendix: *Environmental modulators of algae-bacteria interactions at scale*

**9 Hierarchical clustering of carbon sources** **16**

**10 Plate experiment assaying *E. coli* growth on carbon sources** **17**

### 1 Strains

The heterotroph was a bacterium, *Escherichia coli*, strain MG1655 (Coli Genetic Stock Center (CGSC) #8237), which was transformed to constitutively express a green fluorescent protein (GFP) on a plasmid (protocol below). We used a fluorescent protein coded on a plasmid to increase fluorescence intensity per cell which we found to be too low for the imaging modalities used here when the protein was genomically integrated. The bacteria were cryogenically preserved at -80 °C. The phototroph in the study was an alga, *Chlamydomonas reinhardtii*, strain UTEX2244 obtained from the University of Texas Austin Culture Collection of Algae `utex.org`. Algae were cryogenically preserved in liquid nitrogen `https://utex.org/pages/cryopreservation#liquid`.

### 2 Plasmid transformation

The transformation of the wild type strain of *E. coli* MG1655, previously used in [1], to express GFP on a plasmid, was done to enable the measurement of bacterial abundances via fluorescence microscopy. Firstly, the plasmid for the transformation was extracted from the *E. coli* strain, DH10B pZA 1R GFP [2], following the protocol in the GeneJET Plasmid Miniprep kit #K0503 for the low copy number plasmids. Following this, the transformation protocol involved the following steps:

#### 2.1 Preparation of electrocompetent cells

The wild-type MG1655 cells were grown from frozen stocks in a 5 mL overnight culture of Lysogeny Broth (LB) at 30 °C in a shaker incubator. 1/2 mL of the overnight culture was added to a flask containing 30 mL of LB and grown at 30 °C with shaking at 200 RPM until the OD600 reached 0.5-0.7. The flask was removed and the culture was cooled by swirling in an ice water slurry for five minutes, then placed on ice for ten minutes. The culture was transferred to a pre-chilled centrifuge tube and pelleted by centrifugation (5 min, 5000 RPM) in a refrigerated centrifuge chilled to 4 °C. The supernatant was dumped and the cells were washed in 10 mL of ice-cold 10% glycerol. Pelleting was repeated in the same way and two more glycerol washes were performed, followed by a final resuspension in 200 µL. The cells were immediately placed on ice and kept cold until electroporation.

#### 2.2 Electroporation of plasmid

100 µL of the prepared electrocompetent cells were mixed with 5 µL of the extracted plasmid mix in a pre-chilled microcentrifuge tube before being transferred to a pre-chilled 0.1 cm gap electroporation cuvette (USA Scientific) and electroporated at 2 kV in an Electroporator 2510

(Eppendorf). 1 mL of LB was immediately added and 1h outgrowth at 30 °C with shaking was allowed before plating on an LB + ampicillin plate, which was grown overnight at 30 °C. A colony from the plate was grown overnight (30 °C, shaking) in 5 mL of LB+ampicillin, and a frozen glycerol stock of MG1655+GFP was created from the culture.

### 3 Media

#### 3.1 Modified Taub medium and nutrient sources

Modified Taub formed the base media in our experiments. Taub media is a freshwater mimic media that was originally created to support co-cultures of *Daphnia pulex* and *Chlorella pyrenoidosa* [3]. Several previous studies of microbial ecosystems that used the 1/2X Taub media with undefined carbon and nitrogen sources (proteose peptone) [4, 5, 6] demonstrated the ability of both *E. coli* and *C. reinhardtii* to grow on Taub. However, to probe the effect of nutrient concentration and nutrient sources on algae-bacteria interactions, we required the media in our experiments to be chemically defined. Hence, the undefined Taub media was modified to include chemically defined carbon and nitrogen sources in place of the proteose peptone, similar to that in [7]. $NH_4Cl$ formed the nitrogen source in all our experiments and one of the five sources - glucose, glycerol, galactose, acetate, and pyruvate formed the carbon source in our experiments. The phosphate source in the media was also replaced with an equal mix of potassium phosphate monobasic ($KH_2PO_4$) and potassium phosphate dibasic ($K_2HPO_4$) salts. Modified 1X Taub stock was first prepared by removing the small amount of phosphate that is traditionally present in 1X Taub solution. It was later supplemented with buffers and nutrient sources at different concentrations to generate the desired environmental conditions for the kChip experiments (discussed below). The chemical composition of the modified 1X Taub medium is shown in Table S1.

#### 3.2 Preparation of environmental conditions for the kChip experiments

For each carbon source, 16 environmental conditions varying in initial pH, buffering capacity, phosphorus concentration, and carbon concentration were prepared. The values of the environmental factors were chosen to be in the biological range: 6.1-7.5 for initial pH, 0-3.5 mM for buffering capacity, 0.01 mM - 4 mM for phosphorus concentration, 2 mM - 10 mM for carbon concentration. The environmental conditions were also barcoded at the time of their preparation by adding the three fluorescent dyes Alexa Fluor 555 (Thermo- Fisher Scientific A33080), Alexa Fluor 594 (Thermo- Fisher Scientific A33082), and Alexa Fluor 647 (Thermo- Fisher Scientific A33084) such that each environmental condition gets a unique combination of the dyes with the total dye concentration always summing to 1 $\mu$M (The dye concentrations will be later used to infer the environmental conditions of the communities formed on the kChip as discussed in section 6.2) The initial pH, buffering capacity, nutrient concentrations along with the dye concentration values of the 16 environments are reported in Table S2. Note that the reported pH and buffering capacity values are not the measured but estimated values from a model we developed and experimentally validated (details in section 7). Additionally, these values are reported for environments having glucose/glycerol/galactose as the carbon source. When acetate/pyruvate is the source of carbon in the environments, the pH and buffering capacity slightly differ from those reported here and are computed as described in section 7. Depending upon the carbon source, the appropriate values of pH and buffering capacity are used in all the analyses.

To prepare the 16 environmental conditions, we first prepared the following sets of stock solutions -

1. Stock solutions of the base media 1X Taub:
   **B1** - The modified 1X Taub media with 0.11% w/v bovine serum albumin (BSA)
   **B2** - The modified 1X Taub media with 20 mM MOPS buffer and titrated to a pH of 6.95 with 0.11% w/v BSA
   **B3** - The modified 1X Taub media with 30 mM Tris buffer and titrated to a pH of 7.5 with 0.11% w/v BSA
   (BSA is added to the media to improve retention of the fluorescent dyes in droplets on the kChip)

2. Stock solutions of carbon:
   **C1** - $\tilde{4}$17 mM of glucose/glycerol/galactose/pyruvate/acetate by moles of carbon
   **C2** - $\tilde{6}$7 mM stocks glucose/glycerol/galactose/pyruvate/acetate by moles of carbon

3. Stock solution of Nitrogen:
   **N1** - 50 mM of $NH_4Cl$ stock

4. Stock solutions of Phosphate:
   **P1** - 50 mM of phosphate stock ($KH_2PO_4$+$K_2HPO_4$ in 1:1 ratio by moles)
   **P2** - 1 mM of phosphate stock ($KH_2PO_4$+$K_2HPO_4$ in 1:1 ratio by moles)
   **P3** - 0.12 mM of phosphate stock ($KH_2PO_4$+$K_2HPO_4$ in 1:1 ratio by moles) titrated to a pH of 7.5
   **P4** - 12 mM of phosphate stock ($KH_2PO_4$+$K_2HPO_4$ in 1:1 ratio by moles) titrated to a pH of 7.5

5. Stock solution of Alexa Fluor 555:
   **D1** - 25 $\mu$M of Alexa Fluor 555 dye

6. Stock solution of Alexa Fluor 594:
   **D2** - 25 $\mu$M of Alexa Fluor 594 dye

7. Stock solution of Alexa Fluor 647:
   **D3** - 25 $\mu$M of Alexa Fluor 647 dye

These stock solutions formed the key components in setting the various properties of the environmental conditions - initial pH, buffering capacity, nutrient concentrations, and barcodes. The prepared stock solutions were mixed in the desired ratios using a liquid handling robot Opentrons OT-2 to obtain the 16 environmental conditions. The stock solutions and their volumes used for making each of the 16 environments are reported in Table S3. 480 $\mu$L of each of the 16 environments were prepared such that we obtained the indicated concentrations of nutrients and dyes (in Table S2) when 10 $\mu$L of the *E. coli* and *C. reinhardtii* cells suspended in the modified 1X Taub media were later independently added to 240 $\mu$L of each of the environments. In the 480 $\mu$L of the environments, 230 $\mu$L was composed of one of the three base media stock solutions - B1/B2/B3, and the rest of the 250 $\mu$L was made up of stock solutions of carbon, nitrogen, phosphorus, and the dyes, depending on the media type.

Specifically, in environments E1-E8 (Table S3), the modified and unbuffered 1X Taub media, B1, formed the 230 $\mu$L volume of the 480 $\mu$L. E4-E8 differed from E1-E4 in the phosphate stock used. While untitrated phosphate stocks P1 and P2 were used to get the desired phosphorus levels in E1-E4, the titrated phosphate stocks P3 and P4 were used to obtain the desired phosphorus levels in E4-E8. The use of the phosphate stocks P3 and P4 titrated to a pH of 7.5 (greater than ~7 - the ~pH of the stocks P1 and P2) caused the environments E7 and E8 to have higher pH than E1-E4. And, in the environments, E9-E12 and E13-E16, the buffered 1X Taub media B2 (having the MOPS buffer) and B3 (having the Tris buffer) respectively formed the 230 $\mu$L volume of the 480 $\mu$L. The strong buffers- Tris and MOPS were chosen to obtain the environments E9-E16 with high buffering capacities (~3.5 mM). And the low initial pH of the environments E9-E12(~6.9) is due to the modified 1X Taub media buffered with MOPS and titrated to a low pH of 6.95. Similarly, the higher initial pH of the environments E9-E12 (~7.4) is due to the modified 1X Taub media buffered with Tris and titrated to a high pH of 7.5. The pKa values of MOPS and Tris (7.1 and 7.9 at 30 °C) make them ideal choices as buffering agents at low pH and high pH respectively. Lastly, as is reported in Table S2, we chose higher phosphorus levels (0.03-3 mM) for the lower buffering capacity environments E1-E8 but lower phosphorus levels (0.01-0.08 mM) for the environments E9-E16 which have higher buffering capacity. This was because the source of phosphorus in our experiments (i.e $KH_2PO_4$+$K_2HPO_4$ in 1:1 ratio by moles) acts as both a nutrient and a buffer, for example, against the potential acidification of the environment arising from carbon metabolism by *E. coli* [8]. As a result, lower phosphorus levels were sufficient to give rise to appreciable growth of *E. coli* in monocultures in the high buffering capacity environments whereas higher phosphorus levels were required in the lower buffering environments to result in the similar growth of *E. coli* in its monocultures. Therefore, our choice of different phosphorus levels at the different buffering capacities allowed us to investigate differences in the algae-bacteria interactions between environments giving rise to similar growth of *E. coli*.

### 4 Culturing and harvesting of the microbes for kChip experiments

#### 4.1 Culturing

Before beginning the experiment, the bacteria and algae were cultured separately in distinct media, with both microbes undergoing two growth cycles in their respective media.

Bacteria were cultured from a freezer stock in 5 mL lysogeny broth (LB) with ampicillin added at a concentration of 50 $\mu$g/mL to retain the plasmid. The culture was incubated at 30 °C (New Brunswick Scientific C24 Incubator-Shaker), shaking at 200 RPM for ~16 hrs. It was then passaged into fresh 5 mL LB + 50 $\mu$g/mL ampicillin at 2500X dilution and grown for ~24 hrs at 30 °C shaking at 200 RPM, before finally harvesting for the experiments.

The alga, *C. reinhardtii* was cultured in a 30 °C shaker-incubator (New Brunswick Scientific C24 Incubator-Shaker), shaking at 200 RPM with 68.5 $\mu$mol $m^2s^{-1}$ illumination in 10 mL Tris-Acetate-Phosphate (TAP) media, inoculated directly from a freezer stock. TAP is a defined media with acetic acid as a carbon source `https://www.chlamycollection.org/methods/media-recipes/tap-and-tris-minimal/`. After ~4 days, the algal culture was passaged into fresh 20 mL of TAP media at 250X dilution and grown for ~3 days at 30 °C shaking at 200 RPM, before finally harvesting for the experiments.

#### 4.2 Cultures preparation

Prior to setting up the experiment on kChip, the harvested microbial cultures were washed thrice into modified 1X Taub media.

1 mL of the MG1655 culture was centrifuged in an eppendorf at 3000 RPM (eppendorf centrifuge 5417R) for 5 mins. The supernatant was immediately discarded and the pellet was resuspended in 1 mL of fresh modified 1X Taub. This process was repeated thrice and OD590 of the final suspension was adjusted to obtain 0.005 in the droplets by diluting it with the modified 1X Taub media.

20 mL of the UTEX 2244 culture was also centrifuged thrice in 20 mL falcon tubes at 500 RCF for 10 mins. The culture was concentrated sequentially after every centrifugation from 20 mL to 7.5 mL to 2 mL. By further concentration or dilution, the OD750 of the final suspension was adjusted to obtain 0.145 in the droplets by diluting it with the modified 1X Taub media. The optical densities were measured using the BioTek Synergy HT microplate reader.

### 5 Setting up the experiments on kChip

#### 5.1 Droplet preparation

The cultures of *E. coli* and *C. reinhardtii* that were washed into the modified 1X Taub media and with their ODs set were independently added to the 16 barcoded environments of one of the five carbon sources at 25X dilution.Each of the environments was thoroughly mixed using an electronic pipettor by pipetting up and down at least three times to ensure thorough mixing of the barcode dyes and the cells. 20 $\mu$L aliquots of these environments harboring the *E. coli* and *C. reinhardtii* cells independently were transferred to a Bio-Rad QX200 cartridge and were emulsified into ~20,000 1 nl droplets in fluorocarbon oil (3M Novec 7500) stabilized with 2% (w/v) fluorosurfactant (RAN Biotech 008 FluoroSurfactant). For each carbon source, there were 32 kinds of droplets - 16 environmental conditions each having cells of *E. coli* and *C. reinhardtii* separately.

#### 5.2 Setting up the kChip platform

The generated droplets of all the 16 environmental conditions having cells were pooled together into a 1 mL Eppendorf and mixed by pipetting up and down with a 200 $\mu$L pipette. 180 $\mu$L of the pooled and mixed droplets were loaded into kChip(k=2) as described in [9]. kChip is made of PDMS and contains an array of ~25,000 microwells each of which can take two droplets (~130 $\mu$m in diameter). Briefly, the kChip was suspended in the chip loader made of acrylic, such that a ~300–500 $\mu$m flow space was created between the chip and a hydrophobic glass substrate. The flow space was filled with fluorous oil (3 mL 3M, 7500) prior to loading, followed by the addition of the droplet pool to the loading slot. By flushing the flow space with oil, the droplets were made to spread around in the flow space and enter the microwells due to buoyancy. Also, the loader was tilted to further facilitate the movement of the droplet foam within the flow space until the microwells were filled with droplets. The flow space was then again replenished with 3 mL of the fluorous oil. On the side, a fresh MicroAmp Optical Adhesive film (ThremoFisher #4311971) was laid out on the bench with its sticky side facing up and wetted with ~1 mL of the fluorous oil. The kChip was carefully lifted off the acrylic loader and sealed with the film by running the chip against the wetted film on the edge of the bench.

The kChip was then imaged to infer the barcode identities and the starting cell densities in the wells (imaging discussed in section 5.3). Following this, the droplet pairs in the microwells were merged by running the tip of a corona treater (Model BD-20, Electro-Technic Products) over the sealed side of the chip for 10 seconds. The merging of the droplets resulted in the formation of monocultures and cocultures of algae and bacteria in all environmental combinations of the initial 16 environments. Overall, 3 culture types (*E. coli* monoculture, *C. reinhardtii* monoculture, *E. coli* - *C. reinhardtii* coculture) in 105 environments were generated upon droplet merging for each of the carbon sources, with the number of replicates ranging from 3 to 330 (The median number of replicates ranged from ∼30-85 depending upon the culture type). The kChip was then transferred with its film side facing up and covered with a glass slide, into a Ziploc bag containing a moist towel to maintain high humidity and minimize evaporation. The entire setup was housed in an environmental chamber at 30 °C and illuminated with a bulb (Utilitech pro L9PAR20/LEDG5) at (68.5 $\mu$mol $m^2s^{-1}$, measured with LED light meter PCE-LED 20). The kChip was imaged at 12 h, 21 h, 45 h, and 68 h from the time of the first scan. For each carbon source, a separate kChip experiment was set up.

##### 5.3 Fluorescence Microscopy

A widefield fluorescence microscope (Axio Observer.Z1) with X-CITE 120 lamp (Excelitas Technologies #012-63000) as the light source for fluorescence imaging, was used to scan the kChip for barcodes and the growth of the microbes. Images were acquired with a 5X/0.16 NA objective (Zeiss EC Plan-Neofluar) with FOV (Field of view) of 2.47×2 mm, which required collecting 644 images to scan one full kChip area covering all the microwells. Images were collected by a camera (Axiocam 506 monochromatic) at a bit depth of 14 with 5 × 5 binning and at an exposure time of 50 ms. The following filter sets were used to detect the five fluorophores: Alexa Fluor 555: Semrock Brightline SpOr-B-CSC-ZERO; Alexa Fluor 594: Omega optical Excitation filter-XB102/Dichroic-XF2014/Emission filter-XF3028; Alexa Fluor 647: Semrock, Brightline Cy5-4040B-CSC-ZERO; GFP: Zeiss filter Set 38 HE; chlorophyll: Chroma Technology 31017. The lamp power was manually set to obtain ∼71 mW with the Alexa Fluor 555 filter/∼8 mW with the Alexa Fluor 594 filter/∼27 mW with the Alexa Fluor 647 filter, measured at 540 nm/590 nm/630 nm respectively using a Thorlabs power meter (with power sensor S121B). In addition to the fluorescence images, brightfield images were also acquired with a TL Halogen lamp (set to 1.51 V) as the light source at an exposure of 1.1 ms. The duration of an entire scan was about 50 min.

#### 6 Image processing and analysis

The tiled images acquired at each time point were stitched together to form a single image of the entire chip having all the microwells, using the stitching module in the Zeiss Zen blue image analysis software. Also, the stitched images across the time points were aligned by manually estimating the rotation and the shift in the chip at each time point with respect to the image acquired at the first time point, and correcting for them using the rotate and shift features in the zen software. The aligned images were then used for further processing and analyses in Python. First, the aligned images were computationally redivided and cropped in Python to obtain 644 tiles with 10% overlap as processing a single large image would require too much memory. From here on, the image analysis pipeline involved (a) Correcting for chlorophyll bleed-through in the A647 image (see below); (b) Inferring barcodes to identify the environmental conditions in the droplet pairs in each microwell using the three fluorescence dye signals; (c) estimating abundances of *E. coli* and *C. reinhardtii* in all the environmental conditions. All analyses were performed with either custom Python scripts, or code obtained from [9, 10, 11].

#### 6.1 Correcting for chlorophyll bleed-through in the Alexa Fluor 647 images

Inspection of the microscopy images of the fluorophores showed algal cells to appear in the images acquired with the Alexa 647 filter (Fig. S1 (left panel)). This bleed-through of the chlorophyll signal into the Alexa Fluor 647 channel is due to the overlap between the fluorescence spectrum of the chlorophyll pigment and the Alexa fluor 647 dye. The chlorophyll signal bleed-through into the Alexa Fluor 647 images would corrupt the barcode clustering process (discussed in 6.2), which is crucial for identifying the environmental conditions formed on the chip. A computational solution was developed to address this issue that involved the following steps:

1. Apply sobel transform (using scikit-image) to the Alexa 647 image to find the edges of the algal cells
2. Obtain the mask of the sobel transformed image to extract the edges of the algal cells.
3. Use a gaussian filter (with sigma = 1 pixel, in SciPy) to set the intensities of the pixels within the edges in the mask to greater than 0.
4. Set all the pixel intensities greater than 0 to NaN. This step would essentially set the intensity of all the pixels corresponding to the algal cells in the mask to NaN.
5. Multiply the mask obtained in step 4 with the original Alexa 647 image with the chlorophyll bleed-through and using a 2D interpolation scheme (interpolate.griddata in SciPy), estimate the intensity values of the pixels that were set to NaN. The Alexa 647 image obtained after this correction algorithm is free from the bright signal from the chlorophyll fluorescence (Fig. S1 (right panel)).

#### 6.2 Inferring barcodes to identify environmental conditions

Following the correction of Alexa 647 images for chlorophyll bleed-through, the three dye channel images were analyzed to detect the barcodes of the droplets in the wells of kChip and thereby infer the environmental conditions formed in the microwells. Similar to the pipeline in [9], the algorithm began with creating images by summing up the three dye channels images and then applying a circular hough transform (scikit-image) on the summed images to detect the circular droplets in the wells. Using the positions of the droplets reported by the hough transform, the three-color dye fluorescence intensities of each of the droplets were extracted. The fluorescence of a dye in a droplet was calculated as the median of the pixel values in the respective dye's image in a square of size 10×10 pixels at the droplet center, after locally subtracting for the background fluorescence intensities. The three dye channel images were also smoothed by applying a median filter (SciPy) with a kernel size of 8 pixels, prior to computing the dye fluorescence intensities. Obtaining the three-color fluorescence intensities of all the droplets in this manner, the intensities were projected to a 2-dimensional plane on the basis of the constraint that the intensities summed to a constant (as the sum of the dye concentrations is a constant equal to 1 $\mu$M). The clusters of droplets formed in the 2-dimensional plane based on the dye ratios were identified by bounding the data points with manually defined polygons (using matplotlib.path). Finally, using the apriori knowledge of fluorescence-dye-ratios to environmental conditions mapping (from while designing the environmental conditions and barcoding), the droplets/clusters were assigned to the environmental conditions. This knowledge of the droplet positions and their environmental conditions allowed inference of the environmental conditions of the communities formed in the different microwells after the merging of the droplets.

#### 6.3 Analyses for estimating abundances of algae and bacteria

##### 6.3.1 Computing local background GFP and chlorophyll intensities

To account for any spatial and temporal variation in the background intensities in the images, we computed the background fluorescence intensities in the GFP and chlorophyll images locally. We defined rectangular regions around the droplets/wells at each of the time points. Then the local background GFP/chlorophyll intensity for a given droplet/well was obtained as the median of the top 5% of the pixel intensities in the GFP/chlorophyll images respectively in the region bounded by the rectangle but excluding the droplet/well area containing the cells.

##### 6.3.2 Detection of algae and bacteria cells

The GFP and the chlorophyll images were segmented to detect algal and bacterial cells by intensity thresholding the original images on a well-by-well basis. The GFP threshold for any given well was set to 100-pixel intensity units above the local background GFP intensity computed for that well (from above). Likewise, the chlorophyll threshold for any given well was set to 500-pixel intensity units above the local background chlorophyll intensity computed for that well. We refer to the disconnected regions of GFP/chlorophyll pixels in the segmented images of GFP/chlorophyll as GFP/chlorophyll clusters respectively. These GFP/chlorophyll clusters represent the aggregated or planktonic cells of *E. coli* or *C. reinhardtii.* Using scikit-image (regionprops), we extracted the area, total fluorescence intensity, and mean fluorescence intensity (i.e intensity per pixel) of the GFP and chlorophyll clusters, used in the further analyses below.

##### 6.3.3 Estimating single-cell intensities

To estimate the single-cell intensities of *E. coli* and *C.reinhartdii*, we first estimated the typical areas of a single cell of *E. coli* and *C. reinhardtii.* The distribution of the areas of the GFP and chlorophyll clusters at the first and the last time point across the kChip were plotted (Fig. S2). By visual investigation of these distributions, we inferred the typical areas of a single *E. coli* cell and a single *C. reinhardtii* cell at the first time point and the last time point to be around the peak of the distributions as marked in Fig. S2. On average, *E. coli* showed a decline in the single-cell areas in all the carbon sources. On the other hand, *C. reinhardtii* showed a lower reduction in the single-cell areas and only in the case of acetate and galactose.

Using the typical areas of single cells, we were able to estimate the single-cell intensities of *E. coli* and *C. reinhardtii.* To account for any difference in the single-cell intensities between monoculture and coculture, we obtained estimates of the single-cell intensities in monoculture and coculture separately. This was done by first classifying the wells as having monoculture or coculture communities as follows -

1. If a well only has GFP clusters but no chlorophyll clusters at the first time-point, the well has an *E. coli* monoculture community

2. If a well has no GFP clusters but only chlorophyll clusters at the first time-point, the well has a *C. reinhardtii* monoculture community

3. If a well has both GFP clusters and chlorophyll clusters at the first timepoint, the well has a *E. coli* - *C. reinhardtii* coculture community

Then, the steps for estimating the single-cell intensities for *E. coli* and *C. reinhardtii* in monoculture and coculture involved:

**Computing single-cell areas of *E.coli* and *C.reinhardtii* in monoculture and coculture**: Area of a single *E. coli* cell $A_{Sc}^{Ec}$ in monoculture/coculture was estimated as the median of the areas of GFP clusters in monocultures/cocultures across the kChip with the typical single *E. coli* cell areas estimated from above. In the same way, the area of a single *C. reinhardtii* cell $A_{Sc}^{Cr}$ in monoculture/coculture was computed considering the chlorophyll clusters. The single-cell areas were independently computed for the first and the last time points.

**Computing single-cell mean intensities of *E.coli* and *C.reinhardtii* in monoculture and coculture**: Mean intensity of a single *E. coli* cell $MI_{Sc}^{Ec}$ in monoculture/coculture was estimated as the median of the mean intensities of GFP clusters in monocultures/cocultures across the kChip with the typical single *E. coli* cell areas estimated from above. In the same way, the mean intensity of a single *C.reinhardtii* cell $MI_{Sc}^{Cr}$ in monoculture/coculture was computed considering the chlorophyll clusters. The single-cell mean intensities were also independently computed for the first and the last time points.

Finally, the intensity of a single *E. coli*/*C. reinhardtii* cell $I_{Sc}^{Ec}/I_{Sc}^{Cr}$ in monoculture/coculture was computed by multiplying $A_{Sc}^{Ec}/A_{Sc}^{Cr}$ with $MI_{Sc}^{Ec}/MI_{Sc}^{Cr}$ obtained in the respective culture types.

$$I_{Sc}^{Ec-mono} = MI_{Sc}^{Ec-mono} \times A_{Sc}^{Ec-mono} \tag{S1}$$

$$I_{Sc}^{Ec-co} = MI_{Sc}^{Ec-co} \times A_{Sc}^{Ec-co} \tag{S2}$$

$$I_{Sc}^{Cr-mono} = MI_{Sc}^{Cr-mono} \times A_{Sc}^{Cr-mono} \tag{S3}$$

$$I_{Sc}^{Cr-co} = MI_{Sc}^{Cr-co} \times A_{Sc}^{Cr-co} \tag{S4}$$

The estimated values of the single-cell intensities of *E. coli* and *C. reinhardtii* at the first and the last timepoint are shown in (Fig. S3). We note that the median intensity of an *E. coli* cell computed this way showed a reduction in the single-cell intensity of *E. coli* from the first timepoint to the last timepoint by more than 50% in most cases. Whereas the single-cell intensity of a *C,reinhardtii* cell was more comparable between the time points, except in the case of galactose and acetate. Additionally, *E. coli* also showed a difference in the single-cell intensity between monoculture and coculture unlike *C. reinhardtii.*

**6.3.3.1 Error in single-cell intensity estimates** The standard error in the single-cell intensity estimates, $\delta I_{Sc}$, in monoculture/coculture at any timepoint is calculated as follows:

$$\delta I_{Sc} = I_{Sc} \times \sqrt{\left(\frac{\delta MI_{Sc}}{MI_{Sc}}\right)^2 + \left(\frac{\delta A_{Sc}}{A_{Sc}}\right)^2} \tag{S5}$$

(S6)

where $\delta MI_{Sc}$ and $\delta A_{Sc}$ represent the standard errors in the mean fluorescence intensity and the area of the single cells of the respective microbes at the corresponding time points and in the corresponding culture types (monoculture/coculture).

##### 6.3.4 Obtaining abundances of algae and bacteria

We computed the abundances of the microbes in the wells by dividing their total fluorescence intensity by the appropriate single-cell intensity estimate. For example, if the community in a well was found to be a coculture community, the abundance of *E. coli* in the well was computed by dividing the total sum of the background-subtracted fluorescence intensities of the GFP clusters in the well by the single-cell intensity of *E. coli* estimated for coculture. The mathematical expressions for calculating abundances in each of the cases are given below.

Abundance of *E. coli* $N^{Ec}$ in a well having *E. coli* monoculture community with $n$ GFP clusters,

$$I^{GFP}_{well} = \sum_{i=1}^{n} I^{GFP}_{i}, \quad N^{Ec} = \frac{I^{GFP}_{well}}{I^{Ec-mono}_{Sc}} \tag{S7}$$

where $I^{GFP}_{i}$ represents the total GFP intensity of an $i^{th}$ GFP cluster in the well and $I^{GFP}_{well}$ represents the total GFP intensity of all the GFP clusters in the well.

Abundance of *C. reinhardtii* $N^{Cr}$ in a well having *C. reinhardtii* monoculture with m chlorophyll clusters,

$$I^{Chl}_{well} = \sum_{i=1}^{m} I^{Chl}_{i}, \quad N^{Cr} = \frac{I^{Chl}_{well}}{I^{Cr-mono}_{Sc}} \tag{S8}$$

where $I^{Chl}_{i}$ represents the total chlorophyll intensity of an $i^{th}$ chlorophyll cluster in the well and $I^{Chl}_{well}$ represents the total chlorophyll intensity of all the clusters in the well.

Abundance of *E. coli* $N^{Ec}$ and abundance of *C. reinhardtii* $N^{Cr}$ in a well having *E. coli*-*C. reinhardtii* coculture with $n$ GFP clusters and $m$ chlorophyll clusters,

$$I^{GFP}_{well} = \sum_{i=1}^{n} I^{GFP}_{i}, \quad N^{Ec} = \frac{I^{GFP}_{well}}{I^{Ec-co}_{Sc}} \tag{S9}$$

$$I^{Chl}_{well} = \sum_{j=1}^{m} I^{Chl}_{i}, \quad N^{Cr} = \frac{I^{Chl}_{well}}{I^{Cr-co}_{Sc}} \tag{S10}$$

where $I^{GFP}_{i}$ and $I^{Chl}_{j}$ are respectively the total GFP and total chlorophyll intensity of an $i^{th}$ GFP and $j^{th}$ chlorophyll cluster in the well, and $I^{GFP}_{well}$ and $I^{Chl}_{well}$ are respectively the total GFP and the total chlorophyll intensity of all clusters in the well.

The growth of *E. coli*/*C. reinhardtii*, represented by $Y^{Ec}$ and $Y^{Cr}$ respectively in any well, is then obtained by subtracting the initial abundances of the microbes in the well at t = 0 h from their final abundances in the well at t = 68 h.

$$Y^{Ec} = N^{Ec}_{(t=68h)} - N^{Ec}_{(t=0h)} \tag{S11}$$

$$Y^{Cr} = N^{Cr}_{(t=68h)} - N^{Cr}_{(t=0h)} \tag{S12}$$

### 7 Determining initial pH and buffering capacity of the environments on the kChip

#### 7.1 Model to predict titration curves

To infer the pH and buffering capacity of all the barcoded environments and the environments formed by merging of the droplets on kChip, we developed a model to predict the pH titration curve of any environment given the concentrations of the nutrients and buffers in it.

A solution's buffering capacity is its resilience to pH change from additional acid or base. To characterize the buffering behavior of a defined media, we calculate the titration curve, which relates the change in pH of a solution to additions of strong acid or base.

Consider a medium consists of $a$ molar $K_2HPO_4$, $b$ molar $KH_2PO_4$, $c$ molar Tris, $d$ molar MOPS, and $e$ molar $NH_4Cl$, titrated by HCl. We denote the quantity of acid ($HCl$) added by $x$. The objective is to calculate pH as a function of $x$, given by

$$[H_2PO_4^-] + [HPO_4^{2-}] = a + b \tag{S13}$$

$$[Tris] + [TrisH^+] = c \tag{S14}$$

$$[MOPS] + [MOPS^-] = d \tag{S15}$$

$$[NH_3] + [NH_4^+] = e \tag{S16}$$

$$\frac{[HPO_4^{2-}][H^+]}{[H_2PO_4^-]} = K_1 \tag{S17}$$

$$\frac{[Tris][H^+]}{[TrisH^+]} = K_2 \tag{S18}$$

$$\frac{[MOPS^-][H^+]}{[MOPS]} = K_3 \tag{S19}$$

$$\frac{[NH_3][H^+]}{[NH_4^+]} = K_4 \tag{S20}$$

$$[H^+][OH^-] = K_w \tag{S21}$$

$$2a + b + [TrisH^+] + [H^+] + [NH_4^+] = 2[HPO_4^{2-}] + [HPO_4^-] + x + [OH^-] + [MOPS^-] + d \tag{S22}$$

Equations S13-S16 are atom conservation. Equations S17-S21 are chemical equilibrium, where $K_1, K_2, K_3, K_4$ are equilibrium constants between weak acids ($H_2PO_4^-$, $TrisH^+$, MOPS, $NH_4^+$ and their conjugate bases ($HPO_4^{2-}$, Tris, $MOPS^-$, $NH_3$), and $K_w$ is the equilibrium constant of water. Equation S21 is the charge conservation. Solving the equation gives

$$x = \frac{[H^+]}{K_1 + [H^+]}a - \frac{K_1}{K_1 + [H^+]}b + \frac{[H^+]}{K_2 + [H^+]}c - \frac{K_3}{K_3 + [H^+]}d + [H^+] - \frac{K_w}{[H^+]} - \frac{K_4}{K_4 + [H^+]}e \tag{S23}$$

Acetate and pyruvate have buffering effects too when used as carbon sources. When $f$ mol of sodium pyruvate having equilibrium constant of $K_5$ is present, the titration curve is calculated similarly:

$$x = \frac{[H^+]}{K_1 + [H^+]}a - \frac{K_1}{K_1 + [H^+]}b + \frac{[H^+]}{K_2 + [H^+]}c - \frac{K_3}{K_3 + [H^+]}d + [H^+] - \frac{K_w}{[H^+]} - \frac{K_4}{K_4 + [H^+]}e + \frac{[H^+]}{K_5 + [H^+]}f \tag{S24}$$

When $g$ mol of sodium acetate having equilibrium constant of $k_6$ is present, the titration curve is calculated similarly:

$$x = \frac{[H^+]}{K_1 + [H^+]}a - \frac{K_1}{K_1 + [H^+]}b + \frac{[H^+]}{K_2 + [H^+]}c - \frac{K_3}{K_3 + [H^+]}d + [H^+] - \frac{K_w}{[H^+]} - \frac{K_4}{K_4 + [H^+]}e + \frac{[H^+]}{K_6 + [H^+]}g \tag{S25}$$

Table S4 shows chemical constants for all buffering agents in the experiment. Note that the equilibrium constants depend on temperature. Because equilibrium constants in the literature are often measured at $25°C$, we calculate the corrected equilibrium constant at the experimental temperatures ($30°C$) using the following equation [12]:

$$pK_T = pK_\theta - \frac{1}{R\ln 10}\left[\Delta H(\frac{1}{\theta} - \frac{1}{T}) + \Delta C_p\left(\frac{\theta}{T} - 1 + \ln\frac{T}{\theta}\right)\right]$$

Here $pK$ is defined as $pK = -\log_{10} K$. $\theta$ denotes the reference temperature ($25°C$) and $T$ denote the temperature of interest ($30°C$). $R$ is the ideal gas constant. $\Delta H$ is the ionization enthalpy and $\Delta C_p$ is the ionization thermal capacity at constant pressure [12].

#### 7.2 Computing initial pH and buffering capacity

The initial pH of an environment is obtained by computationally solving the equations S23/S24/S25 at $x = 0$, depending upon the carbon source. For our purpose, we define buffering capacity as the quantity of HCl that drops the pH to a point just before the pH can abruptly change with [HCl]. Hence, we compute buffering capacity as the smallest $x$ (from equations S23/S24/S25) where the change in the first derivative before the inflection point on the pH curve is just greater than 30 (pH/[HCl] units). This method yields buffering capacity values that agree with our definition as shown in several examples (Fig. S4). We expect that our measure of buffering capacity determines the allowed acidification in the environment before the pH drops to very low values at which the microbial growth will be negatively impacted [8, 13, 14]. Using these definitions of initial pH and buffering capacity, we were able to compute the initial pH and buffering capacity of all the environments formed by the merging of the droplets on kChip from the model. We note that only in the case of acetate, the buffering capacity was evaluated without taking acetate into consideration. As the $pK_a$ of acetate is $\sim$4.98, the titration curve of the environments having acetate do not have the abrupt drop in the pH with an increase in [HCl] as in the examples shown in Fig. S4. However, as the buffering capacity values computed for environments without acetate correspond to low values of pH ($\sim$<6) where the microbial growth is negatively affected, using these buffering capacity values for environments with acetate agrees with our definition of buffering capacity and hence should be valid.

##### 7.2.1 Correcting the initial pH

We experimentally validated our titration model for a set of environmental conditions given in Table S2 (Fig. S4). We found that the predicted initial pH was in good agreement with the experimental data in cases where [MOPS] was $\sim$10 mM (Fig. S5A). And the predicted initial pH deviated from the observed initial pH in cases where [MOPS]<10 mM, the deviations being high at low concentrations of Tris and at low values of experimentally observed pH. The conditions with low observed pH were also the conditions with very low buffering capacity. We speculate that the low buffering capacity could be making the environment susceptible to pH changes (from uncharacterized chemicals in the water source or atmospheric gases) and causing poor agreement between the model and data.

We corrected the deviation in the predicted initial pH using linear regressions (Fig. S5B). As can be observed, the qualitative nature of the disagreement between the initial pH values predicted from the model and the initial pH values obtained from the experiments, in environments with no MOPS buffer and with MOPS at ∼5 mM, differed. Hence, two separate linear regressions were set up, one to correct the data with no MOPS and another to correct the data with ∼5 mM MOPS. Using these regression models, the predicted initial pH values of all the environments formed on the kChip were corrected appropriately.

### 8 Linear regression analyses

#### 8.1 Model formulation

Linear models were set up to predict the growth of microbes $Y$ from the environmental factors - initial pH ($pH$), buffering capacity ($BC$), phosphorus concentration ($[P]$), and carbon concentration ($[C]$). As discussed in the main text, the model was formulated as follows:

$$Y = \vec{\beta}_M \begin{bmatrix} 1 \\ [P] \\ [C] \\ pH[P] \\ pH[C] \\ BC[P] \\ BC[C] \end{bmatrix} + I\vec{\beta}_I \begin{bmatrix} 1 \\ [P] \\ [C] \\ pH[P] \\ pH[C] \\ BC[P] \\ BC[C] \end{bmatrix} + \beta_A A \tag{S26}$$

The variable $I$ is the indicator variable that is 0 for all monoculture wells and 1 for co-culture wells. $A$ represents the area of the merged droplets in the well at 68 h, as inferred by the fluorescent dye images. The area feature is included to account for the differences in the merged droplet volumes across communities. The area feature is not included with the indicator variable $I$ as we do not expect any difference in the contribution of area to the microbial growth in monoculture and coculture. The $\vec{\beta}_M$ and $\vec{\beta}_I$ denote the vectors of monoculture and interaction coefficients for the corresponding features, and $\beta_A$ represents the coefficient of the area feature $A$.

$$\vec{\beta}_M = [\beta_{1,M}, \beta_{[P],M}, \beta_{[C],M}, \beta_{pH[P],M}, \beta_{pH[C],M}, \beta_{BC[P],M}, \beta_{BC[C],M}, \beta_{[P][C],M}] \tag{S27}$$

$$\vec{\beta}_I = [\beta_{1,I}, \beta_{[P],I}, \beta_{[C],I}, \beta_{pH[P],I}, \beta_{pH[C],I}, \beta_{BC[P],I}, \beta_{BC[C],I}, \beta_{[P][C],I}] \tag{S28}$$

For each carbon source, two such regression models were set up, one for predicting the growth of *E. coli* $Y^{Ec}$ and another for predicting the growth of *C. reinhardtii* $Y^{Cr}$.

#### 8.2 Data preprocessing

The growth data from the kChip experiments was preprocessed for the regression modelling to facilitate the interpretation of the regression results, as indicated in the main text.

Firstly, we classified the growth data into the different culture types based on the following scheme:

1. The data with $N^{Ec}_{(t=0h)} > 0.5$ and $N^{Cr}_{(t=0h)} < 0.2$ were classified as *E. coli* monocultures

2. The data with $N^{Ec}_{(t=0h)} < 0.2$ and $N^{Cr}_{(t=0h)} > 0.5$ were classified as *C. reinhardtii* monocultures
3. The data with $N^{Ec}_{(t=0h)} > 0.5$ and $N^{Cr}_{(t=0h)} > 0.5$ were classified as *E. coli* - *C. reinhardtii* cocultures

Following this, the data with merged-droplets area of the community between 850 pixels and 2000 pixels at 68 h were retained (The median merged-droplets area of the communities at 68 h were in the range of 1100-1400 pixels for the different carbon sources). The discarding of the data with merged-droplets area outside of 850 pixels and 2000 pixels at 68 h removed wells that have undergone excessive evaporation. Then again, as it is not feasible to examine these large datasets one by one to remove those with imaging artifacts, stray fluorescence signals, imperfect wells on the microfluidic chip etc, the data was again filtered to account for any extreme outliers. In the case of monoculture data, the highest and the lowest 0.05% of the growth data of the microbes considering all the monoculture wells were discarded. In the case of coculture data, wells with *C. reinhardtii* growth in the highest and the lowest 0.05% of the *C. reinhardtii* growth and with *E. coli* growth in the highest 2% and lowest 0.05% of the *E. coli* growth considering all the coculture wells were discarded. This method of discarding the data ensured that the growth of the microbes in the discarded data lay well beyond the lowest and the highest median growth of the microbes across replicate environmental conditions. Overall, the above filtering schemes led to data losses of ∼4.8%, ∼4.5%, ∼3.4%, ∼8.1%, and ∼4.2% in the glycerol, glucose, galactose, pyruvate, and acetate datasets respectively.

Following the removal of outliers, the yields of each microbe (*E. coli*/*C. reinhardtii*) within its culture types (monoculture/coculture) were independently Z-score normalized. That is, for each microbe within its culture type (*E. coli* monoculture/*C. reinhardtii* monoculture/*E. coli* coculture/*C. reinhardtii* coculture), the mean and standard deviation of the growth $Y$ were computed and all of the growth data was subtracted from the mean and then divided by the standard deviation to obtain the standardized growth values.

The values of the independent variables - initial pH ($pH$), buffering capacity ($BC$), carbon concentration ($[C]$), and area ($A$) were also independently transformed to range from 0 to 1 for each carbon source. Only phosphorus concentrations ($[P]$) were first log-transformed (owing to the order-of-magnitude variation in the phoshporus concentrations across environments) and then scaled to range from 0 to 1 independently for each carbon source. This scaling brought the values of the independent variables to similar ranges, avoiding the domination of a variable with the highest magnitude in training the regression model.

#### 8.3 Implementing the regressions

A weighted least squares approach was used to solve for the coefficients in equation S26. The weighted least squares approach optimizes the cost function to find the regression coefficients by accounting for the variability in the number of data points across environments (e.g. number of wells with the same environment and culture type). In our case, the weighted least squares approach works by weighting the squared error of the data by 1/variance in growth across its replicates. Replicates here refer to wells with the same culture type (*E. coli* monoculture/*C. reinhardtii* monoculture/*E. coli*-*C. reinhardtii* coculture) and environmental condition. Consider $z$ environments indexed by $j$. In each environment, we have $n_j^{mono}$ replicate wells having monocultures and $n_j^{co}$ replicate wells having cocultures. Within each environment, we compute a variance across monoculture replicates $\sigma^2(Y^j_{data,m})$ where $Y^j_{data,m}$ is the growth in well $m$ having monoculture that contains environment $j$, and variance across coculture replicates $\sigma^2(Y^j_{data,c})$

where $Y^j_{data,c}$ is the growth in well $c$ having coculture that contains environment $j$. We then optimize the following objective function:

$$\sum_{j=1}^{z}\left(\sum_{i=1}^{n_j^{mono}}\frac{1}{\sigma^2(Y^j_{data,m})}(Y^j_{data,m}-\hat{Y}^j_{model,m})^2+\sum_{i=1}^{n_j^{co}}\frac{1}{\sigma^2(Y^j_{data,c})}(Y^j_{data,c}-\hat{Y}^j_{model,c})^2\right) \tag{S29}$$

(S30)

Using the WLS function in the statsmodels package in python, two regression models, one for predicting *E. coli* growth and another for predicting *C. reinhardtii* growth, were fitted to the standardized growth data of the respective microbes in each of the wells. The fits obtained from the regressions with the pearson coefficients and the RMSE values are shown in Fig. S9. And the $\beta$ coefficients obtained for the regression models are shown in Fig. S10. The 95% confidence intervals and the p-values of the coefficients reported here were obtained from the summary output of the regressions in python.

#### 8.4 Computing coculture coefficients

As discussed in the main text, the monoculture coefficient $\beta_{X,M}$ and the interaction coefficient $\beta_{X,I}$ where $X \in ([P],[C],pH[P],pH[C],BC[P],BC[C],[P][C])$, respectively indicate the change in the growth in monoculture per unit change in X and the change in growth per unit change in X in coculture relative to monoculture. Hence, the coculture coefficient $\beta_{X,C}$, representing the change in growth in coculture per unit change in X, can be obtained by adding $\beta_{X,M}$ and $\beta_{X,I}$. And the 95% confidence interval of a coculture coefficient $\beta_{X,C}$ was computed using the covariance matrix of the features (obtained from regression analyses output in Python) as follows:

$$\beta_{X,C} = 1.96\left(\sqrt{var(\beta_{X,M})+var(\beta_{X,I})+2cov(\beta_{X,M},\beta_{X,I})}\right) \tag{S31}$$

The p-values of the coculture coefficients were obtained as outlined in [15].

### 9 Hierarchical clustering of carbon sources

Two Hierarchical clusterings were performed (using Scipy (cluster.hierarchy) with Ward's distance as the linkage metric) to find similarities between the carbon sources based on

1. similarities in the microbial growth (Fig. 6C, main text)

2. similarities in the regression coefficients (Fig. 6B, main text)

We began by constructing the data matrices for hierarchical clusterings. In the case of (1), carbon sources formed the columns, and environmental conditions in the different culture types for each microbe type (*E. coli* monoculture/*C. reinhardtii* monoculture/*E. coli* coculture/*C. reinhardtii* coculture) formed the rows, with matrix entries the median standardized growth $Y$ of *E. coli* or *C. reinhardtii* mapping to the environmental conditions, culture types, and microbe types.

In the case of (2), carbon sources formed the columns again and features with and without the indicator variable $I$ i.e ($[P]$,$[C]$, $pH[P]$, $pH[C]$, $BC[P]$, $BC[C]$,$[P][C]$, $I[P]$,$I[C]$, $IpH[P]$, $IpH[C]$, $IBC[P]$, $IBC[C]$,$I[P][C]$) in the regression models of *E. coli* and *C. reinhardtii* growth formed the rows, with matrix entries the monoculture or interaction coefficients obtained from regressing *E. coli* growth, or monoculture or interaction coefficients obtained from regressing *C. reinhardtii* growth i.e $\beta_M^{Ec}$ or $\beta_I^{Ec}$ or $\beta_M^{Cr}$ or $\beta_I^{Cr}$, mapping to the features and the microbe type.

The correlation matrix for hierarchical clusterings was computed accounting for the error in the data. If $v_k$ represents the column vectors of the data matrices where $k \in (1, 2, 3, 4, 5)$ represents the five carbon sources, we compute the following quantities to arrive at the correlation coefficient between $v_k$ and $v_l$:

1. **Weighted mean of $v_k$ and $v_l$:**

$$\mu_{v_k} = \frac{\sum_p \frac{1}{\sigma^2(v_{kp})} v_{kp}}{\sum_p \frac{1}{\sigma^2(v_{kp})}}, \quad \mu_{v_l} = \frac{\sum_p \frac{1}{\sigma^2(v_{lp})} v_{lp}}{\sum_p \frac{1}{\sigma^2(v_{lp})}} \tag{S32}$$

   where $v_{kp}$ and $v_{lp}$ represent the $p^{th}$ entry in $v_k$ and $v_l$ respectively, and $\sigma(v_{lp})$ and $\sigma(v_{kp})$ represent the standard errors/95% confidence interval (as appropriate), in the $p^{th}$ entry in $v_k$ and $v_l$ respectively

2. **Weighted covariance between $v_k$ and $v_l$:**

$$cov(v_k, v_l) = \frac{\sum_p w_p (v_{kp} - \mu_{v_k})(v_{lp} - \mu_{v_l})}{\sum_p w_p} \tag{S33}$$

$$\tag{S34}$$

   where $\quad w_p = \frac{1}{\sigma^2(v_{kp}) + \sigma^2(v_{lp})}$

Then $corr(v_k, v_l)$, the correlation between the carbon sources $k$ and $l$ in $v$ is computed as:

$$corr(v_k, v_l) = \frac{cov(v_k, v_l)}{\sqrt{cov(v_k, v_k) cov(v_l, v_l)}} \tag{S35}$$

$$\tag{S36}$$

The correlations between all pairs of carbon sources are computed using the same formulae.

### 10 Plate experiment assaying *E. coli* growth on carbon sources

The growth rate of *E. coli* measured in microtiter plates in the five carbon sources are reported in Table S5. For this experiment, the bacterial culture was grown and prepared as described in section 4. The growth was assayed in the low pH, low buffering capacity media conditions formed by combining the environments E3 and E4 and in the low pH, high buffering capacity media condition formed by combining the environments E11 and E12 (Table S2), via continuous measurement of OD590 for $\tilde{6}8$h using the Tecan infinite F200 PRO plate reader. The growth rate was inferred by fitting a straight line to the linear portion of the natural logarithm of OD in time. The initial pH and buffering capacities of the environments (predicted by the titration model described in section 7), along with the growth rates and OD590 (at 68 h) of *E. coli* and the final pH (at 68 h) of the cultures (measured using VWR pH paper BDH35309.606) are reported in Table S5. The final pH was measured to be different in the two environments in the case of glucose and glycerol but similar between the respective environments in the two carbon sources - the pH drop is higher in E3+E4 environment which has lower buffering capacity than the environment E11+E12 with higher buffering capacity. On the other hand, the final pH is similar in both E3+E4 and E11+E12 in the case of galactose, pyruvate, and acetate.

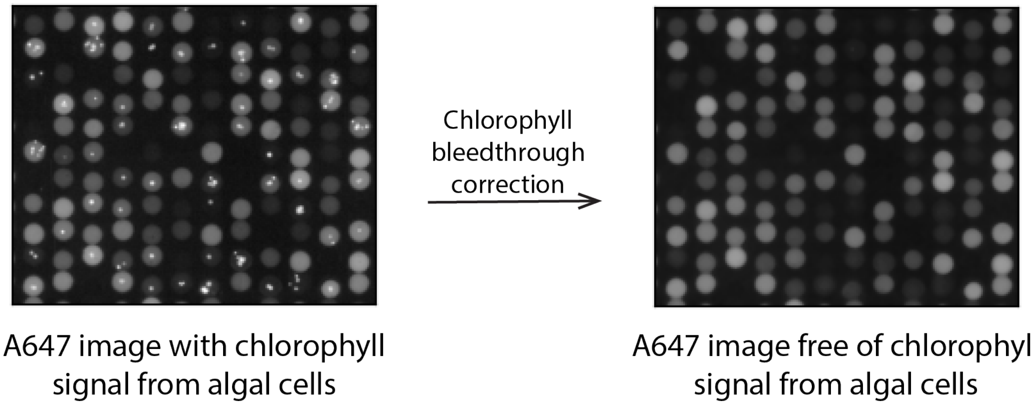

**Figure S1: Correcting the Alexa Fluor 647 image for chlorophyll bleed-through.** On the left, an example image of the droplets on kChip acquired with the filter used for imaging the Alexa Fluor 647 dye. The pixel intensity values have been log transformed to show both the dimmer Alexa Fluor 647 dye signal in the droplets and the brighter chlorophyll signal (the tiny white spots) from the algal cells. On the right, the corresponding chlorophyll-corrected image of the Alexa Fluor 647 dye image, free of the chlorophyll signal from the algal cells, after processing the image as described in Section 6.1.

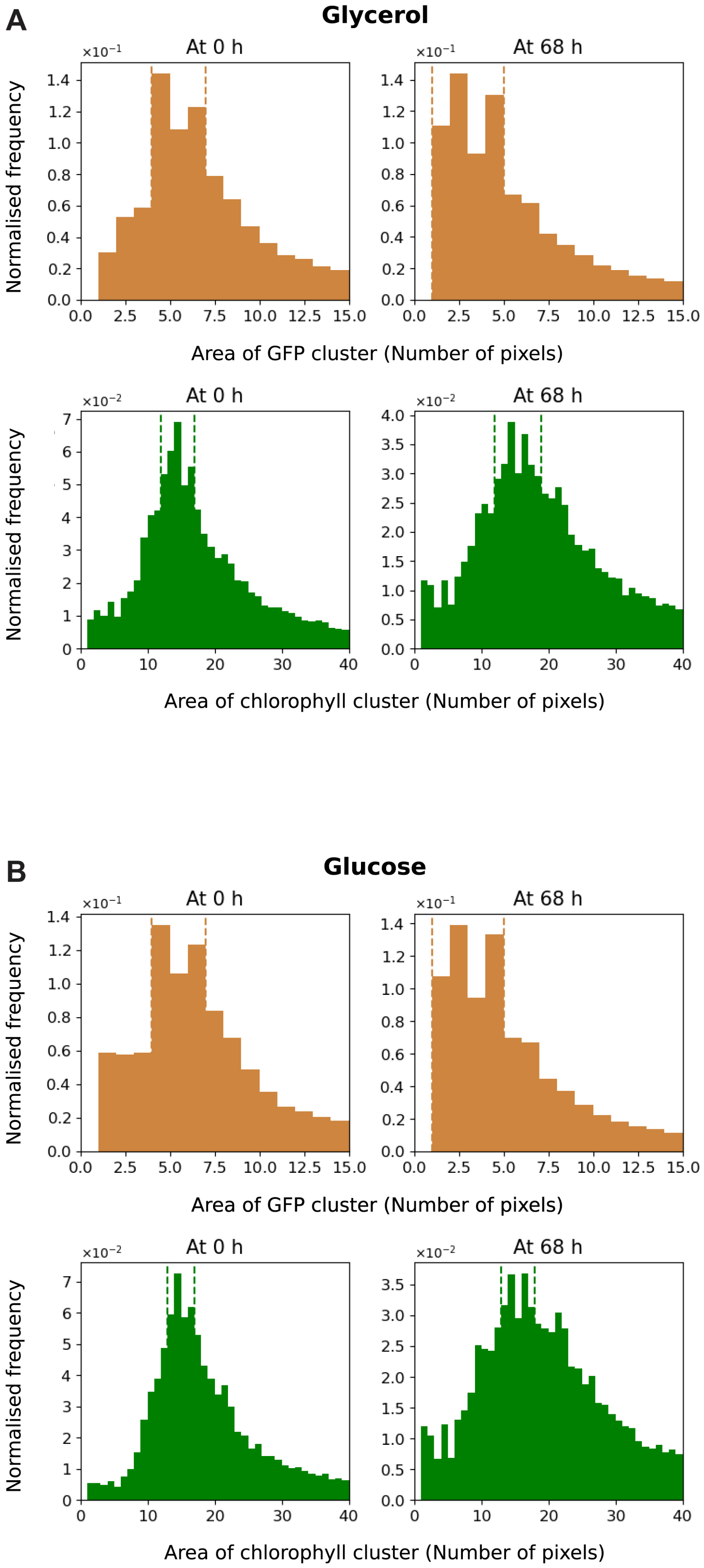
A
Glycerol
At 0 h
At 68 h
Normalised frequency
Area of GFP cluster (Number of pixels)
Area of chlorophyll cluster (Number of pixels)
B
Glucose
At 0 h
At 68 h
Normalised frequency
Area of GFP cluster (Number of pixels)
Area of chlorophyll cluster (Number of pixels)

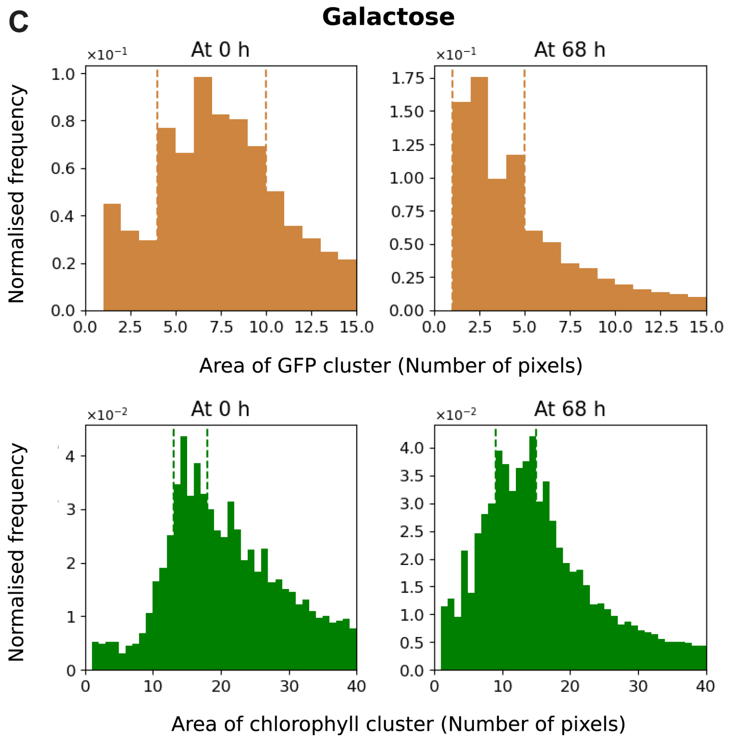
C
Galactose
At 0 h
At 68 h
Normalised frequency
Area of GFP cluster (Number of pixels)
At 0 h
At 68 h
Normalised frequency
Area of chlorophyll cluster (Number of pixels)

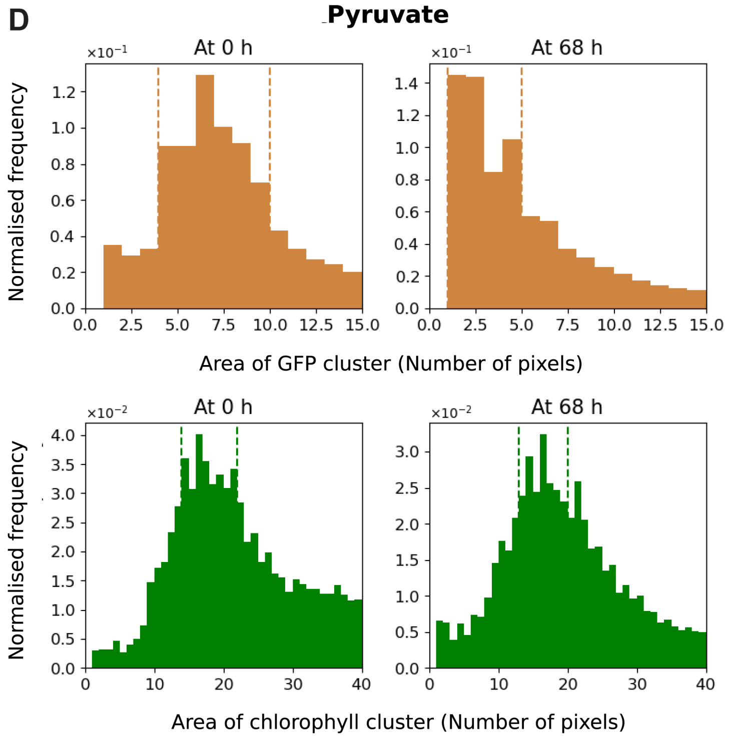
D
Pyruvate
At 0 h
At 68 h
Normalised frequency
Area of GFP cluster (Number of pixels)
At 0 h
At 68 h
Normalised frequency
Area of chlorophyll cluster (Number of pixels)

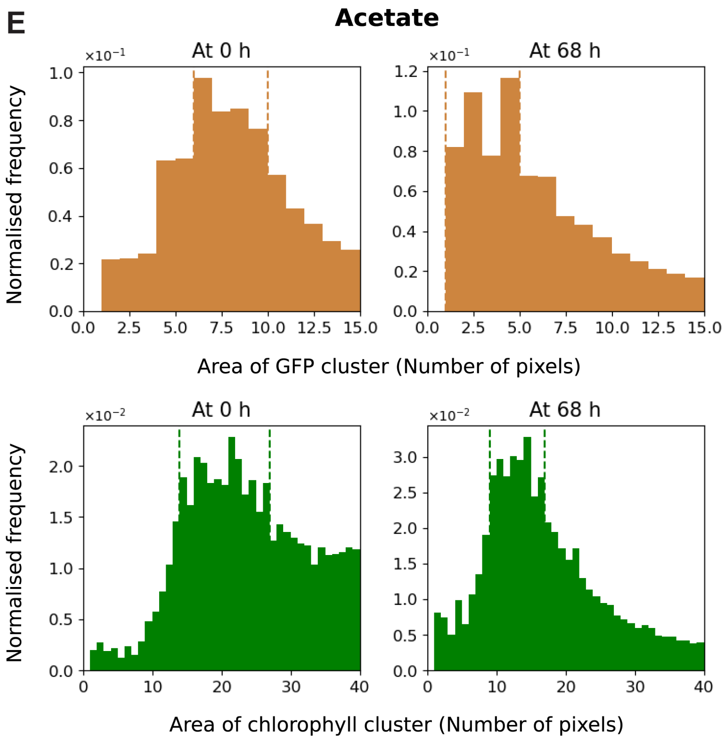

**Figure S2: Inferring typical areas of single-cells of *E. coli* and *C. reinhardtii*.** Distribution of the areas (in units of pixels) of GFP and chlorophyll clusters (in brown and green respectively) across the kChip, at 0 h (plots on the left) and 68 h (plots on the right) in glucose in panel (A), glycerol in panel (B), galactose in panel (C), pyruvate in panel (D), and acetate in panel (E). The dashed lines mark the chosen bounds for the typical single-cell areas of *E. coli* and *C. reinhardtii* at 0 h and 68 h.

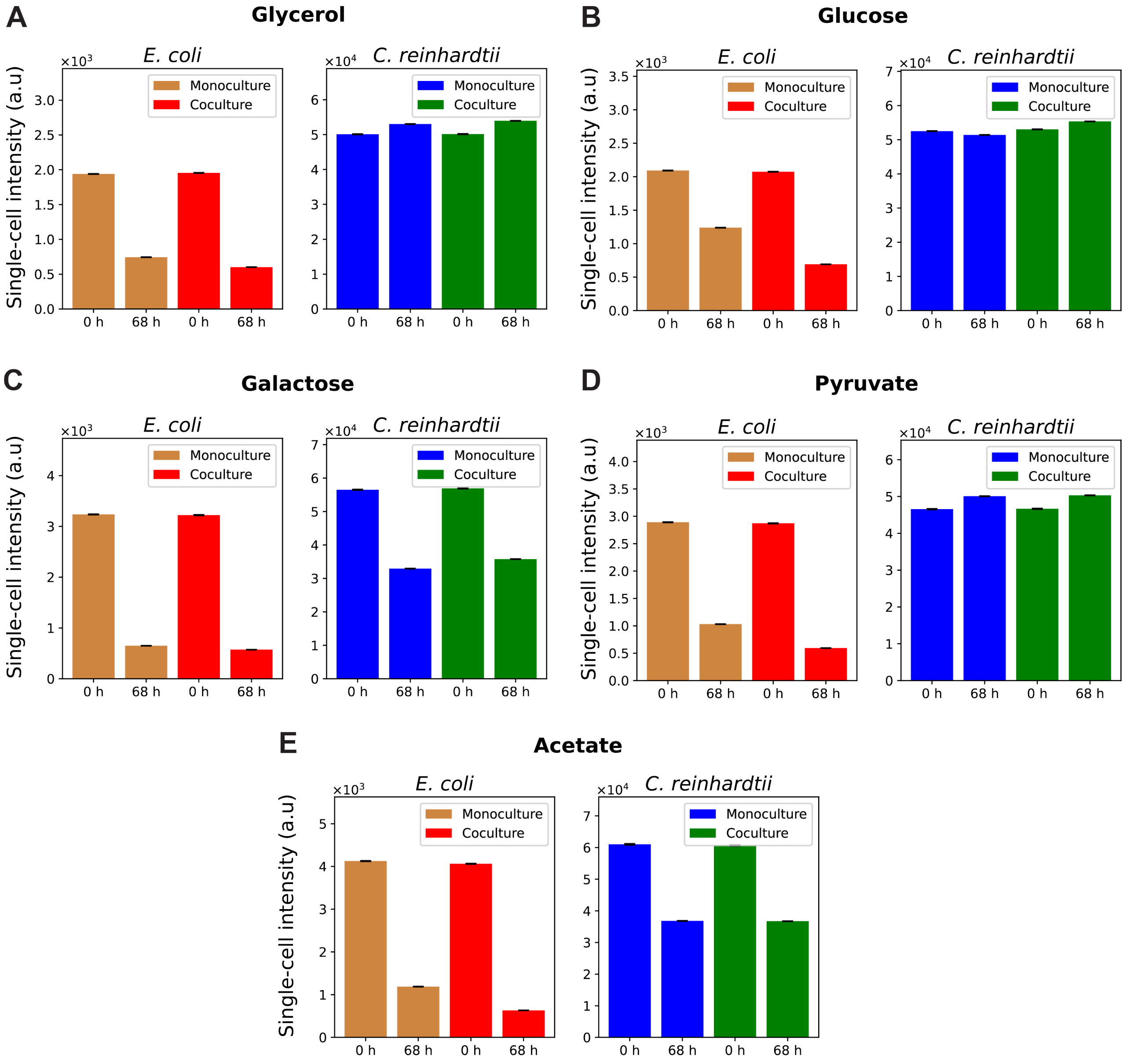

**Figure S3: Single-cell intensity estimates for *E. coli* and *C. reinhardtii*.** Single-cell intensity of *E.coli* and *C. reinhardtii* in GFP and chlorophyll pixel intensity units, estimated in monoculture and coculture at 0 h and 68 h, in glucose in panel (A), glycerol in panel (B), galactose in panel (C), pyruvate in panel (D), and acetate in panel (E). The standard errors about the mean are indicated as black bars and are typically lower than 0.01% of the single-cell intensity estimates.

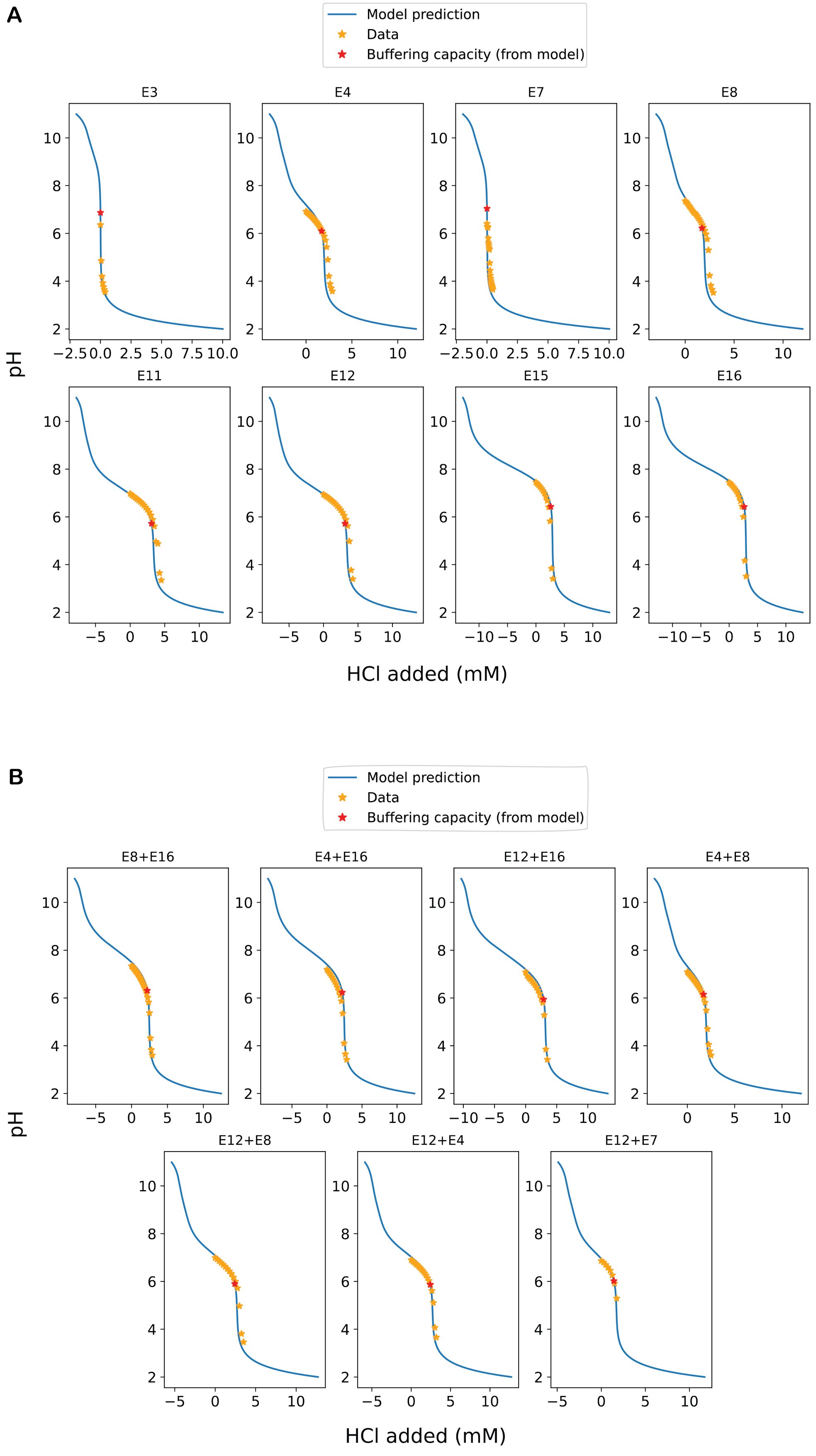

**Figure S4:** (Caption next page)

**Figure S4:** (Previous page.) **Experimental validation of the pH titration model.** The pH vs HCl data obtained from titration experiments (in orange stars) is overlaid on the prediction of the pH curve from the titration model (in solid blue line) for several of the initial 16 barcoded environments in (A) and for several environments formed by combinations of the barcoded environments in (B). The pH-HCl point for estimating the buffering capacities from the model-predicted pH curves are shown with red stars. Refer to Table S2 for details about the environmental conditions E3,E4,E7,E8,E11,E12,E15,E16.

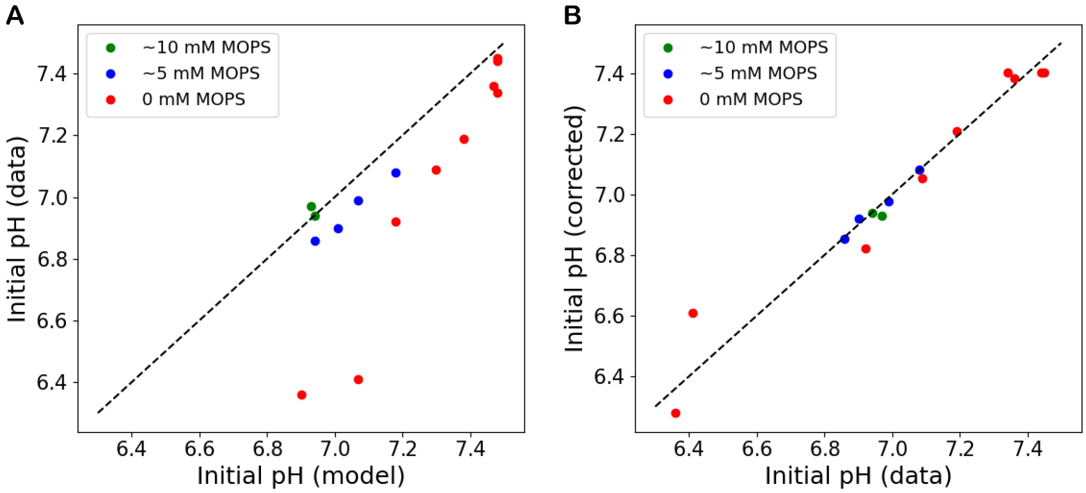

**Figure S5: Correction of the initial pH predicted from the titration model.** (A) Deviation of the model-predicted initial pH (on the x-axis) from the experimentally obtained initial pH (on the y-axis) for environments having ∼10 mM MOPS buffer (in green), ∼mM MOPS buffer (in blue) and no MOPS buffer (in red). (B) Agreement between the experimentally obtained initial pH values (x-axis) and the model-predicted initial pH values that are corrected using linear regressions (y-axis).

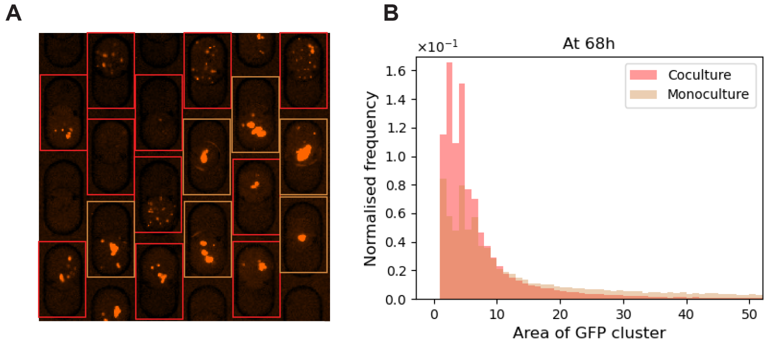

**Figure S6: Aggregated and planktonic bacterial cells in monocultures and cocultures respectively.** (A) An example GFP image showing the bacterial cells (in brown) with wells having *E. coli* monocultures bounded by brown squares and wells having *E. coli* - *C. reinhardtii* cocultures bounded by red squares. (B) Distribution of areas of GFP cluster (i.e connected region in the mask of the GFP image representing the planktonic and aggregated bacterial cells - see section 6.3.2 for details about finding area of GFP clusters) in cocultures (in red) and monocultures (in brown) across the kChip in the case of glucose at 68 h. The x-axis range has been limited to a range of areas of GFP clusters to show the lower proportion of smaller bacterial clusters (planktonic bacterial cells) in monoculture compared to in coculture at the end of the experiment at 68h.

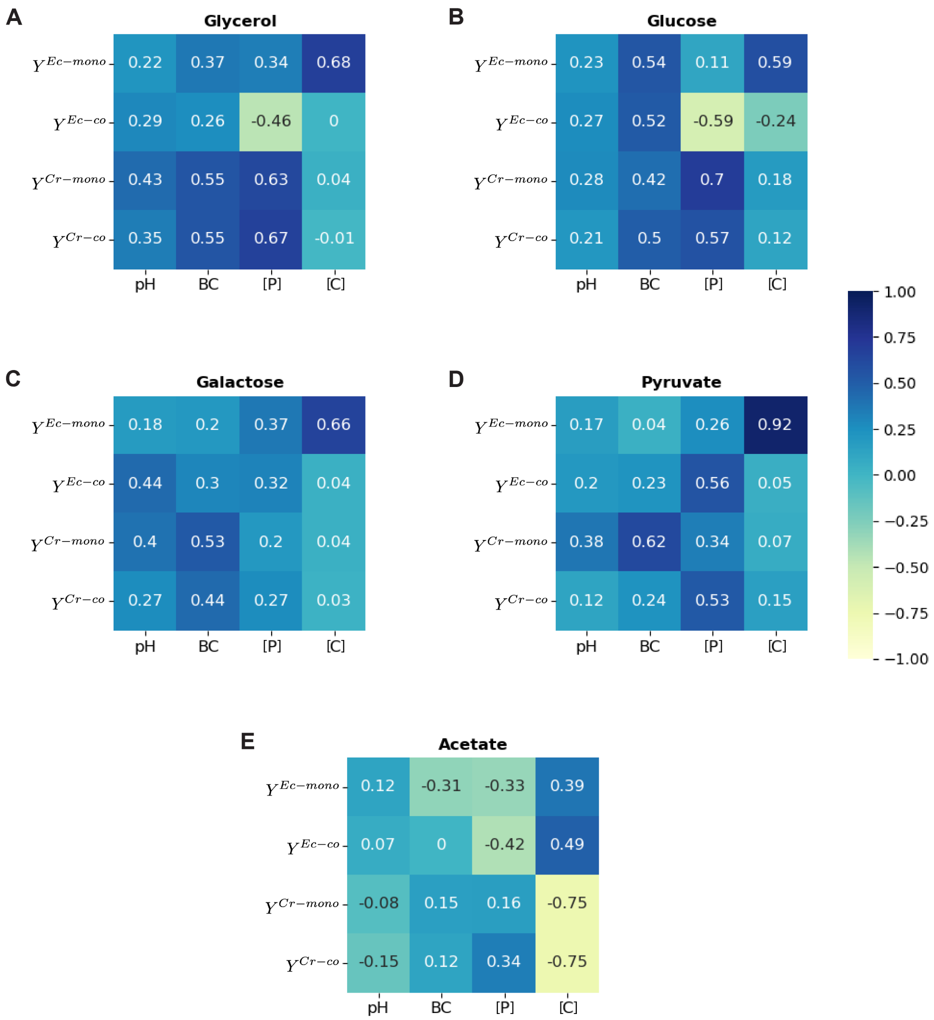

**Figure S7: Correlations between algae-bacteria growth and environmental factors - pH, buffering capacity, phosphorus concentration, and carbon concentration.** The Pearson correlation coefficients computed between the growth of *E. coli* in monoculture $Y^{Ec-mono}$, growth of *E. coli* in coculture $Y^{Ec-co}$, growth of *C. reinhardtii* in monoculture $Y^{Cr-mono}$, growth of *C. reinhardtii* in coculture $Y^{Cr-co}$ and the environmental factors - pH, buffering capacity BC, phosphorus concentration [P] and carbon concentration [C] in the case of glycerol in (A), glucose in (B), galactose in (C), pyruvate in (D) and acetate in (E)

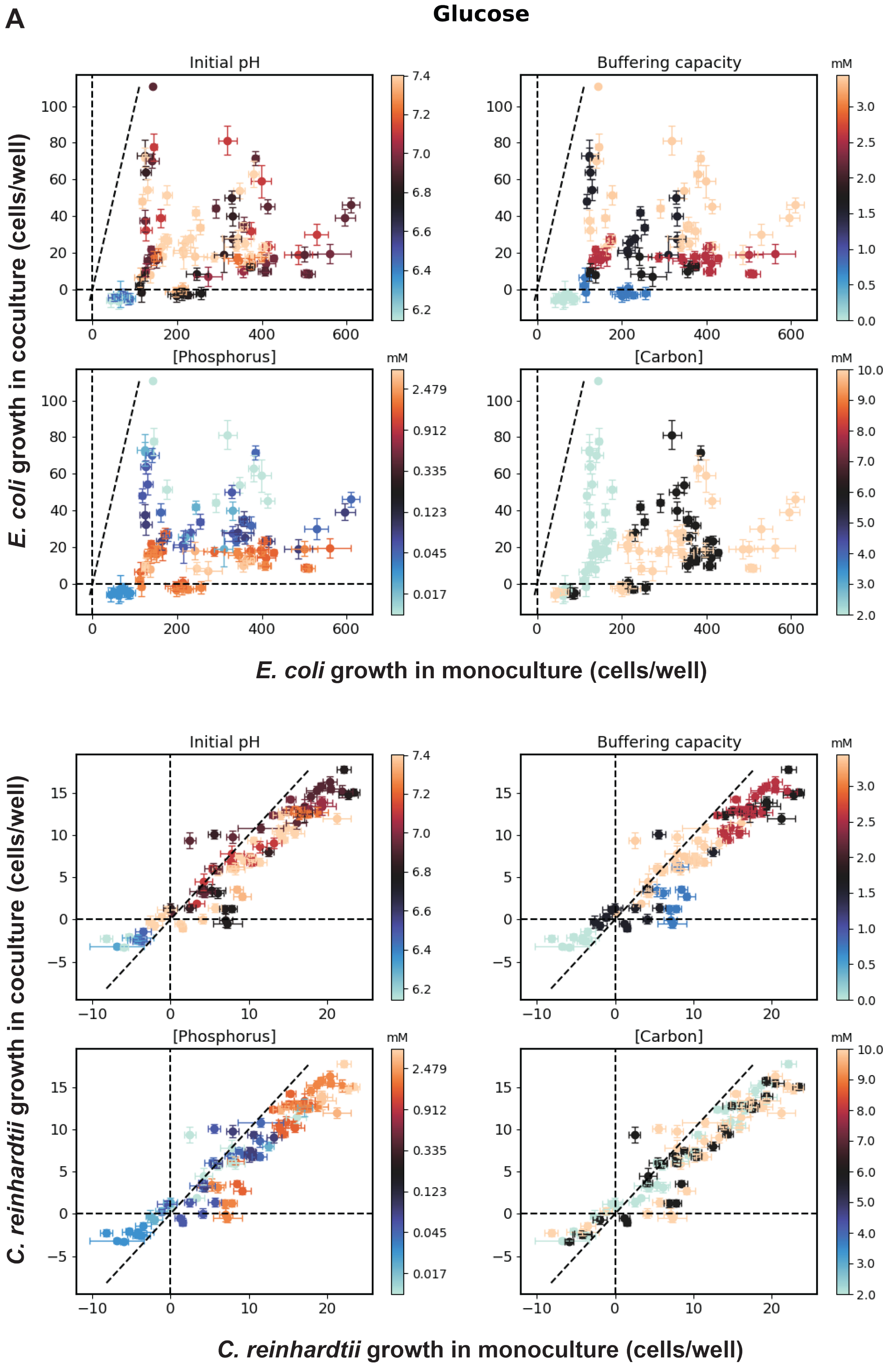
A
Glucose
Initial pH
Buffering capacity
[Phosphorus]
[Carbon]
mM
E. coli growth in coculture (cells/well)
E. coli growth in monoculture (cells/well)
C. reinhardtii growth in coculture (cells/well)
C. reinhardtii growth in monoculture (cells/well)

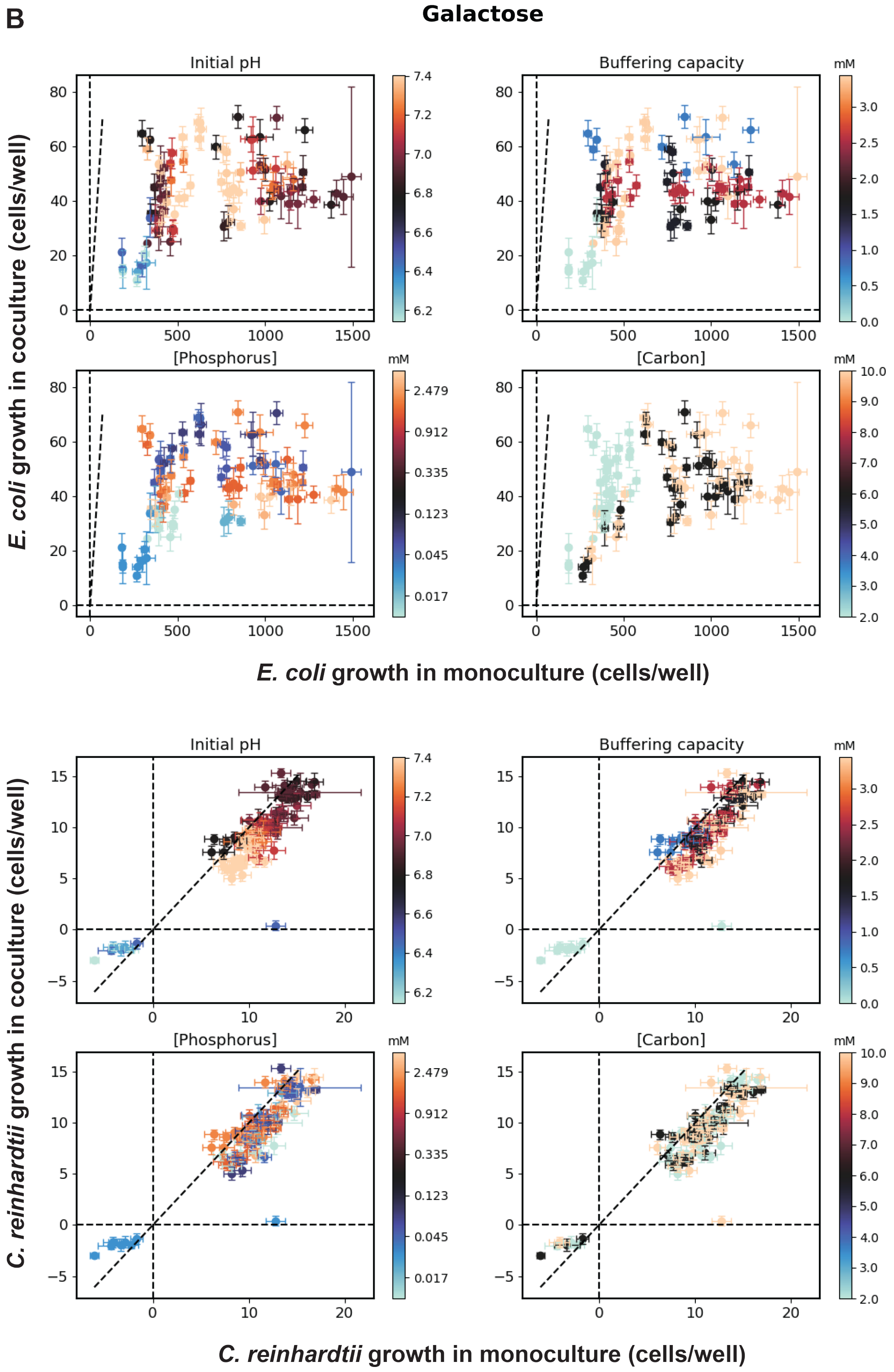

B
Galactose
Initial pH
Buffering capacity
[Phosphorus]
[Carbon]
mM
E. coli growth in coculture (cells/well)
E. coli growth in monoculture (cells/well)
Initial pH
Buffering capacity
[Phosphorus]
[Carbon]
mM
C. reinhardtii growth in coculture (cells/well)
C. reinhardtii growth in monoculture (cells/well)

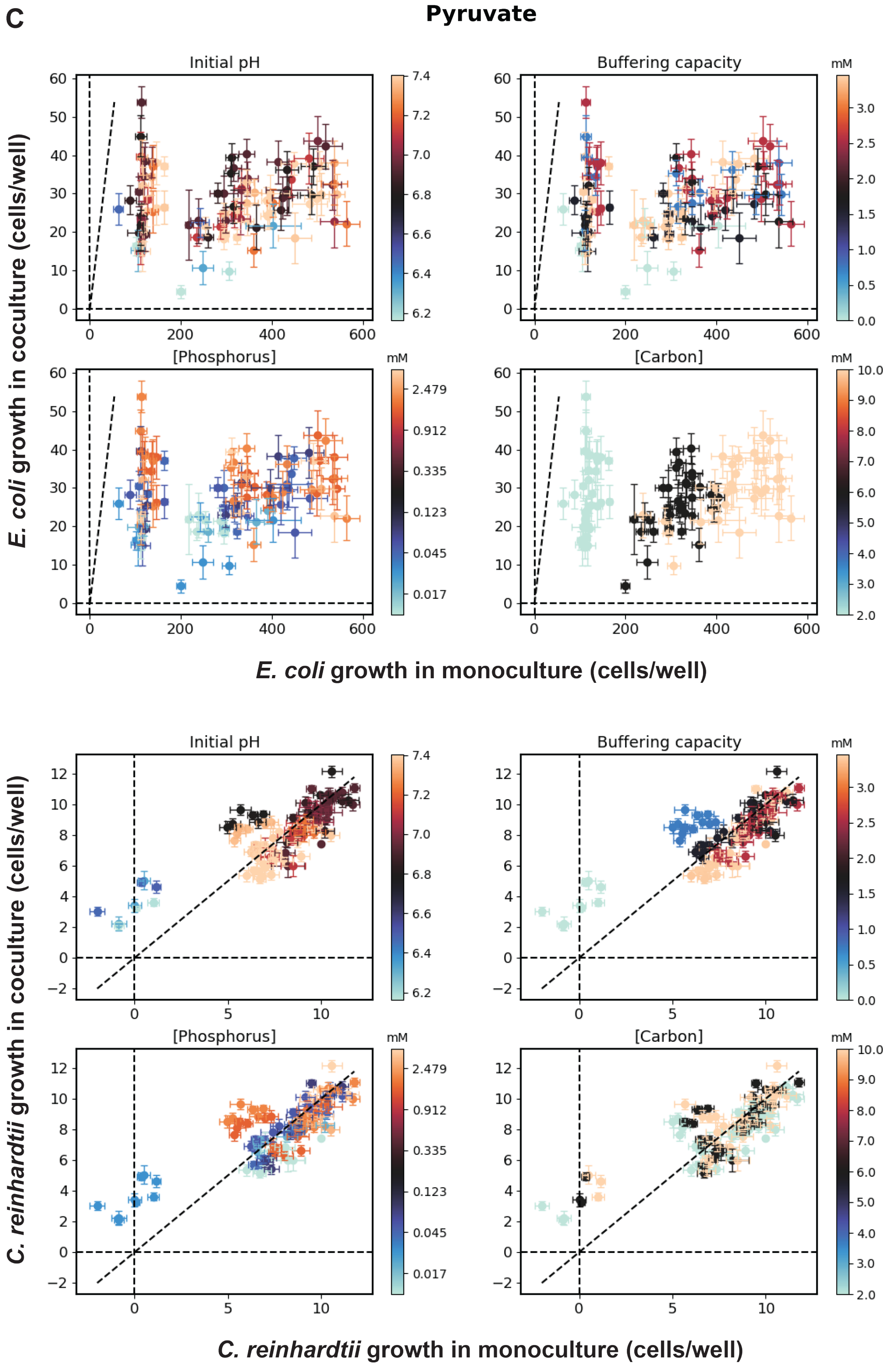
C
Pyruvate
Initial pH
Buffering capacity
[Phosphorus]
[Carbon]
mM
7.4
7.2
7.0
6.8
6.6
6.4
6.2
3.0
2.5
2.0
1.5
1.0
0.5
0.0
2.479
0.912
0.335
0.123
0.045
0.017
10.0
9.0
8.0
7.0
6.0
5.0
4.0
3.0
2.0
E. coli growth in coculture (cells/well)
E. coli growth in monoculture (cells/well)
C. reinhardtii growth in coculture (cells/well)
C. reinhardtii growth in monoculture (cells/well)

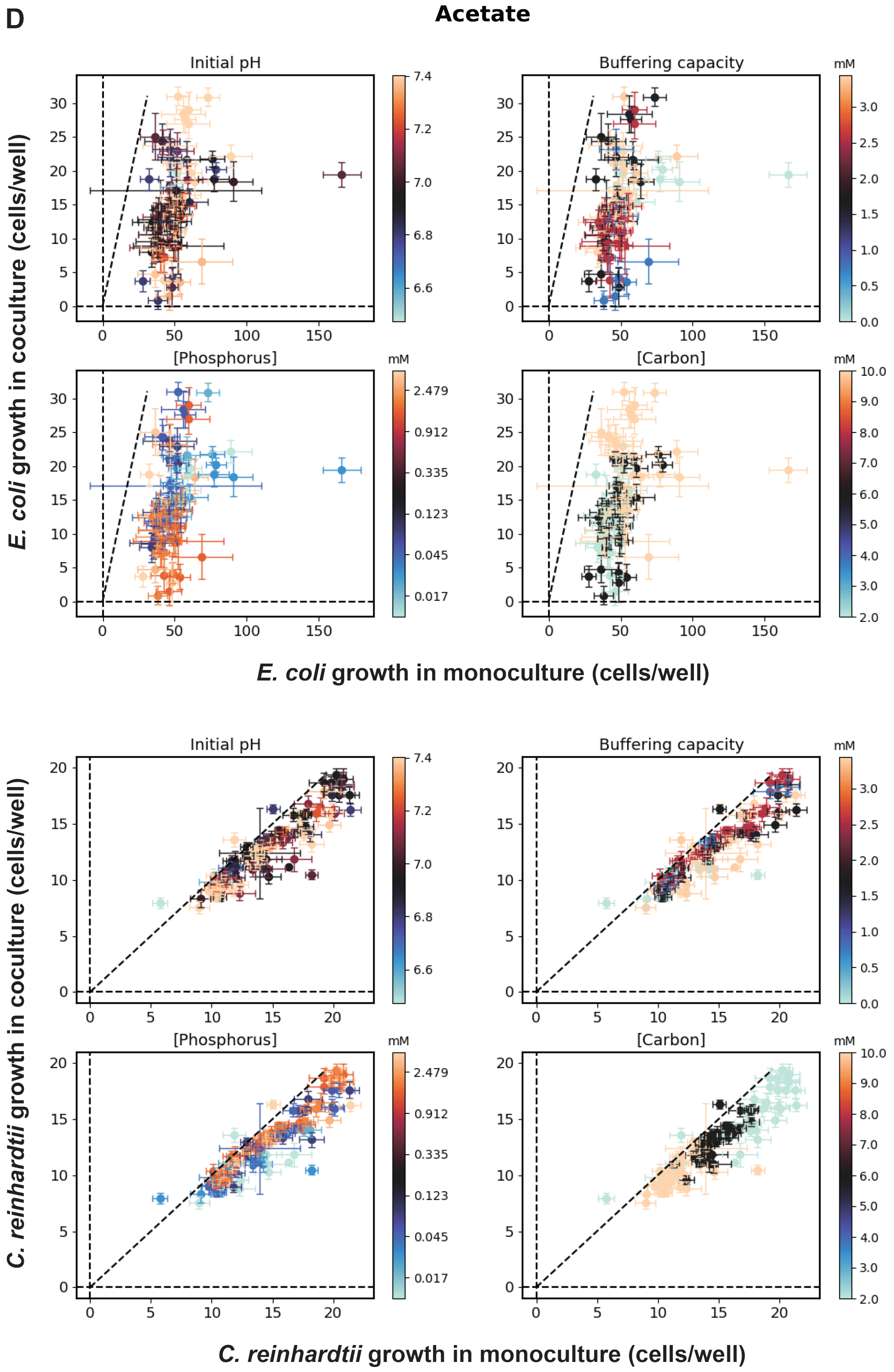

**Figure S8:** (Caption next page)

**Figure S8:** (Previous page) **Complex dependence of algae-bacteria interactions on the environmental factors.** The median growth of *E. coli* computed across coculture replicates (y-axis) for each of the environmental conditions are plotted against the median yields of *E. coli* computed across monoculture replicates (x-axis) in the respective environmental conditions in the top set of 4 panels in the case of glucose in (A), galactose in (B), pyruvate in (C), and acetate in (D). The growth of *C. reinhardtii* computed across coculture replicates (y-axis) for each of the environmental conditions are plotted against the growth of *C. reinhardtii* (x-axis) computed across monoculture replicates in the respective environmental conditions in the bottom set of 4 panels in the case of glucose in (A), galactose in (B), pyruvate in (C), and acetate in (D). The error bars indicate the standard errors about the mean growth. The colors in the different plots correspond to the values of the environmental factors - Initial pH (top left), buffering capacity (top right), phosphorus concentration (bottom left), and carbon concentration (bottom right) in the respective environmental conditions, as indicated by the corresponding color bar to the right of the plots. The values of only the phosphorus concentrations were log-transformed to generate the plot but the color bar is annotated with the original untransformed values of phosphorus concentrations.

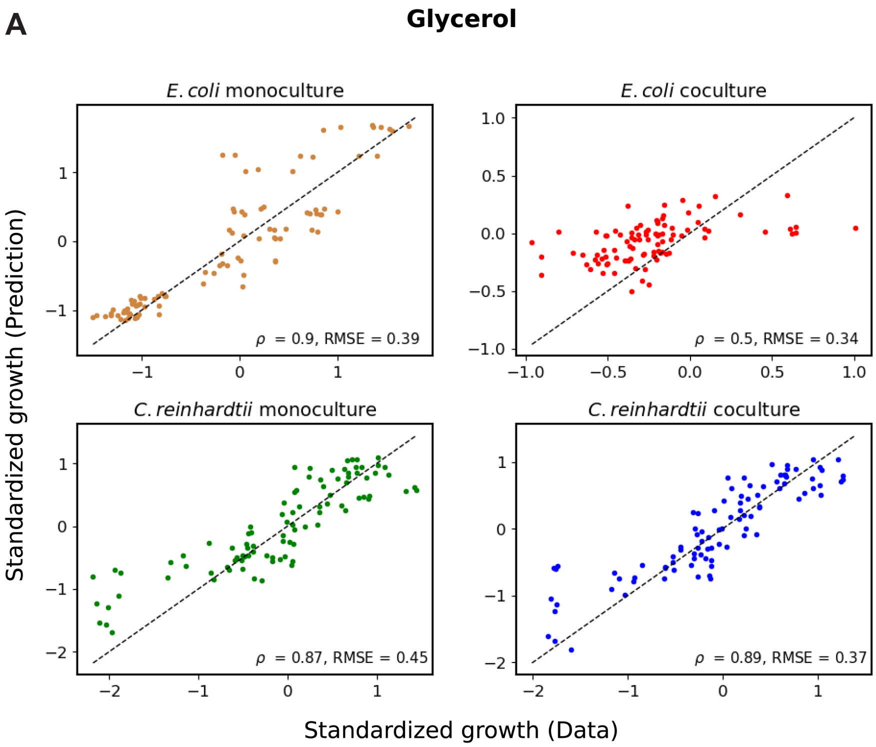

**B** **Glucose**

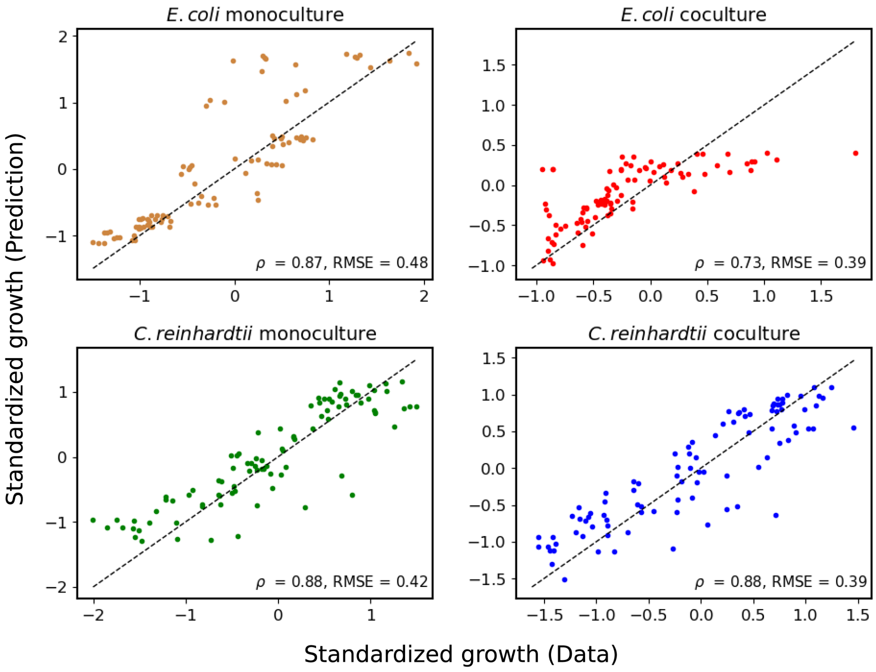

**C** **Galactose**

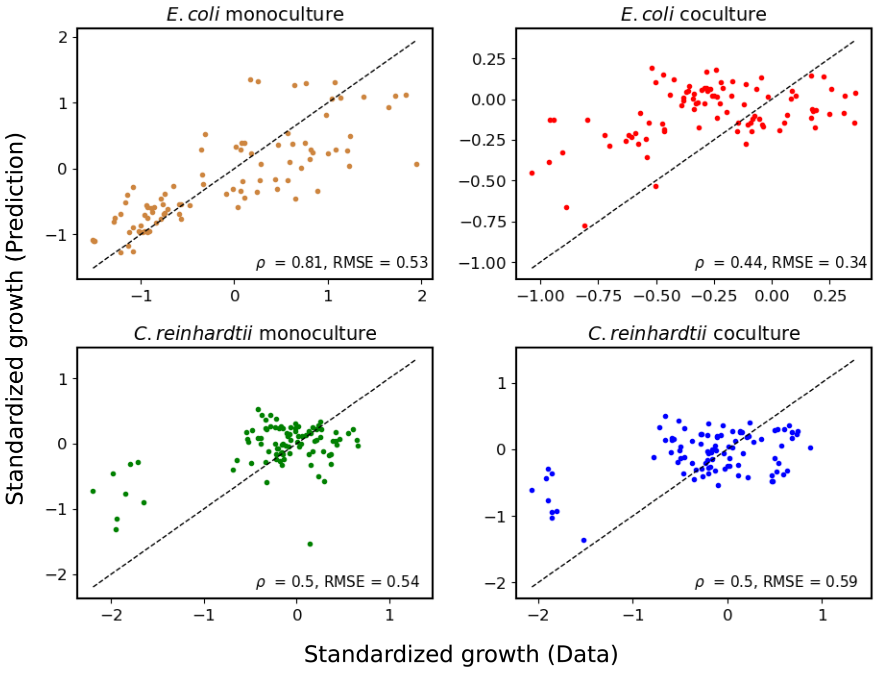

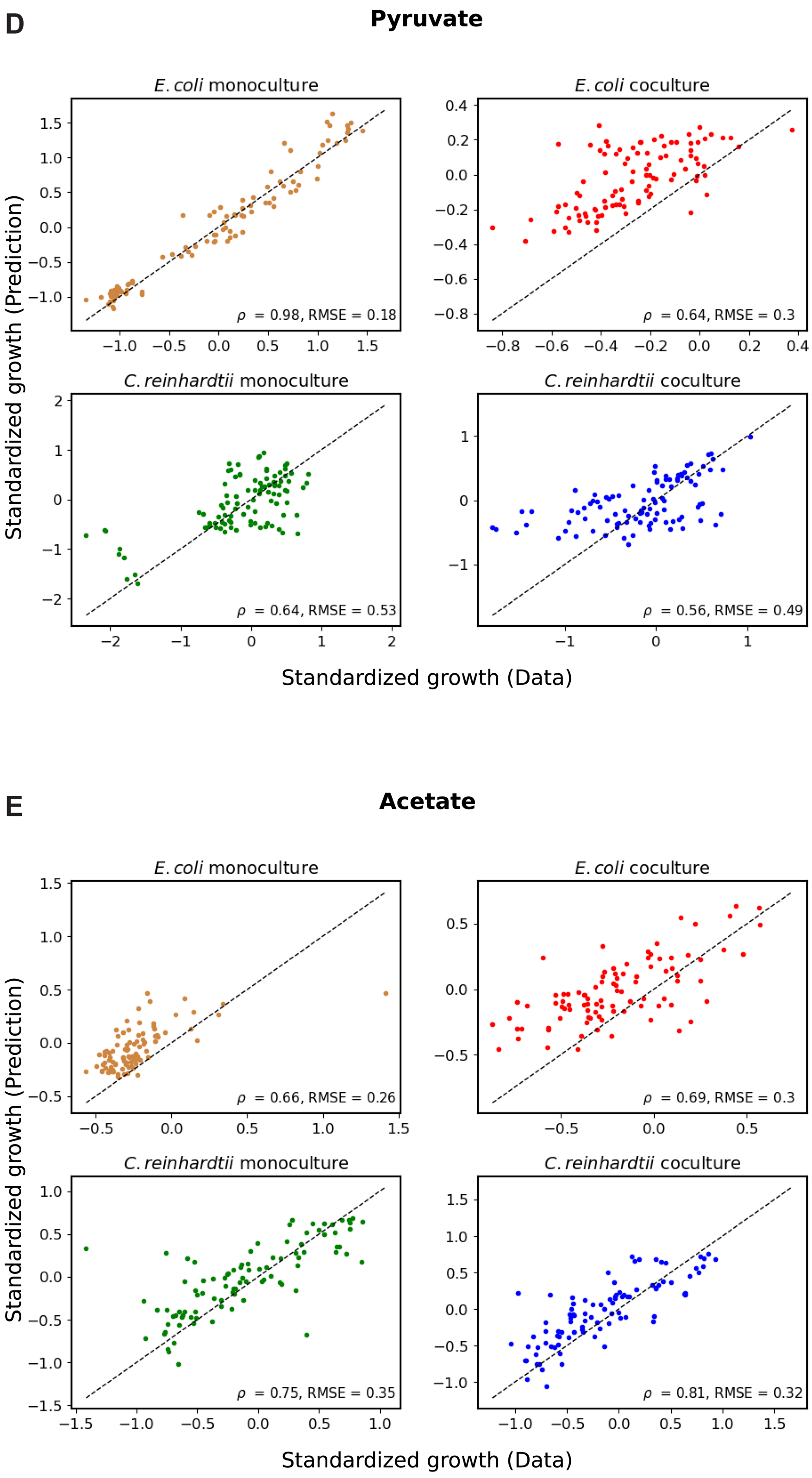

**Figure S9:** (Caption next page)

**Figure S9:** (Previous page) **Comparison between the predicted growth and the observed growth of *E. coli* and *C. reinhardtii* in the environments formed on kChip.** The observed growth was computed as the median of the standardized growth across replicate wells. The predicted growth was computed as the median of the standardized growth obtained from the regression model (Equation, S26) across replicates. See section 8.2 for details about the standardization of the growth data. The comparison between the predicted growth (y-axis) and the observed growth (x-axis) is shown for *E. coli* in monoculture in the top left panel, *E. coli* in coculture in the top right panel, *C. reinhardtii* in monoculture in the bottom left panel, and *C. reinhardtii* in coculture in the bottom right panel for all the environments in the case of glycerol in (A), glucose in (B), galactose in (C), pyruvate in (D), and acetate in (E). The Pearson correlation coefficient $\rho$ and the root mean squared error RMSE of the fits are reported in the corresponding panels.

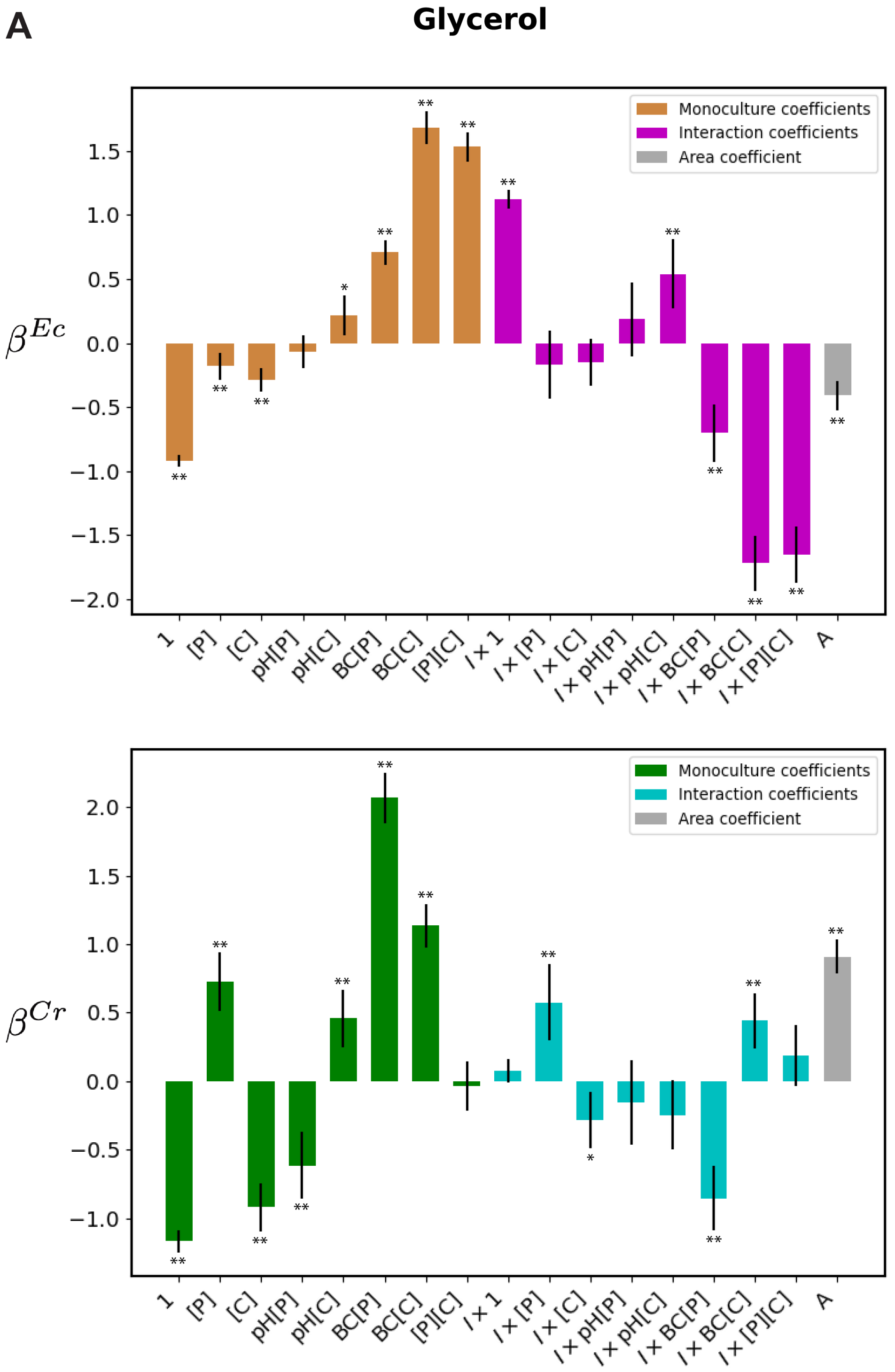
A
Glycerol
Monoculture coefficients
Interaction coefficients
Area coefficient
$\beta^{Ec}$
1
[P]
[C]
pH[P]
pH[C]
BC[P]
BC[C]
[P][C]
$I \times 1$
$I \times$ [P]
$I \times$ [C]
$I \times$ pH[P]
$I \times$ pH[C]
$I \times$ BC[P]
$I \times$ BC[C]
$I \times$ [P][C]
A
Monoculture coefficients
Interaction coefficients
Area coefficient
$\beta^{Cr}$

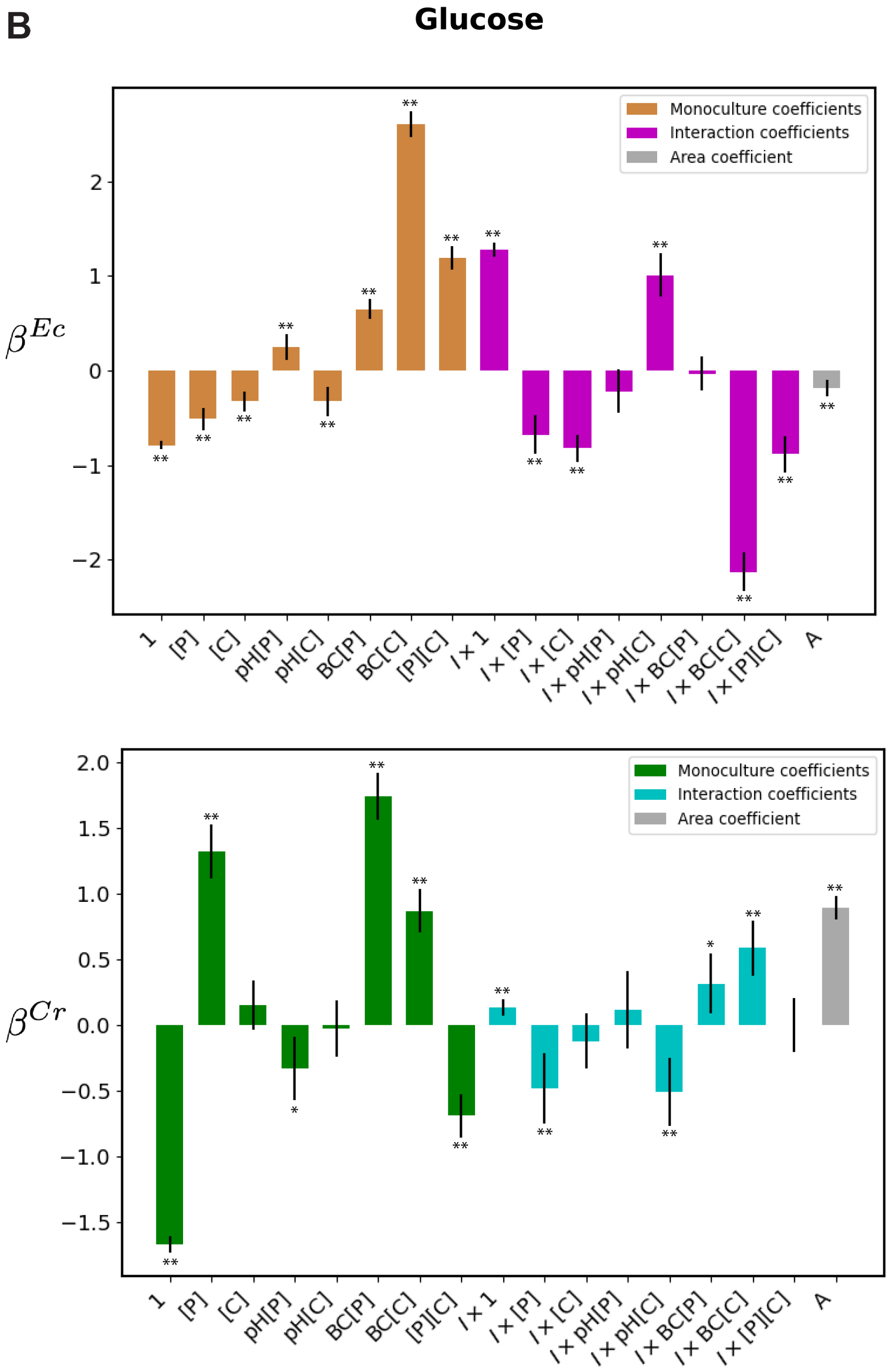
B
Glucose
Monoculture coefficients
Interaction coefficients
Area coefficient
$\beta^{Ec}$
$\beta^{Cr}$
1
[P]
[C]
pH[P]
pH[C]
BC[P]
BC[C]
[P][C]
$I \times 1$
$I \times$ [P]
$I \times$ [C]
$I \times$ pH[P]
$I \times$ pH[C]
$I \times$ BC[P]
$I \times$ BC[C]
$I \times$ [P][C]
A

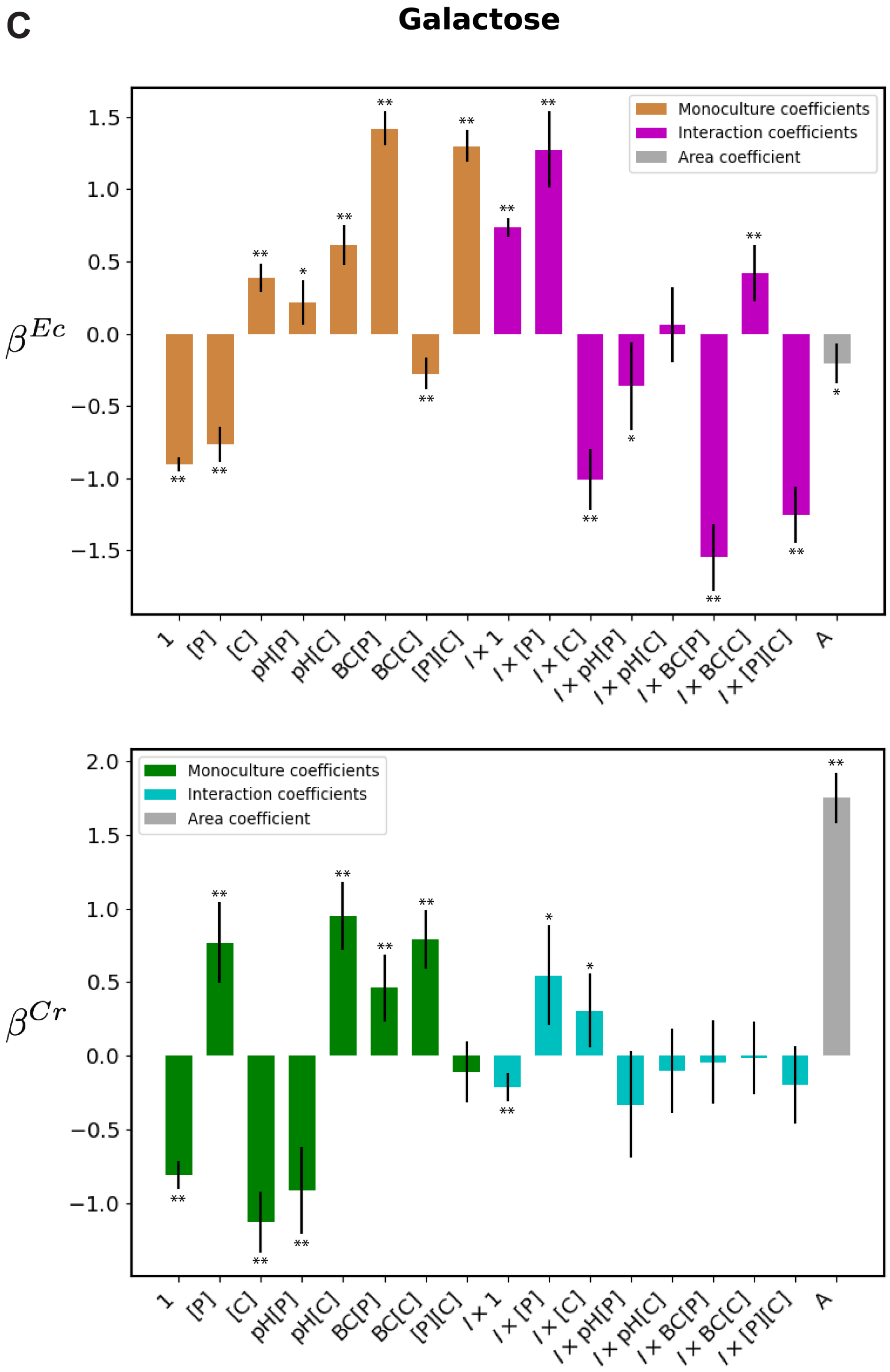
C
Galactose
Monoculture coefficients
Interaction coefficients
Area coefficient
$\beta^{Ec}$
$\beta^{Cr}$
1
[P]
[C]
pH[P]
pH[C]
BC[P]
BC[C]
[P][C]
I×1
I×[P]
I×[C]
I×pH[P]
I×pH[C]
I×BC[P]
I×BC[C]
I×[P][C]
A

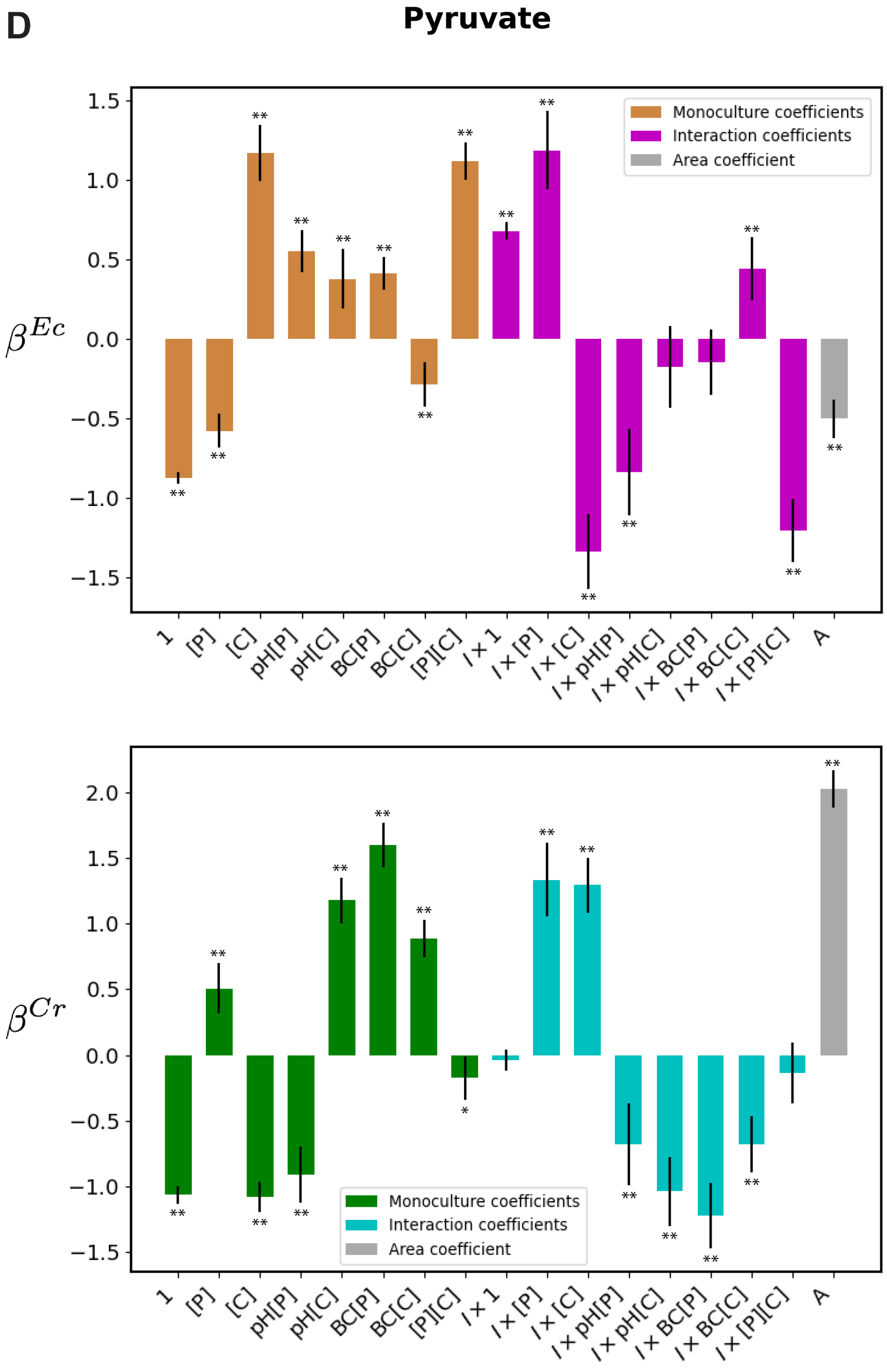
D
Pyruvate
$\beta^{Ec}$
Monoculture coefficients
Interaction coefficients
Area coefficient
1
[P]
[C]
pH[P]
pH[C]
BC[P]
BC[C]
[P][C]
$I \times 1$
$I \times$ [P]
$I \times$ [C]
$I \times$ pH[P]
$I \times$ pH[C]
$I \times$ BC[P]
$I \times$ BC[C]
$I \times$ [P][C]
A
$\beta^{Cr}$
Monoculture coefficients
Interaction coefficients
Area coefficient

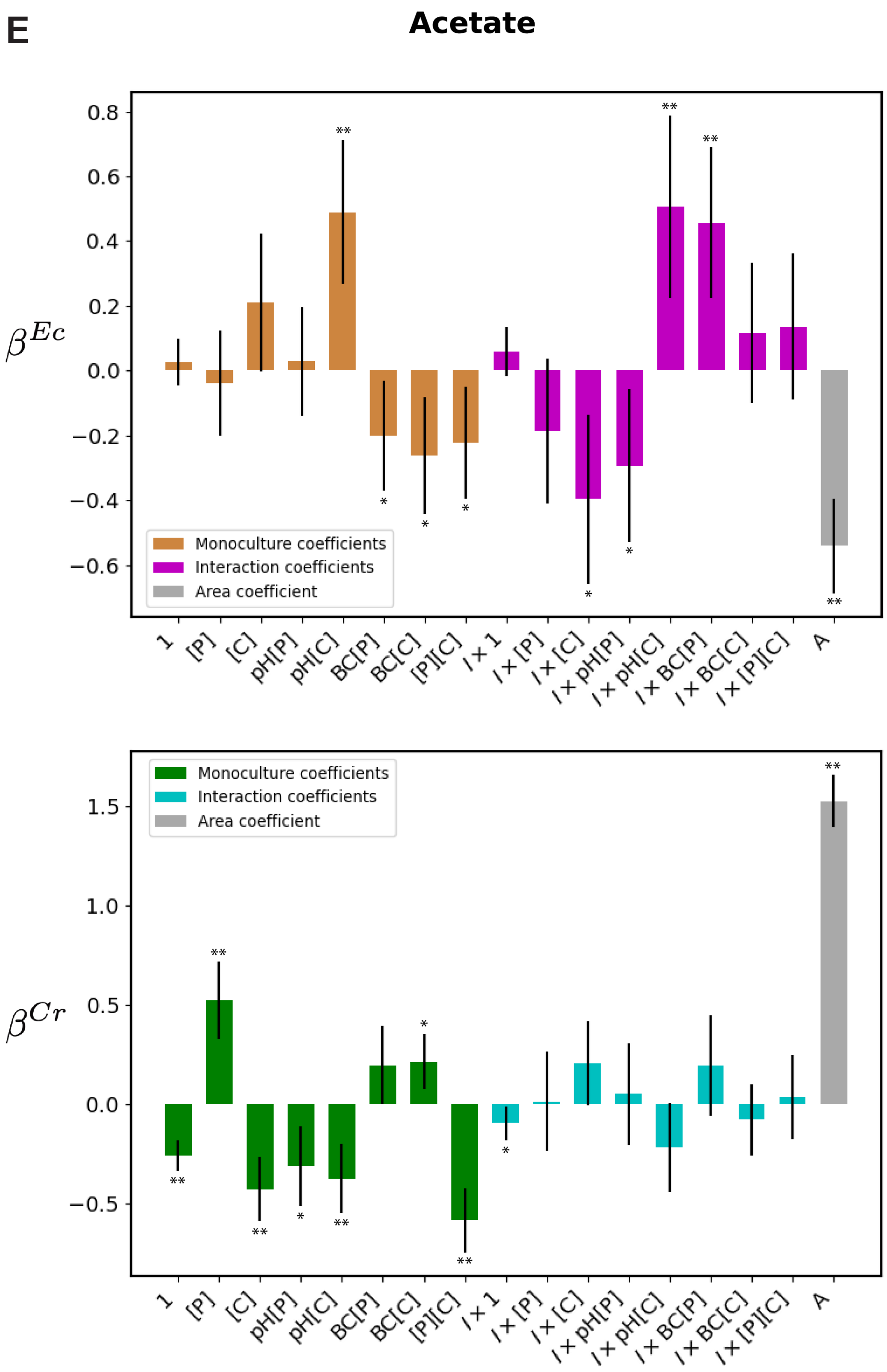

**Figure S10:** (Caption next page)

**Figure S10:** (Previous page) **The complete set of regression coefficients for models predicting growth from environmental conditions.** The top panels in (A),(B),(C),(D), and (E) report the monoculture coefficients (brown bars) and interaction coefficients (magenta bars) of the corresponding features on the x-axis, and the coefficient of the area feature A obtained from the regression model (Equation. S26) predicting the growth of *E. coli* in monocultures and cocultures in the case of glycerol in (A), glucose in (B), galactose in (C), pyruvate in (D), and acetate in (E). Similarly, bottom panels in (A),(B),(C),(D), and (E) report the monoculture coefficients (green bars) and interaction coefficients (cyan bars) of the corresponding features on the x-axis, and the coefficient of the area feature A obtained from the regression model predicting the growth of *C. reinhardtii* in monocultures and cocultures in the case of glycerol in (A), glucose in (B), galactose in (C), pyruvate in (D), and acetate in (E). The error bars represent the 95% confidence intervals. The statistical significance of the results is indicated with **(p-value<0.001) and *(p-value<0.05) atop the bars.

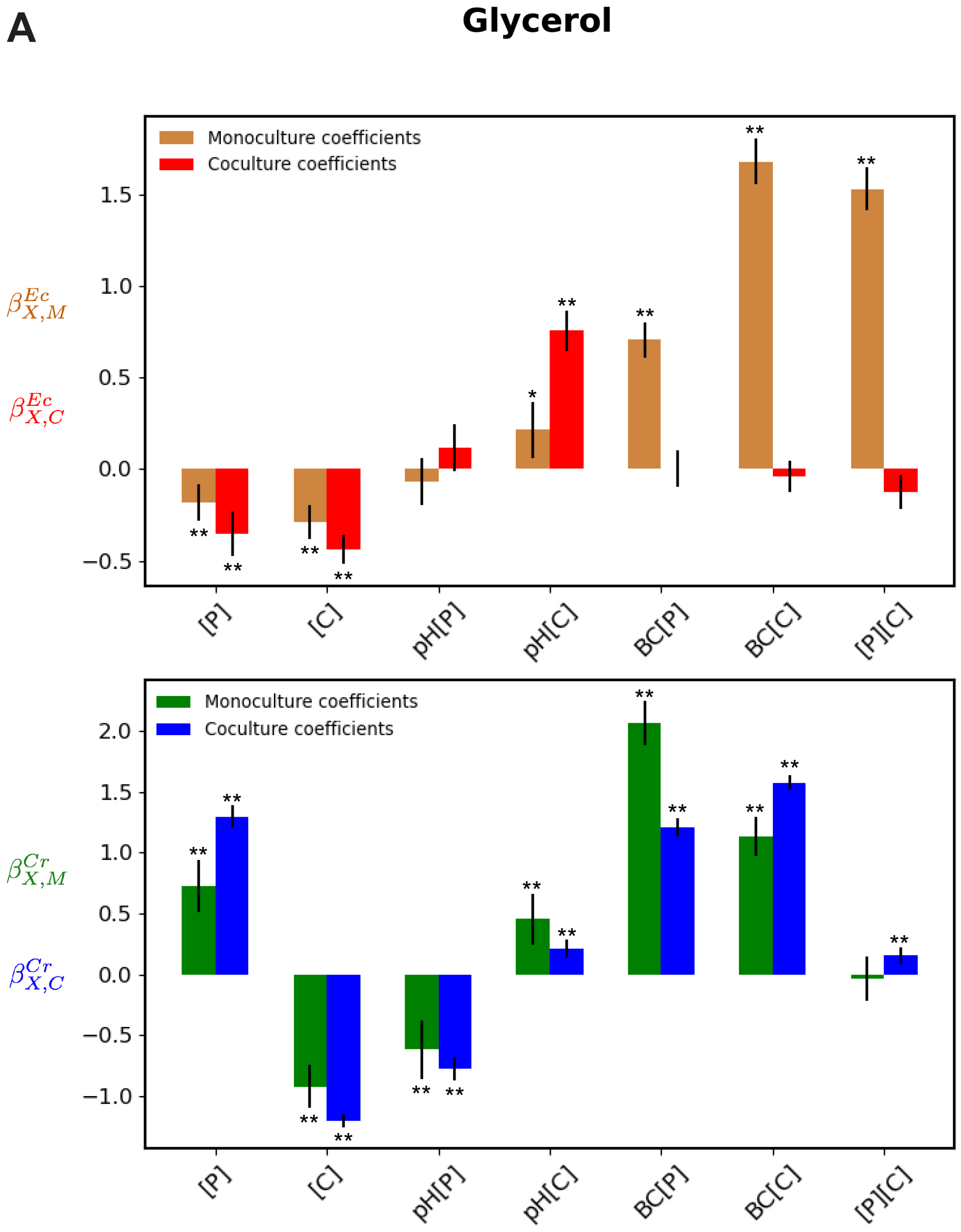
A
Glycerol
Monoculture coefficients
Coculture coefficients
$\beta^{Ec}_{X,M}$
$\beta^{Ec}_{X,C}$
[P]
[C]
pH[P]
pH[C]
BC[P]
BC[C]
[P][C]
$\beta^{Cr}_{X,M}$
$\beta^{Cr}_{X,C}$

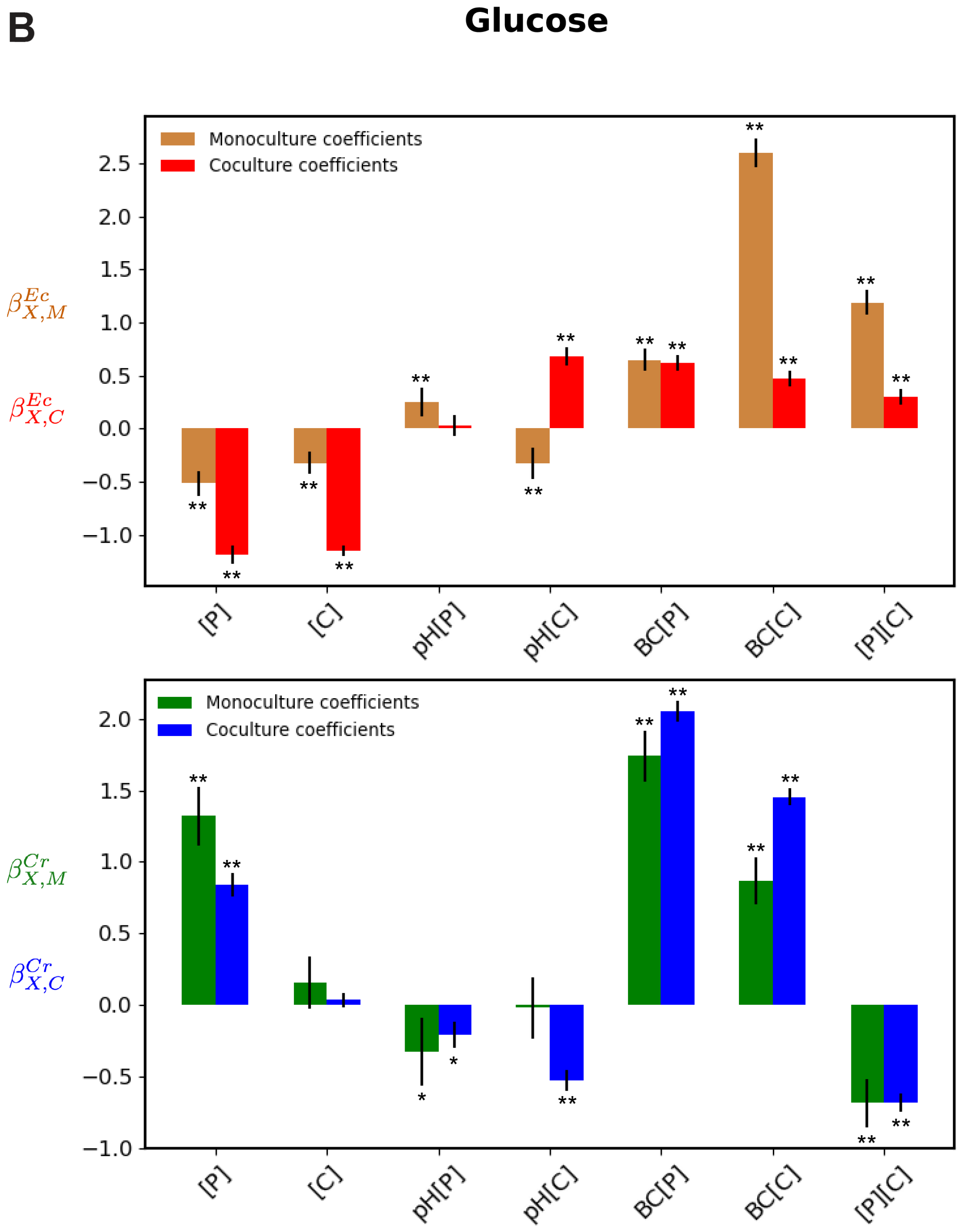
B
Glucose
Monoculture coefficients
Coculture coefficients
$\beta^{Ec}_{X,M}$
$\beta^{Ec}_{X,C}$
[P]
[C]
pH[P]
pH[C]
BC[P]
BC[C]
[P][C]
Monoculture coefficients
Coculture coefficients
$\beta^{Cr}_{X,M}$
$\beta^{Cr}_{X,C}$

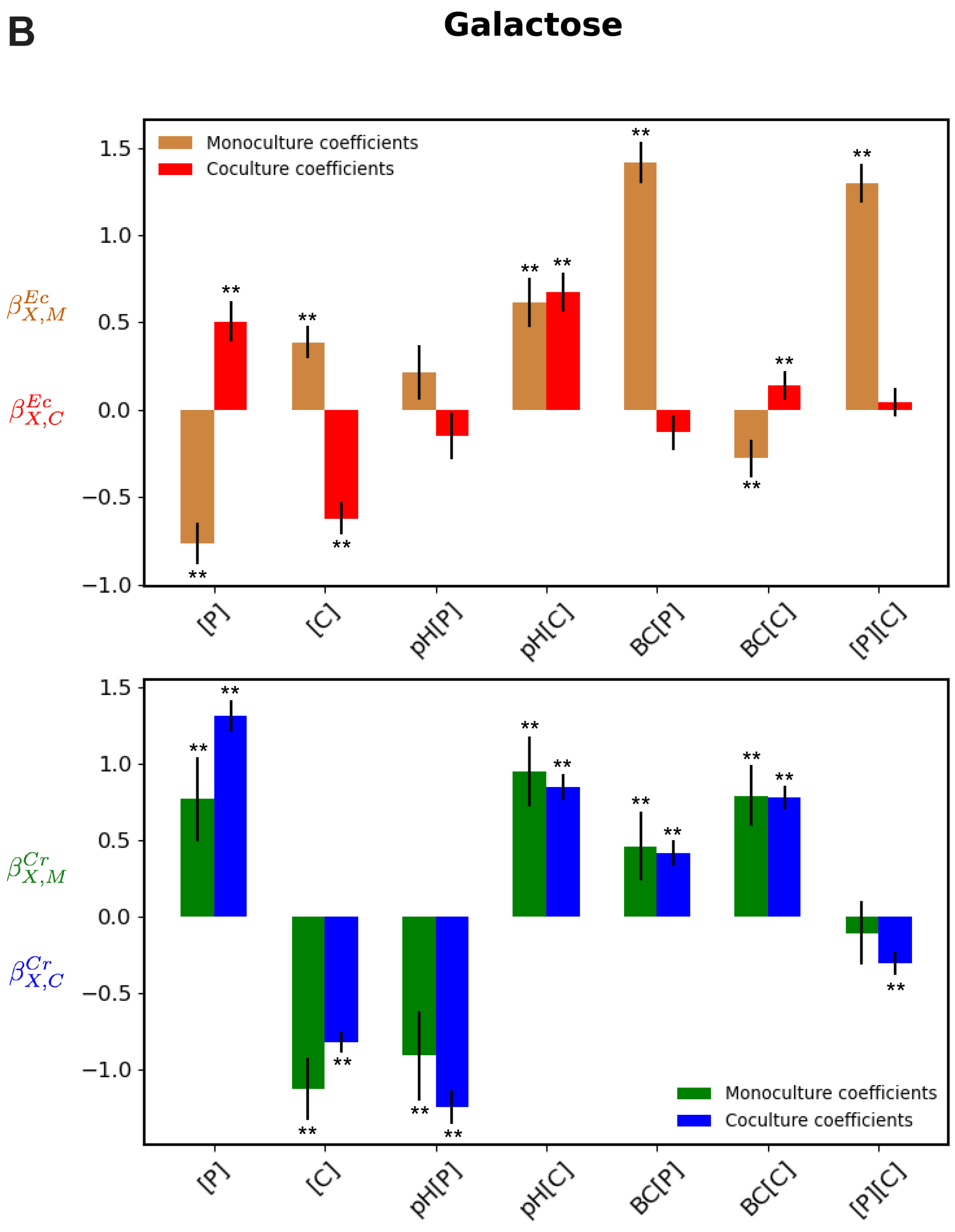
B
Galactose
Monoculture coefficients
Coculture coefficients
$\beta^{Ec}_{X,M}$
$\beta^{Ec}_{X,C}$
$\beta^{Cr}_{X,M}$
$\beta^{Cr}_{X,C}$
1.5
1.0
0.5
0.0
−0.5
−1.0
[P]
[C]
pH[P]
pH[C]
BC[P]
BC[C]
[P][C]
**
Monoculture coefficients
Coculture coefficients

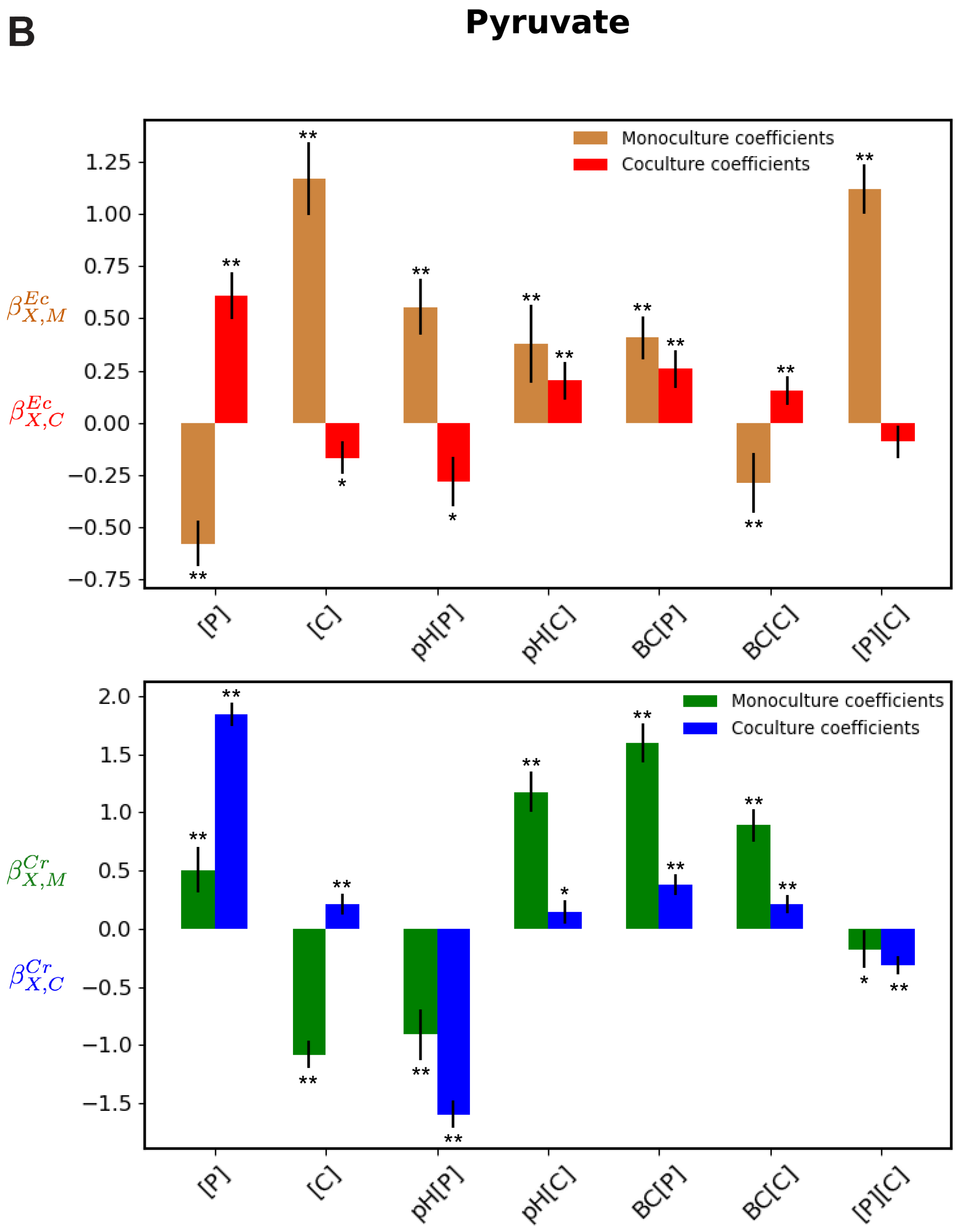
B
Pyruvate
Monoculture coefficients
Coculture coefficients
$\beta^{Ec}_{X,M}$
$\beta^{Ec}_{X,C}$
[P]
[C]
pH[P]
pH[C]
BC[P]
BC[C]
[P][C]
Monoculture coefficients
Coculture coefficients
$\beta^{Cr}_{X,M}$
$\beta^{Cr}_{X,C}$
[P]
[C]
pH[P]
pH[C]
BC[P]
BC[C]
[P][C]

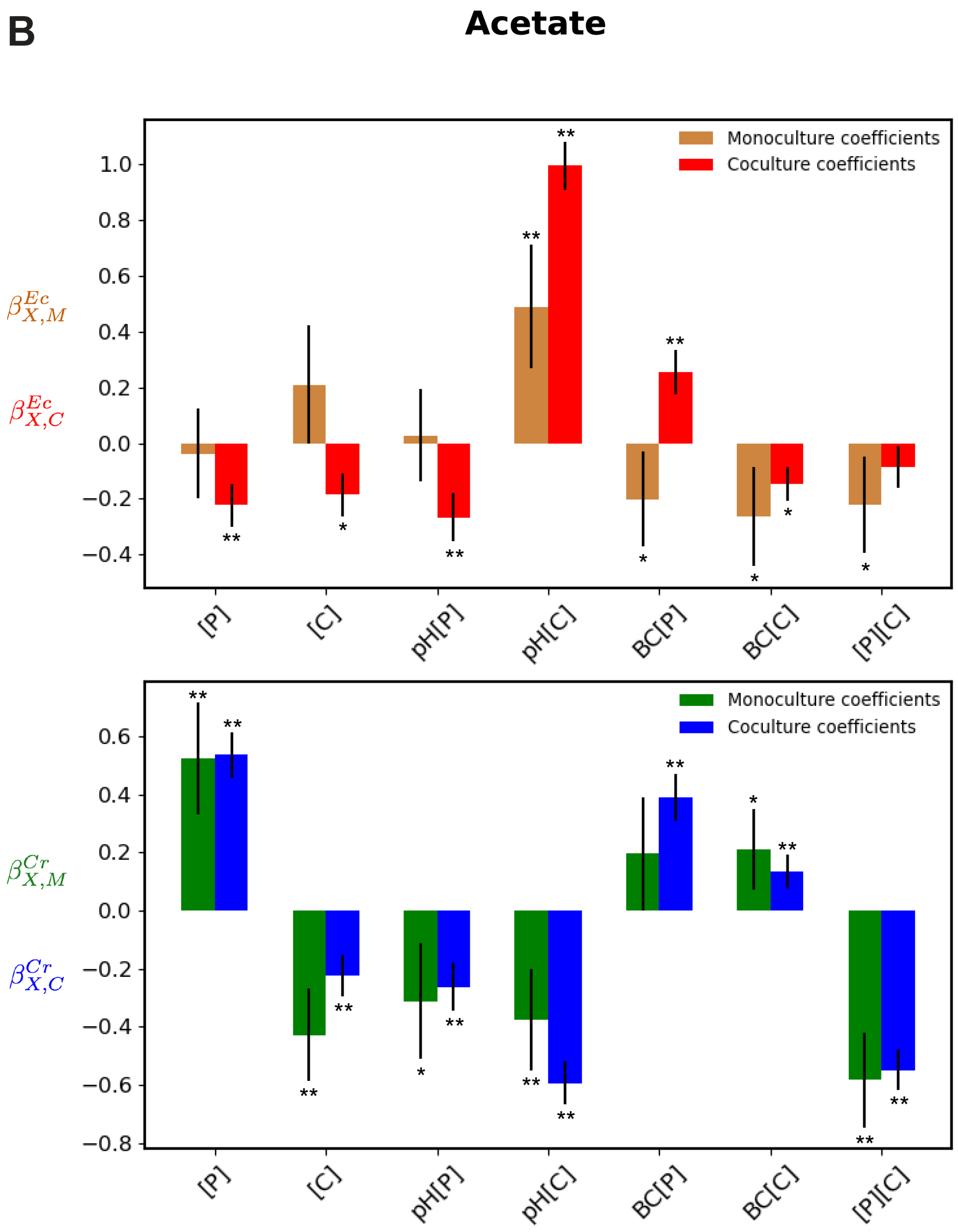

**Figure S11:** (Caption next page)

**Figure S11:** (Previous page) **Comparison between the monoculture and coculture coefficients obtained from regressing algae-bacteria growth** The top panels in (A),(B),(C),(D), and (E) report the monoculture coefficients (brown bars) and the coculture coefficients (red bars) of the corresponding features on the x-axis obtained from the regression model (Equation. S26) predicting the growth of *E. coli* in monocultures and cocultures in the case of glucose in (A), galactose in (B), pyruvate in (C), and acetate in (D). Similarly, the bottom panel reports the monoculture coefficients (green bars) and the coculture coefficients (blue bars) of the corresponding features on the x-axis obtained from the regression model predicting the growth of *C. reinhardtii* in monocultures and cocultures in the case of glucose in (A), galactose in (B), pyruvate in (C) and acetate in (D). The error bars represent the 95% confidence intervals. The statistical significance of the results is indicated with **(p-value<0.001) and *(p-value<0.05) atop the bars. The coculture coefficients, their confidence intervals, and p-values were computed as described in section 8.4

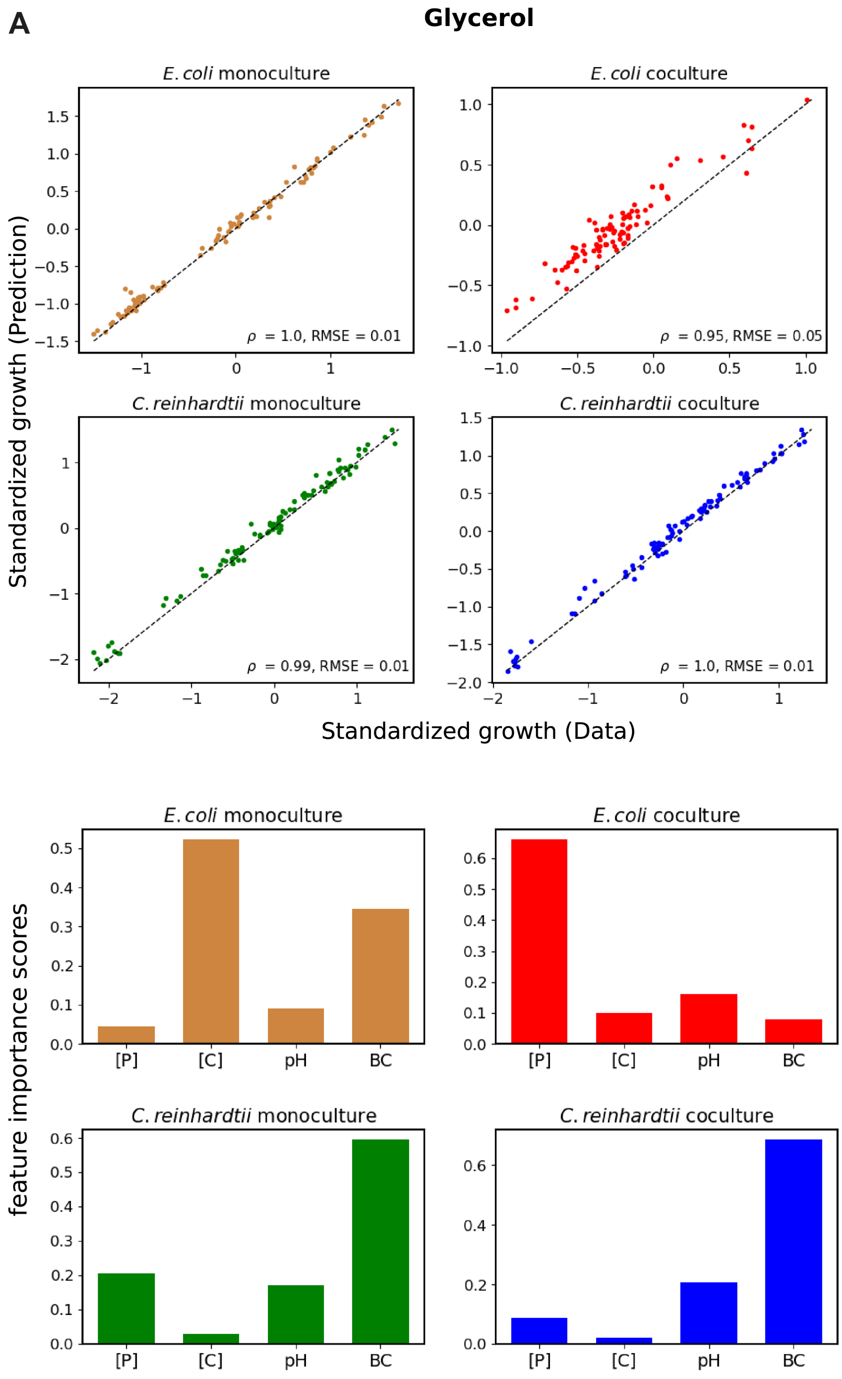
A
Glycerol
E. coli monoculture
E. coli coculture
C. reinhardtii monoculture
C. reinhardtii coculture
ρ = 1.0, RMSE = 0.01
ρ = 0.95, RMSE = 0.05
ρ = 0.99, RMSE = 0.01
ρ = 1.0, RMSE = 0.01
Standardized growth (Prediction)
Standardized growth (Data)
feature importance scores
[P]
[C]
pH
BC

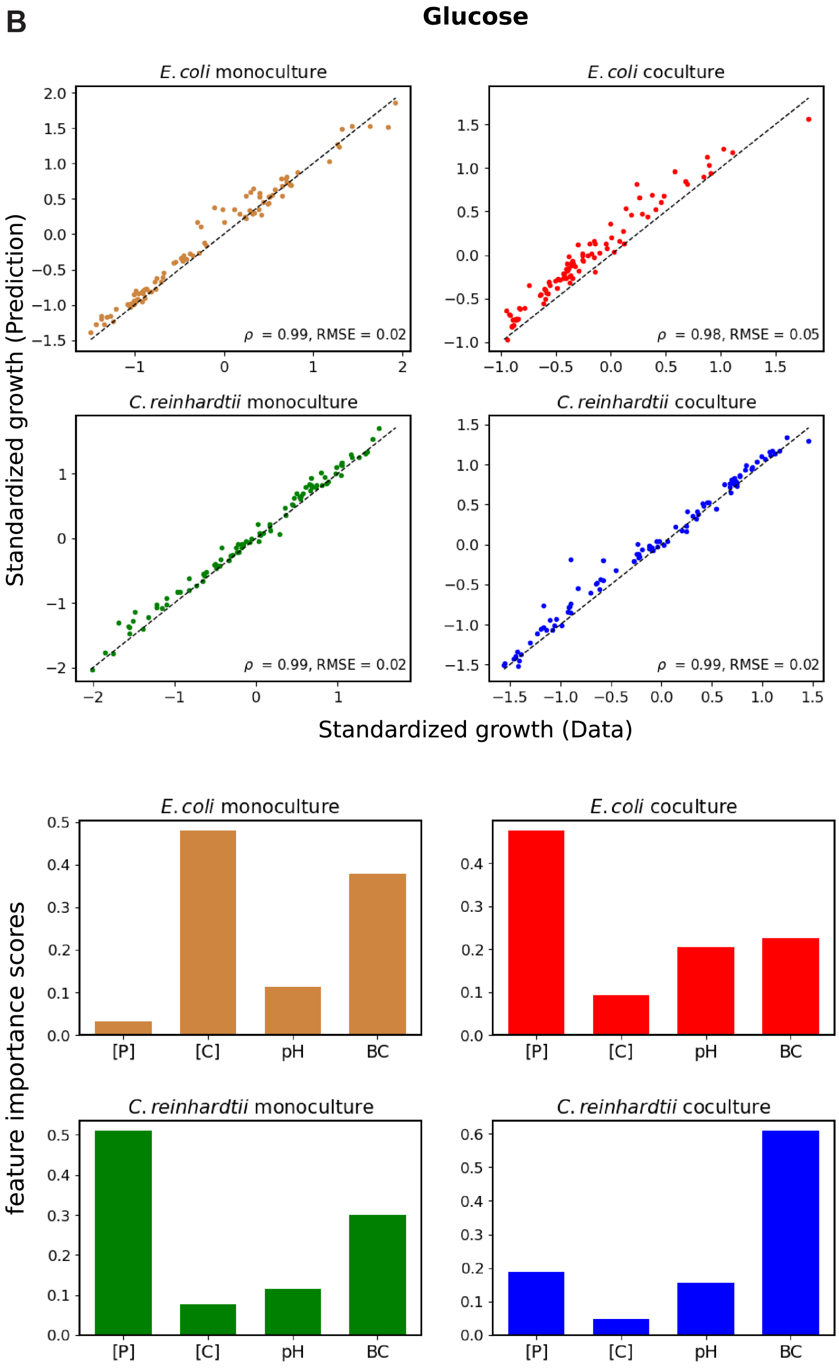
B
Glucose
E. coli monoculture
E. coli coculture
C. reinhardtii monoculture
C. reinhardtii coculture
ρ = 0.99, RMSE = 0.02
ρ = 0.98, RMSE = 0.05
ρ = 0.99, RMSE = 0.02
ρ = 0.99, RMSE = 0.02
Standardized growth (Prediction)
Standardized growth (Data)
feature importance scores
[P]
[C]
pH
BC

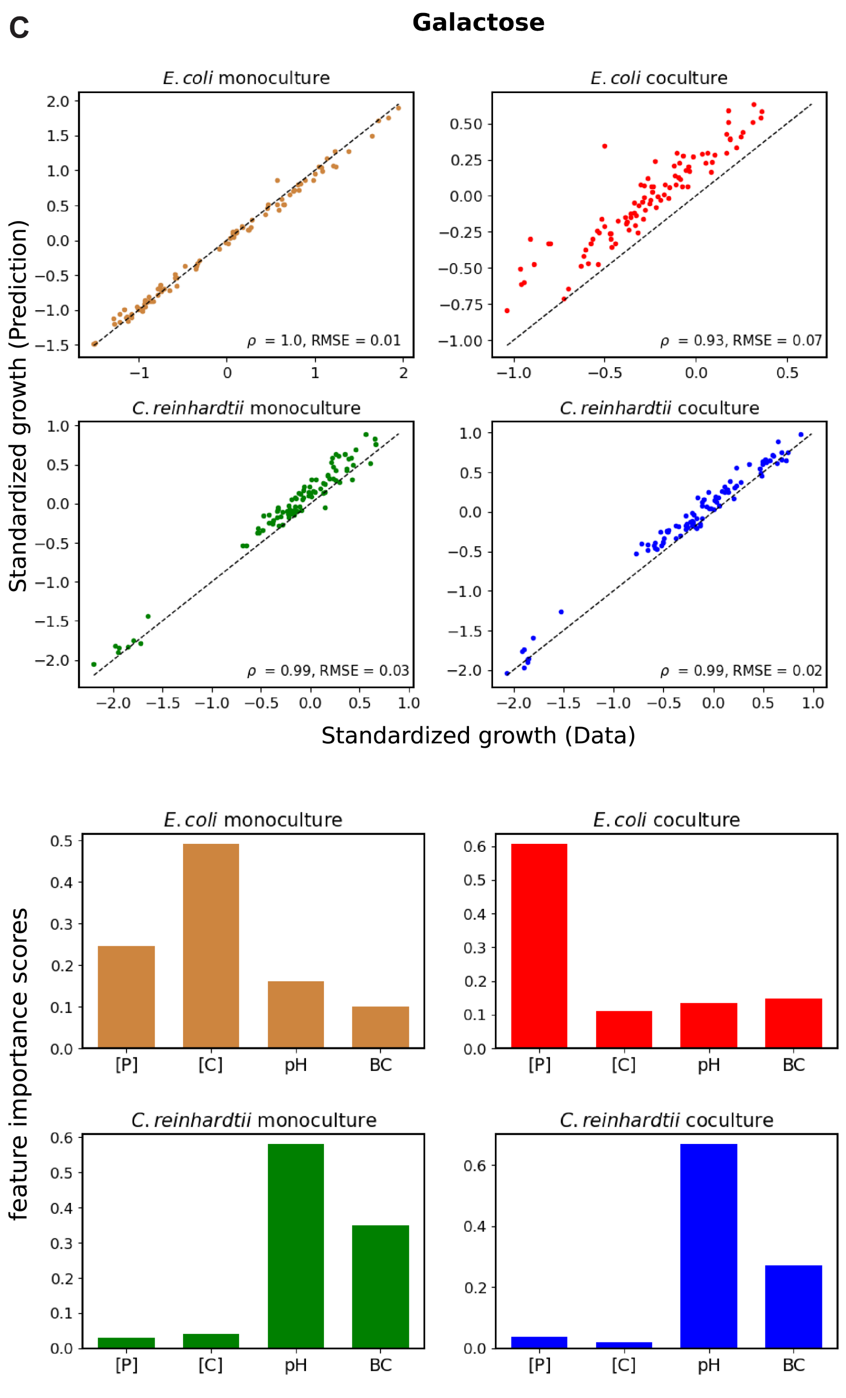
C
Galactose
E. coli monoculture
E. coli coculture
C. reinhardtii monoculture
C. reinhardtii coculture
ρ = 1.0, RMSE = 0.01
ρ = 0.93, RMSE = 0.07
ρ = 0.99, RMSE = 0.03
ρ = 0.99, RMSE = 0.02
Standardized growth (Prediction)
Standardized growth (Data)
E. coli monoculture
E. coli coculture
C. reinhardtii monoculture
C. reinhardtii coculture
feature importance scores
[P]
[C]
pH
BC

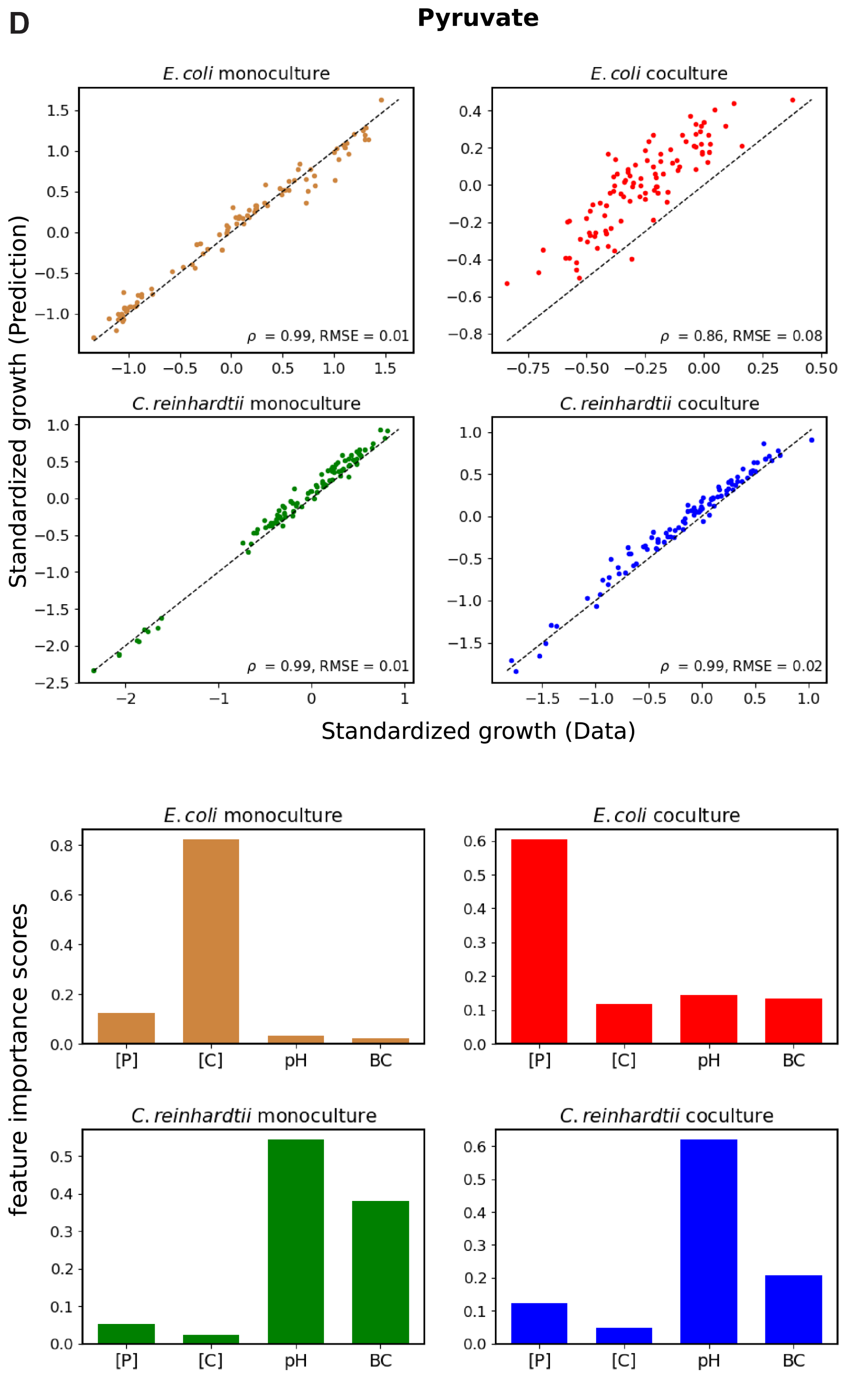
D
Pyruvate
E. coli monoculture
E. coli coculture
C. reinhardtii monoculture
C. reinhardtii coculture
ρ = 0.99, RMSE = 0.01
ρ = 0.86, RMSE = 0.08
ρ = 0.99, RMSE = 0.01
ρ = 0.99, RMSE = 0.02
Standardized growth (Prediction)
Standardized growth (Data)
E. coli monoculture
E. coli coculture
C. reinhardtii monoculture
C. reinhardtii coculture
feature importance scores
[P]
[C]
pH
BC

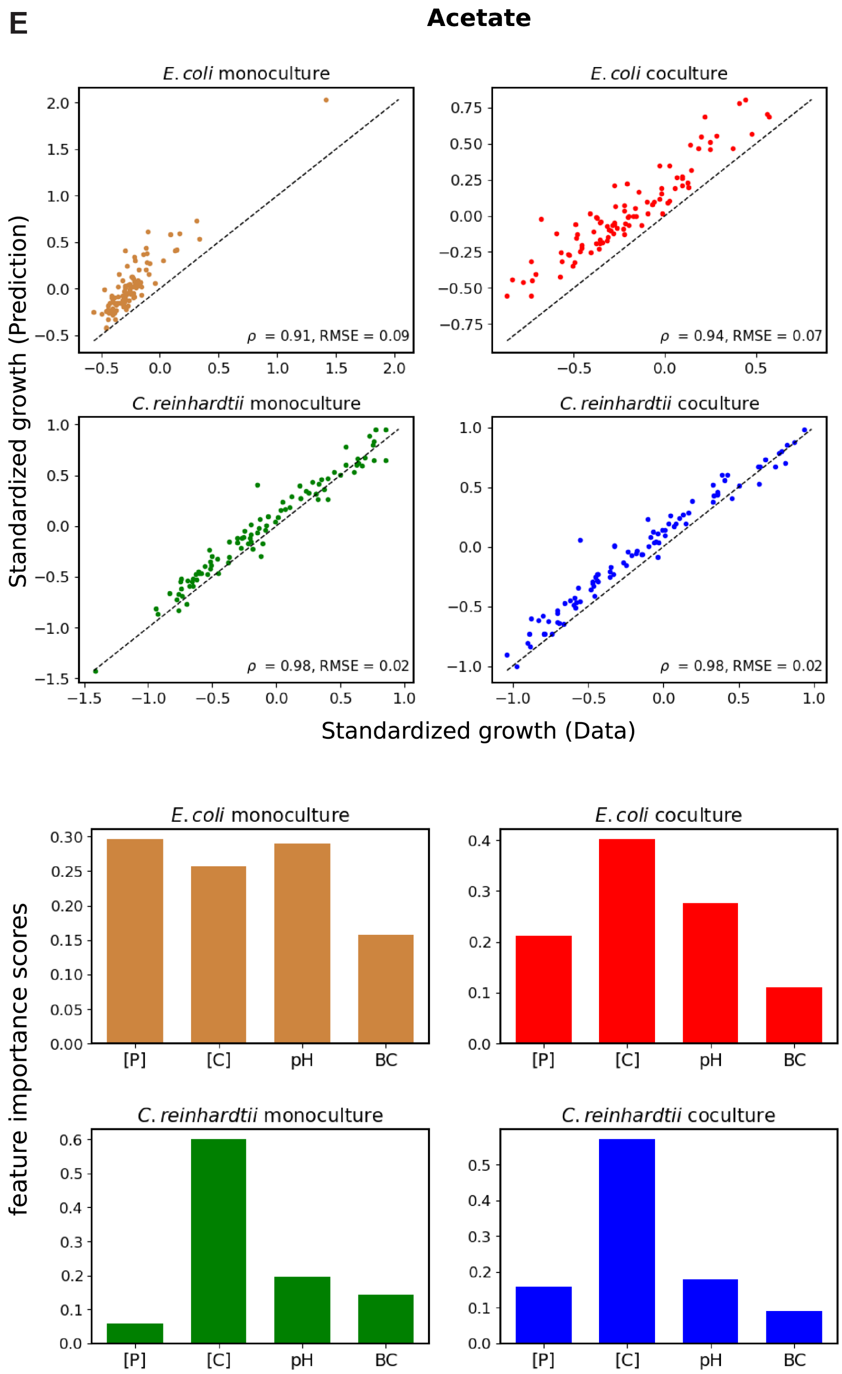

**Figure S12:** (Caption next page)

**Figure S12:** (Previous page) **Improved predictions of the algae-bacteria growth in the different environmental conditions using random forest models.** Random forest model was implemented in python using scikit-learn (RandomForestRegressor) with just pH, buffering capacity BC, phosphorus concentration [P], and carbon concentration [C] as the features, with the number of decision trees set to 100 and using default values for all other parameters. 4 such random forest models were set up, one each to predict the growth of *E. coli* in monoculture, *E. coli* in coculture, *C. reinhardtii* in monoculture, *C. reinhardtii* in coculture. The top set of 4 panels in (A),(B),(C),(D), and (E) show the comparison between the predicted growth from the random forest models and the observed growth of *E. coli* and *C. reinhardtii* in the environments on kChip in the case of glycerol in (A), glucose in (B), galactose in (C), pyruvate in (D), and acetate in (E). The observed growth was computed as the median of the standardized growth across replicate wells. The predicted growth was computed as the median of the standardized growth obtained from the random forest model across replicates. See section 8.2 for details about the standardization of the growth data. The comparison between the predicted growth (y-axis) and the observed growth (x-axis) for all the environments is shown for *E. coli* in monoculture in the top left, *E. coli* in coculture in the top right, *C. reinhardtii* in monoculture in the bottom left, and *C. reinhardtii* in coculture in the bottom right. The Pearson correlation coefficient $\rho$ and the root mean squared error RMSE of the fits are reported in each panel. The bottom set of 4 panels in (A),(B),(C),(D), and (E) report the feature importance scores of the corresponding features on the x-axis obtained from the 4 random forest models predicting the growth of *E. coli* in monoculture in the top left panel, *E. coli* in coculture in the top right panel, *C. reinhardtii* in monoculture in the bottom left panel, and *C. reinhardtii* in coculture in the bottom right panel, in the case of glycerol in (A), glucose in (B), galactose in (C), pyruvate in (D), and acetate in (E).

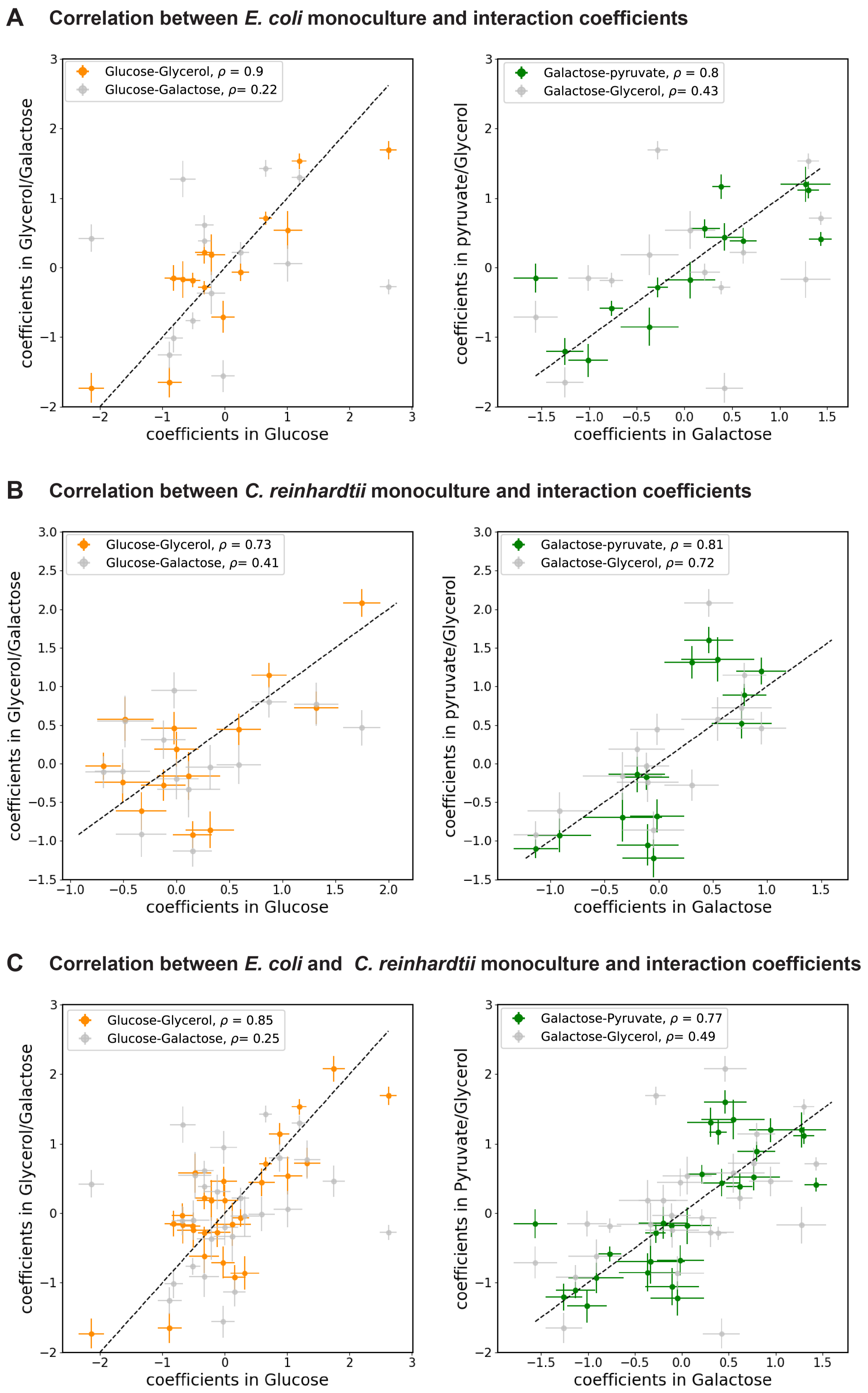

**Figure S13:** (Caption next page)

**Figure S13:** (Previous page) **Correlations between monoculture and interaction coefficients obtained from regressing the algae-bacteria growth in the different carbon sources.** (A) In the left panel, the monoculture and interaction coefficients of the features $[P]$,$[C]$,$pH[C]$,$pH[P]$,$BC[P]$,$BC[C]$,$[P][C]$ obtained from regressing *E. coli* growth in glycerol (orange points) and galactose (light gray points) on the y-axis are plotted against the corresponding coefficients in glucose on the x-axis. In the right panel, the monoculture and interaction coefficients of the features $[P]$,$[C]$,$pH[C]$,$pH[P]$,$BC[P]$,$BC[C]$,$[P][C]$ obtained from regressing *E. coli* growth in pyruvate (green points) and glycerol (light gray points) on the y-axis are plotted against the corresponding coefficients in galactose on the x-axis. (B) shows the same as in (A) for monoculture and interaction coefficients of the features $[P]$,$[C]$,$pH[C]$,$pH[P]$,$BC[P]$,$BC[C]$,$[P][C]$ obtained from regressing *C. reinhardtii* growth (C) shows the same as in (A) for monoculture and interaction coefficients of the features $[P]$,$[C]$,$pH[C]$,$pH[P]$,$BC[P]$,$BC[C]$,$[P][C]$ obtained from regressing both *E. coli* and *C. reinhardtii* growth

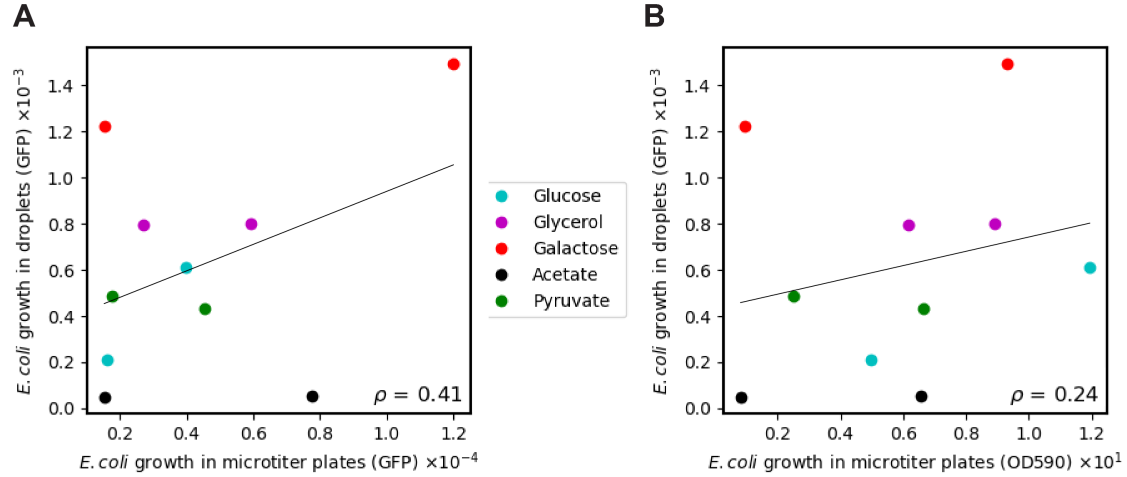

**Figure S14: Comparison between *E. coli* growth in droplets and microtiter plates.** (A) Median monoculture growth of *E. coli* in the environments E3+E4 and E11+E12 (Table S2) in droplets (y-axis) is plotted against the median monoculture growth of *E. coli* in the corresponding environments in microtiter plates as measured by the GFP fluorescence using the Tecan infinite F200 PRO plate reader (x-axis), for all the carbon sources (legend). (B) Median monoculture growth of *E. coli* in the environments E3+E4 and E11+E12 (Table S2) in droplets (y-axis) is plotted against the median monoculture growth of *E. coli* in the corresponding environments in microtiter plates as measured by OD590 using the Tecan infinite F200 PRO plate reader (x-axis), for all the carbon sources (legend).
In both (A) and (B), the Pearson correlation coefficient $\rho$ between the data is reported within the panels and the thin black line is the fit from simple linear regression predicting the median *E. coli* growth in droplets from either the median *E. coli* growth in microtiter plates as inferred by GFP or OD590.

| Compound | Concentration |
|---|---|
| $MgSO_4 \cdot 7H_2O$ | 0.2 mM |
| $CaCl_2$ | 2 mM |
| $C_{10}H_{16}N_2O_8$ (EDTA) | 11 $\mu$M |
| $FeSO_4 \cdot 7H_2O$ | 11 $\mu$M |
| $H_3BO_4$ | 30 $\mu$M |
| $ZnSO_4 \cdot 7H_2O$ | 1 $\mu$M |
| $MnCl_2 \cdot 4H_2O$ | 7 $\mu$M |
| $Na_2MoO_4 \cdot 2H_2O$ | 1 $\mu$M |
| $CuSO_4 \cdot 5H_2O$ | 0.2 $\mu$M |
| $Co(NO_3)_2 \cdot 6H_2O$ | 1 $\mu$M |
| NaOH | 33 $\mu$M |
| NaCl | 3 mM |

**Table S1:** Modified 1x Taub medium composition

| Environment | pH | BC (mM) | [P] (mM) | [C] (mM) | [A555] ($\mu$M) | [A595] ($\mu$M) | [A647] ($\mu$M) |
|---|---|---|---|---|---|---|---|
| E1 | 6.14 | 0.001 | 0.03 | 2 | 1 | 0 | 0 |
| E2 | 6.80 | 1.705 | 4 | 2 | 0 | 1 | 0 |
| E3 | 6.14 | 0.001 | 0.03 | 10 | 0 | 0 | 1 |
| E4 | 6.80 | 1.705 | 4 | 10 | 0.8 | 0.2 | 0 |
| E5 | 6.47 | 0.002 | 0.03 | 2 | 0.7 | 0 | 0.3 |
| E6 | 7.38 | 1.73 | 3 | 2 | 0.3 | 0 | 0.7 |
| E7 | 6.47 | 0.002 | 0.03 | 10 | 0.6 | 0.4 | 0 |
| E8 | 7.38 | 1.73 | 3 | 10 | 0.5 | 0 | 0.5 |
| E9 | 6.93 | 3.396 | 0.01 | 2 | 0 | 0.8 | 0.2 |
| E10 | 6.94 | 3.436 | 0.08 | 2 | 0 | 0.4 | 0.6 |
| E11 | 6.93 | 3.396 | 0.01 | 10 | 0 | 0.6 | 0.4 |
| E12 | 6.94 | 3.436 | 0.08 | 10 | 0.1 | 0.8 | 0.1 |
| E13 | 7.40 | 3.392 | 0.01 | 2 | 0.9 | 0 | 0.1 |
| E14 | 7.40 | 3.426 | 0.08 | 2 | 0 | 0.2 | 0.8 |
| E15 | 7.40 | 3.392 | 0.01 | 10 | 0.4 | 0.3 | 0.3 |
| E16 | 7.40 | 3.426 | 0.08 | 10 | 0.4 | 0.6 | 0 |

**Table S2:** Initial pH (pH), buffering capacity (BC), phosphorus concentration ([P]), carbon concentration ([C]), Alexa Fluor 555 dye concentration ([A555]), Alexa Fluor 594 dye concentration ([A594]), and Alexa Fluor 647 dye concentration ([A647]) in the 16 barcoded environmental conditions E1-E16.

| Environment | Base media | | Carbon | | Nitrogen | | Phosphorus | | Alexa 555 | | Alexa 594 | | Alexa 647 | | Water ($\mu$L) |
|---|---|---|---|---|---|---|---|---|---|---|---|---|---|---|---|
| | Stock | vol ($\mu$L) | Stock | vol ($\mu$L) | Stock | vol ($\mu$L) | Stock | vol ($\mu$L) | Stock | vol ($\mu$L) | Stock | vol ($\mu$L) | Stock | vol ($\mu$L) | |
| E1 | B1 | 230 | C2 | 15 | N1 | 10 | P2 | 15 | D1 | 0 | D2 | 16 | D3 | 4 | 190 |
| E2 | B1 | 230 | C2 | 15 | N1 | 10 | P1 | 40 | D1 | 0 | D2 | 8 | D3 | 12 | 165 |
| E3 | B1 | 230 | C1 | 12 | N1 | 10 | P2 | 15 | D1 | 20 | D2 | 0 | D3 | 0 | 193 |
| E4 | B1 | 230 | C1 | 12 | N1 | 10 | P1 | 40 | D1 | 0 | D2 | 20 | D3 | 0 | 168 |
| E5 | B1 | 230 | C2 | 15 | N1 | 10 | P3 | 125 | D1 | 0 | D2 | 12 | D3 | 8 | 80 |
| E6 | B1 | 230 | C2 | 15 | N1 | 10 | P4 | 125 | D1 | 2 | D2 | 16 | D3 | 2 | 80 |
| E7 | B1 | 230 | C1 | 12 | N1 | 10 | P3 | 125 | D1 | 0 | D2 | 0 | D3 | 20 | 83 |
| E8 | B1 | 230 | C1 | 12 | N1 | 10 | P4 | 125 | D1 | 16 | D2 | 4 | D3 | 0 | 83 |
| E9 | B2 | 230 | C2 | 15 | N1 | 10 | P2 | 5 | D1 | 18 | D2 | 0 | D3 | 2 | 200 |
| E10 | B2 | 230 | C2 | 15 | N1 | 10 | P2 | 40 | D1 | 0 | D2 | 4 | D3 | 16 | 165 |
| E11 | B2 | 230 | C1 | 12 | N1 | 10 | P2 | 5 | D1 | 14 | D2 | 0 | D3 | 6 | 203 |
| E12 | B2 | 230 | C1 | 12 | N1 | 10 | P2 | 40 | D1 | 6 | D2 | 0 | D3 | 14 | 168 |
| E13 | B3 | 230 | C2 | 15 | N1 | 10 | P2 | 5 | D1 | 8 | D2 | 6 | D3 | 6 | 200 |
| E14 | B3 | 230 | C2 | 15 | N1 | 10 | P2 | 40 | D1 | 8 | D2 | 12 | D3 | 0 | 165 |
| E15 | B3 | 230 | C1 | 12 | N1 | 10 | P2 | 5 | D1 | 12 | D2 | 8 | D3 | 0 | 203 |
| E16 | B3 | 230 | C1 | 12 | N1 | 10 | P2 | 40 | D1 | 10 | D2 | 0 | D3 | 10 | 168 |

**Table S3:** Preparing the initial barcoded environments E1-E16. The volumes of the different stock solutions B1,B2,B3,C1,C2,N1,P1,P2,P3,P4,D1,D2,D3, and water used to make each of the environments E1-E16 are reported. See section 3.2 for details about the stock solutions.

| Buffering agent | $pK_a$ (25C) | $\Delta H(kJ)$ | $C_p$ | $pK_a$ (30C) |
|---|---|---|---|---|
| $H_2PO_4^-$ | 7.198 | 3.6 | -230 | 7.189 |
| Tris | 8.072 | 47.25 | -59 | 7.936 |
| MOPS | 7.184 | 21.10 | 25 | 7.123 |
| $NH_4^+$ | 9.245 | 51.95 | 8 | 9.095 |
| Acetate | 4.756 | -0.41 | -142 | 4.758 |
| Pyruvate | 2.93 | NA | NA | NA |

**Table S4:** $pK_a$ of buffering agents in the medium. For a buffering agent $X$, the equilibrium constant $K$ for the reaction $HX \leftrightarrow H^+ + X^-$ is given by $K = 10^{-pK_a}$. The equilibrium constant depends on temperature and $pK_a$ value at experimental temperature ($30°C$) is calculated using formula in section 7. All chemical values are collected from [12], except for pyruvate, which we collected the standard $pK_a$ value from [16]. $\Delta H$ and $\Delta C_p$ values for pyruvate are not found so we used the $pK_a$ value at $25°C$ as an approximate.

| Carbon source | Environment | Initial pH | Buffering capacity (mM) | Growth rate ($hr^{-1}$) | Growth (OD590) | Final pH |
|---|---|---|---|---|---|---|
| Glycerol | E3+E4 | 6.78 | 0.74 | 0.161 | 0.063 | ~5 |
| | E11+E12 | 6.93 | 3.42 | 0.192 | 0.091 | ~6 |
| Glucose | E3+E4 | 6.78 | 0.74 | 0.246 | 0.052 | ~5 |
| | E11+E12 | 6.93 | 3.42 | 0.276 | 0.121 | ~6 |
| Galactose | E3+E4 | 6.78 | 0.74 | 0.073 | 0.014 | ~6 |
| | E11+E12 | 6.93 | 3.42 | 0.086 | 0.097 | $\tilde{6}$ |
| Pyruvate | E3+E4 | 6.78 | 0.74 | 0.137 | 0.028 | ~7 |
| | E11+E12 | 6.93 | 3.45 | 0.18 | 0.07 | ~7 |
| Acetate | E3+E4 | 6.82 | 0.74 | 0.017 | 0.011 | ~7 |
| | E11+E12 | 6.94 | 3.42 | 0.081 | 0.069 | ~7 |

**Table S5:** Comparison of the growth of *E. coli* in the environments E3+E4 and E11+E12 across the different carbon sources. The initial pH and buffering capacities of the environments (predicted by the titration model described in section 7), the growth rates and OD590 (at 68 h) of *E. coli*, and the final pH (at 68 h) of the cultures (measured using VWR pH paper BDH35309.606) in microtiter plates are reported for the environments E3+E4 and E11+E12 in the different carbon sources. The reported growth rates were computed as described in section 10

| Carbon source | *E. coli* monoculture growth | | *E. coli* coculture growth | | *C. reinhardtii* monoculture growth | | *C. reinhardtii* coculture growth | |
|---|---|---|---|---|---|---|---|---|
| | median | standard deviation | median | standard deviation | median | standard deviation | median | standard deviation |
| Glycerol | 751.77 | 411.05 | 54.7 | 61.17 | 10.55 | 7.1 | 9.13 | 6.16 |
| Glucose | 270.31 | 165.95 | 21.16 | 42.58 | 9.25 | 9.01 | 7.59 | 6.97 |
| Galactose | 723.18 | 379.01 | 44.79 | 42.89 | 10.79 | 8.05 | 9.49 | 6.25 |
| Pyruvate | 279.48 | 179.73 | 26.12 | 40.56 | 8.01 | 4.37 | 8.16 | 3.56 |
| Acetate | 46.64 | 70.02 | 14.26 | 20.97 | 14.63 | 6.9 | 13.07 | 6.01 |

**Table S6:** Median and standard deviation of the growth of bacteria and algae in monoculture and coculture across all the ($\sim$ 105) environments formed on kChip for the different carbon sources.
